# Temporal tissue dynamics from a spatial snapshot

**DOI:** 10.1101/2024.04.22.590503

**Authors:** Jonathan Somer, Shie Mannor, Uri Alon

## Abstract

Physiological and pathological processes such as inflammation or cancer emerge from the interactions between cells over time. However, methods to follow cell populations over time within the native context of a human tissue are lacking, because a biopsy offers only a single snapshot. Here we present one-shot tissue dynamics reconstruction (OSDR), an approach to estimate a dynamical model of cell populations based on a single tissue sample. OSDR uses spatial proteomics to learn how the composition of cellular neighborhoods influences division rate, providing a dynamical model of cell population change over time. We apply OSDR to human breast cancer data, and reconstruct two fixed points of fibroblasts and macrophage interactions. These fixed points correspond to hot and cold fibrosis, in agreement with co-culture experiments that measured these dynamics directly. We then use OSDR to discover a pulse-generating excitable circuit of T and B cells in the tumor microenvironment, suggesting temporal flares of adaptive anti-cancer responses. Finally, we study longitudinal biopsies from a triple negative breast cancer clinical trial, where OSDR predicts the collapse of the tumor cell population in responders versus non-responders based on early treatment biopsies. OSDR can be applied to a wide range of spatial proteomics assays to enable analysis of tissue dynamics based on patient biopsies.

## Introduction

Physiological processes involve cell populations that change over time (Adler, Chavan, and Medzhitov 2023). Some cell populations expand and others are removed, as occurs in development and in the immune response to pathogens (Medzhitov 2021). A clinically important example of changing cell populations is the evolution of the cancer microenvironment, in which the growing tumor recruits stromal and immune cells that are crucial for tumor survival (Sahai et al. 2020; Mayer et al. 2023; Gascard and Tlsty 2016; Tlsty and Coussens 2006; Schürch et al. 2020; Binnewies et al. 2018; Combes, Samad, and Krummel 2023). Understanding cell population dynamics and the underlying cell-cell communication circuits is a major goal of tissue biology. Understanding these circuits can enable new treatment strategies based on sculpting cell populations in desired ways (S. Wang et al. 2023; Miyara et al. 2023)

However, measuring the dynamics of cell populations in human tissues is currently very difficult. Biopsies provide a single snapshot, and taking multiple biopsies from a single patient is infeasible and does not provide longitudinal evidence from the same cells.

Current strategies to measure tissue dynamics do not apply to human biopsies. For example cell lines, mice, organ-on-a-chip (Leung et al. 2022) or organoid models (Kim, Koo, and Knoblich 2020) are treated as replicas and are analyzed at different time points. Ex-vivo tissues can be followed over time but lack the native physiological context (Kokkinos et al. 2021). Intravital fluorescence microscopy has made it possible to follow living cells within animal models (Pittet et al. 2018; Entenberg, Oktay, and Condeelis 2023). Recent advances employ synthetic biology to engineer cells to record their activity or lineage ((Quinn et al. 2021), *Horns et al. 2023*). These approaches are restricted to animal models or in vitro-settings, limiting their ability to capture the complexity of *in vivo* dynamics in humans.

The emergence of single-cell technologies offers new opportunities for studying tissues at high resolution. Notable approaches use single-cell data for understanding cell-cell communication at a single time point (Efremova et al. 2020; Jerby-Arnon and Regev 2022; Browaeys, Saelens, and Saeys 2020; Armingol, Baghdassarian, and Lewis 2024). Other approaches attempt to infer dynamics of processes such as transcription within individual cells on a timescale of hours. Examples include RNA velocity, ergodic rate analysis and Zman-seq (La Manno et al. 2018; Kafri et al. 2013; Kirschenbaum et al. 2023). These methods do not address the challenge of understanding how cell populations change on the tissue level, processes that could take days to weeks.

Here we present an approach to estimate cell population dynamics based on a spatial biopsy snapshot (Figure 1A). This approach, ‘One-Shot Tissue Dynamics Reconstruction’ (OSDR), is based on using a cell-division marker to determine division rates as a function of neighborhood composition, providing a dynamical model of how cell populations change over time. We apply OSDR to human breast-cancer spatial proteomics samples from three large cohorts (Danenberg et al. 2022; X. Q. Wang et al. 2023; Fischer et al. 2023). OSDR reconstructs a fibroblast-macrophage circuit with two steady states of hot and cold fibrosis (Adler et al. 2020), in agreement with co-culture experiments that measure dynamics directly (Mayer et al. 2023). We then use OSDR to discover a pulse-generating excitable circuit of T cells and B cells, suggesting that cancer surveillance in the tumor microenvironment operates in temporal flares, as opposed to the steady-state picture implicit in current literature. Finally, we validate OSDR using longitudinal data from patients who received either chemotherapy or chemotherapy and immunotherapy. In both treatment regimes OSDR predicts the collapse of the tumor cell population in responders but not in non-responders, based on biopsies at treatment initiation. The present approach opens the way to infer cell population dynamics from spatial snapshots of patient biopsies.

**Figure 1:**
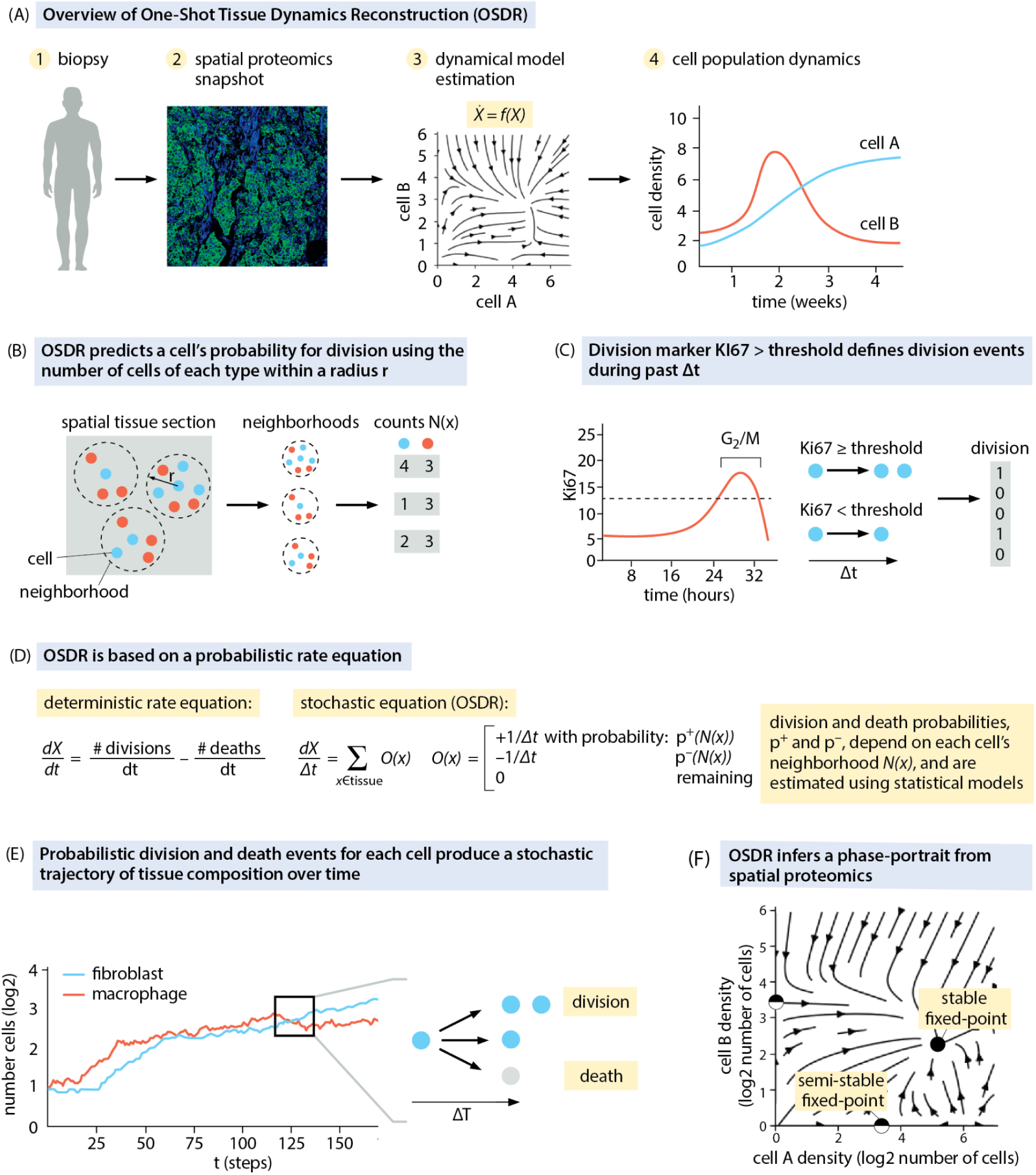
Overview of One-Shot Tissue Dynamics Reconstruction (OSDR). A) OSDR uses spatial proteomics of a biopsy to infer cell population dynamics. B) The division probability is learned based on the numbers of cells of each type in the neighborhood. C) Cell division rates are inferred from a cell division marker. D) Rate of change in the number of cells *X* is the difference between divisions and removals as described by a stochastic model E) Simulations can take an initial spatial arrangement of cells and propagate the populations over time F) OSDR also provides a phase portrait showing the direction of change of cell populations as a function of their neighborhoods, revealing fixed points.

## Results

### Approach for inferring cellular population dynamics from a spatial omics snapshot

The aim of this study is to introduce a method for inferring a dynamical model of cell populations based on a single tissue sample (Fig 1A), and to use this method to understand the dynamics of the breast cancer microenvironment and its response to treatment.

We consider biopsy sections stained for proteins that include cell type markers and cell-division markers. From this data we generate a list of cell coordinates and division status for each cell type. We used imaging mass cytometry (IMC) samples (Danenberg et al. 2022; Fischer et al. 2023; X. Q. Wang et al. 2023) and division marker Ki67 (Methods), but in principle other spatial proteomic or transcriptomic assays can be used (Methods).

The key idea is to use a spatial omics snapshot in order to estimate the division and removal rates of cells as a function of the composition of the cell’s neighborhood. We call this approach one-shot tissue dynamics reconstruction (OSDR). Its essential advance is the use of division rates measured at the level of single cells at one time-point to produce dynamics at the tissue level.

The change in cell counts within a tissue is a balance between division, death, and migration. We separate the dynamics into two parts, one that results from cell division and death within a tissue, and a second part that results from influx of cells from circulation. Our work focuses on inferring cell division and death. We then use this estimated component to quantify the net contribution of migration from external sources of cells (Methods, S3 M-P, S4 M-P). In our analyses local proliferation and removal were sufficient to explain the cell population dynamics, but this might not be the case in settings with massive influx of cells.

In order to estimate the effect of each cell type on the growth and removal of other cells, we assume that the neighborhood surrounding each cell contains growth factors secreted by nearby cells. We thus tabulate, for each cell in a slide, its type, as well as the number and types of cells within a radius *r* (Fig 1B). We use *r* = 80*um* based on measurements of in-vivo cell-cell interaction ranges (Oyler-Yaniv et al. 2017) (Methods). The ability of neighborhood composition to predict division rates, as well as the inferred dynamics, aren’t sensitive to the choice of radius *r* (S2E, S3E, S4G).

We define division events using Ki67 thresholds based on experiments in human cell lines by Uxa et al. 2021 (Fig 1C, Methods). By using data on many different neighborhoods, available from the spatial heterogeneity of the samples, we obtain the probability of division as a function of neighborhood cell composition (Fig 1D, Methods).

This approach is precise when cells across the sample obey the same underlying dynamics. In reality, varying proximity to blood vessels or tumor mass can create zones with different levels of metabolites, hypoxia, inflammation and other factors that can affect the dynamics (Setten et al. 2022). To control for these effects, we show below that the estimated dynamics are preserved across such zones by including terms for density of endothelial or tumor cells in our model (S3H, S4H). We also show that the estimated dynamics are preserved across various patient subgroups defined by external factors such as tumor genetics or stage (S3I, S4I).

Ideally one would also like a marker for cell death, but currently available markers are not considered to be sufficient to quantify the diverse forms of cell death (Kari et al. 2022). To make progress we bypass the death marker by assuming that death rate is a constant for each cell type, namely that removal rate is not affected by the composition of the cell’s neighborhood. We find that the results are robust to wide variations in the value of this constant removal rate (S3G, S4E). We therefore approximate the removal rate by the mean division rate for each cell type - an assumption that is exact in the case of (quasi) steady state tissues in which mean removal and division rates are equal.

We use the estimated models for cell division and death to produce dynamics of cell populations at the scale of a tissue. To produce a trajectory of tissue composition over time we perform the following steps. We compute the probabilities for division and death for each cell in the tissue. We then use the computed probabilities to sample division and death events for each cell. We add a new cell next to each dividing cell and remove cells that died. This produces the tissue’s composition after one time step ΔT. By repeating this process we obtain a trajectory of tissue composition over time (Fig 1E).

In addition to these detailed trajectories, our approach allows a second perspective to study the circuitry of cell interactions in a manner that doesn’t depend on the spatial organization of a specific tissue. To do so, we analyze the dynamical system of *neighborhood composition dynamics* (Fig 1F). Each neighborhood composition is mapped to a location in a state-space where each axis corresponds to the number of cells of one cell type (Fig 2A). Plotting the expected direction of change for each possible neighborhood composition produces a phase portrait (Fig 2B,C), in which arrows mark the direction and magnitude of the rate of change. The phase portrait provides an overview of the dynamics, including its stable and unstable fixed points. The figure denotes two cell types for ease of visualization, yet this approach can provide phase portraits with numerous cell types as shown below.

**Figure 2:**
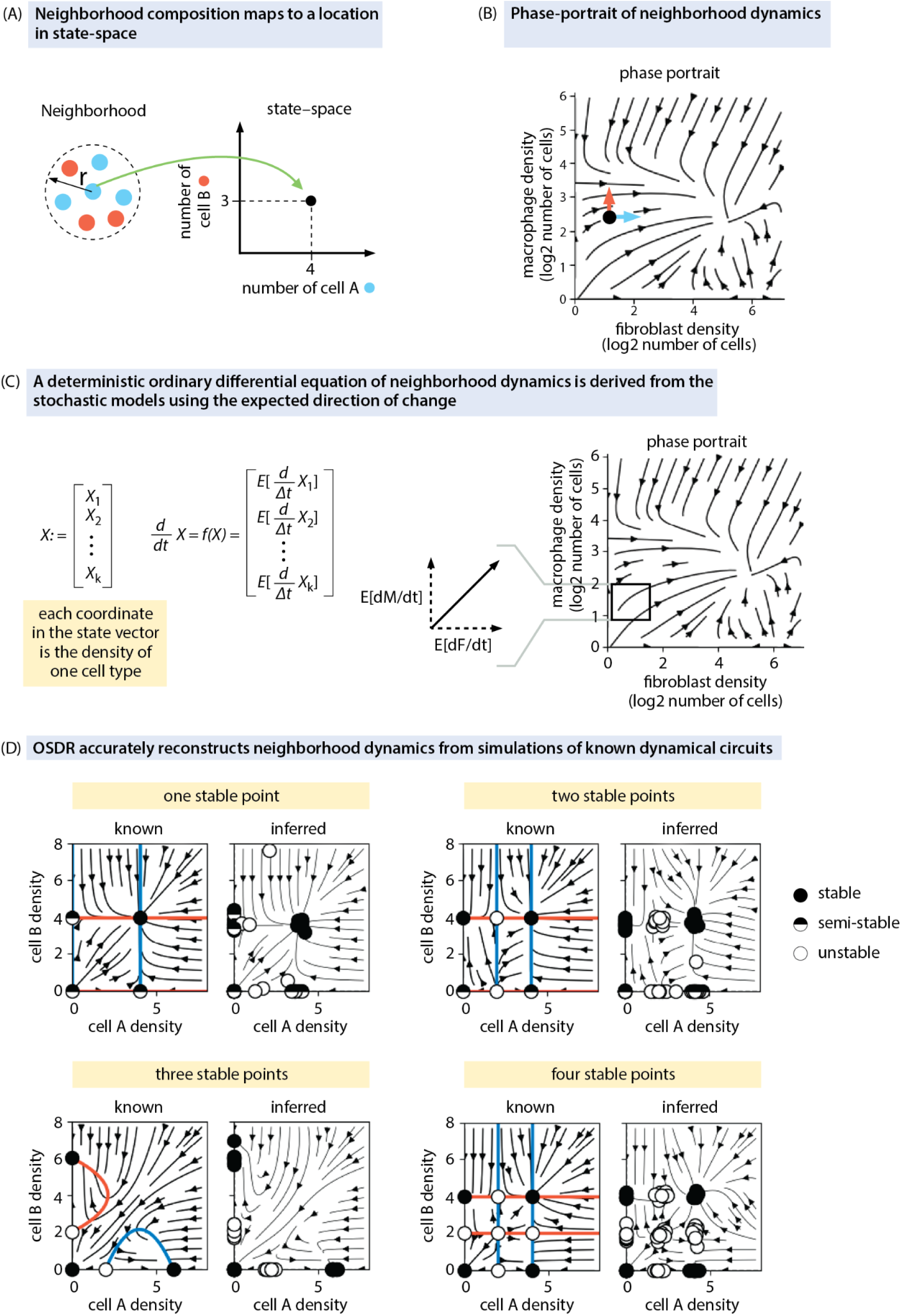
OSDR approach can recover known dynamic systems in simulations using ten thousand cells. A) Neighborhood composition maps to a location in state space. B) The phase portrait shows the change in cell populations. C) The phase portrait corresponds to a set of ordinary differential equations which are derived from the stochastic model. D) OSDR accurately reconstructed dynamics from simulations of known dynamical circuits based on simulated data of 10,000 cells of each type. ‘*Known’* panels (left) display the ground-truth phase portrait of the system used to generate the simulated spatial data. Arrows indicate directions of change and colored lines are nullclines in which only one cell type changes. Black dots and white dots are stable and unstable fixed points. ‘*Inferred’* panels display the estimated OSDR fixed points from 10 simulations, for the four two-cell phase portrait topologies. Each *Inferred* figure also displays the OSDR arrows (phase portrait) from the first simulation of the 10.

We thus obtain two outcomes: stochastic simulation starting from the tissue sample initial condition and phase portrait analysis which provides a general view of the dynamics. The two perspectives are expected to agree in the case of large well-mixed tissues (Methods). For certain spatial tissue configurations the two perspectives can produce different predictions (Methods, S1 H-M), with the stochastic simulation being more accurate for a specific tissue.

To test the feasibility of the OSDR approach, we begin with simulated data. We asked how many cells are required in order to reconstruct a known dynamical system. In each simulation we specify a certain dynamical system in which cells affect each other’s growth rate. We then simulate experimental spatial data by running the dynamics from various initial conditions of cell density, for a period of time that results in distributions of spatial cell concentrations that resemble experimental data. This creates a variety of cell compositions. We then fit OSDR to the simulated data and compare the estimated phase-portrait with the phase portrait of the known dynamical system used in each simulation.

We simulated all possible phase portrait topologies of two cell types, where each cell type can be stable on its own, or stable only in the presence of the other cell type. This results in 4 phase portrait topologies (Fig 2D). A few thousand cells of each type were sufficient to reconstruct the fixed point structure reliably (S2 G-K). We conclude that the OSDR algorithm can reconstruct simple 2D dynamics from simulated cell data and recover the underlying phase portrait given enough cells.

### OSDR reconstructs breast cancer fibroblast and macrophage dynamics in agreement with in-vitro dynamic experiments

To test ODSR, we use an extensive spatial proteomics dataset by Danenberg et al. 2022 which studied biopsies from 693 breast cancer patients using imaging mass cytometry (IMC, (Giesen 2014)). The samples include various breast-cancer genotypes and stages. Each sample is an approximately 500um x 500um tissue section. The samples include a total of 859,000 cells. Cell types were identified using standard markers and include fibroblasts, macrophages, endothelial cells, adaptive immune cells and epithelial cells. We adopted the cell-type definitions from the original publication by (Danenberg et al. 2022, S2A). The data includes the Ki67 division marker.

Since OSDR infers division probability (Ki67>threshold) based on the identity of neighboring cells, we first asked whether variation in Ki67 is indeed explained by the composition of the cell’s neighborhoods in the data. This is not obvious, because this variation may stem from unmeasured factors such as gradients of nutrients, hypoxia or inflammation (Setten et al. 2022). We performed logistic regression with cell counts as features (number of neighboring cells of each type) and Ki67 as target (Ki67 over threshold, see Methods). We find that cell-counts accurately capture a wide range of division rates (Fig 3A, Fig S2 B,C,D). All fits were significant (p-value < 10^-13^) (S2C). We conclude that the counts of neighboring cells are strong predictors of division rates.

**Figure 3:**
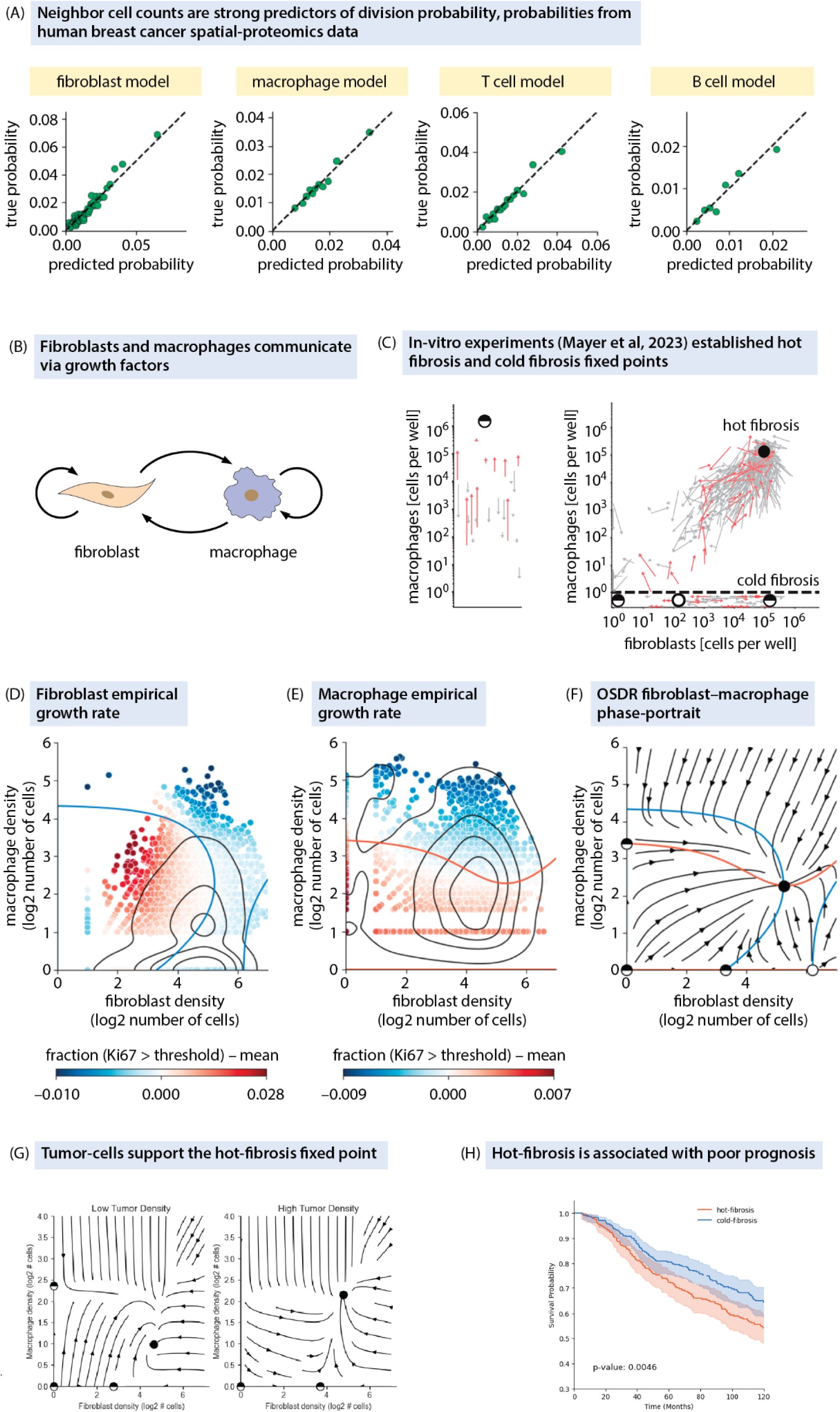
OSDR reconstructs breast-cancer fibroblast and macrophage dynamics in agreement with in-vitro dynamic experiments. A) Statistical inference using IMC data from human breast cancer biopsies (Danenberg et al. 2022) captures a wide range of division probabilities based on cell counts in the neighborhood. Each dot corresponds to a subset of 4000 cells. B) The breast-cancer microenvironment includes interactions between cancer associated fibroblasts F and macrophages M. C) The in vitro co-culture experiments of Mayer et. al. followed F and M cells in breast cancer conditioned medium (red arrows) or standard medium (gray arrows), to establish a phase portrait with several fixed points (colored circles). Arrows are changes in cell concentration over 4 days. D) Ki67 marker in fibroblasts as a function of neighborhood composition (Methods). Each point corresponds to one fibroblast and its position is the composition of that cell’s neighborhood. Colored line separates inferred regions of rising and falling cell numbers (nullcline). Black contours represent density. E) same as D for macrophages. F) OSDR inferred phase portrait for breast-cancer-associated fibroblasts and macrophages. Black and white circles are stable and unstable fixed points respectively. G) Phase-portraits of fibroblasts and macrophages in an OSDR model that includes tumor cells.

The predictive ability of neighboring cell counts could be explained by a number of factors (S2F). Growth factor concentrations are influenced by the number of secreting and consuming cells in a neighborhood (Adler 2018). Neighbor counts also reflect a cell’s chance to directly contact another cell, capturing processes such as contact-inhibition and signaling via direct contact. Some cell types influence tissue hypoxia (endothelial cells, cancer cells) and inflammation (e.g. macrophages, lymphocytes), processes that affect cell proliferation. Counts of such cell types might reflect the extent of hypoxia or inflammation in a region.

In some systems (e.g. rapidly proliferating immune or developmental contexts), the assumption might not hold that the proliferation dynamics only depend on the current observed neighborhood composition, and not on the history of neighborhood compositions. We therefore recommend testing whether the division rate of each cell is predicted accurately based only on the current number of cells in its neighborhood (as in Fig 3C and S2 B,C,D). If this is not the case, the current approach is not relevant (S2F).

We begin testing OSDR by comparing results with recent work which established *in vitro* dynamics for fibroblasts and macrophages in a breast cancer medium (Mayer et al. 2023), supplying a reference phase portrait for the dynamics of these two cell populations. The in-vitro phase portrait was obtained by seeding mice mammary fibroblasts (F) and bone-marrow derived macrophages (M) at different initial concentrations in co-cultures and following the changes in the cell populations over several days (Fig 3B). This is a measurement of the population level dynamics. In the presence of breast-cancer conditioned medium, F and M cells showed a phase portrait with several distinct fixed points (Fig 3C). Fibroblasts and macrophages supported each other in a fixed point called ‘hot fibrosis’ - in the sense that both fibroblasts and macrophages coexist. Fibroblasts alone could support themselves in a ‘cold fibrosis’ fixed point. Macrophages were induced by the cancer conditioned medium to form a macrophage-only fixed point at high macrophage densities.

The OSDR phase portraits from breast cancer samples provide dynamics for fibroblasts and macrophages (Fig 3F). We include all fibroblast and macrophage subsets defined by (Danenberg et al. 2022, S2A). Similar results are found when considering neighborhoods without adaptive immune cells, which were not present in the in vitro experiments, or including all neighborhoods and adding terms for adaptive immune cells in the model (S4 K)

Notably, the reconstructed phase portrait shares most of the key features of the in-vitro portrait (Fig 3B). There is a stable fixed point in which fibroblasts and macrophages support each other, namely a hot fibrosis fixed point. There is a cold fibrosis fixed point with fibroblasts only. There is a third semi stable fixed point with high numbers of macrophages without fibroblasts.

We tested the robustness of the OSDR phase portrait. The fixed-point structure of the phase portrait is robust to resampling of cells and samples (S3 B,C,D). It is also consistently found in a wide range of parameter choices (neighborhood radius, Ki67 threshold) (S3 E,F) and death rates (S3 G), inclusion of endothelial or tumor cells in the statistical model (S3 H), or when considering only patient subgroups with specific cancer stage, tumor size, breast-cancer genotype, patient survival time or type of treatment (S3I).

We conclude that the reconstructed phase portrait captures the salient features of the in-vitro phase portrait. Whereas the in-vitro phase portrait used dynamical measurements of cell populations over days, the reconstructed portrait uses a single snapshot with cell density and division information.

We then asked if cancer cells control the transition between cold and hot fibrosis. We used OSDR to fit a 3-dimensional model of cancer cells-fibroblast-macrophage dynamics. We then plotted 2D fibroblast-macrophage phase-portraits at increasing cancer-cell densities (Fig 3G). Increasing tumor-cell density pushes the stable fixed-point towards a “hot” state with abundant macrophages (Fig 3G, S3 K,L).

To ask whether hot-fibrosis is clinically important, we partitioned the cohort from Danenberg et al. 2022 into two groups according to presence of hot or cold fibrosis (macrophage density > half-way point between hot and cold fibrosis fixed-points), and fit a Kaplan-Meier survival curve to each group. The hot-fibrosis state was associated with poor prognosis (Fig 3H, p = 0.0046), with a decrease in median survival from 192 to 132 months. This result remained significant when the number of tumor cells was controlled for (p = 0.01, S3J).

To further test OSDR, we analyzed two additional IMC breast cancer datasets, with an additional 1014 patients and 2.1 million cells (X. Q. Wang et al. 2023; Fischer et al. 2023). Similar results were found in all three IMC breast cancer datasets (S3KL).

Panels show sections of the 3D phase portrait in low or high tumor density (tumor cells fixed at zero or at their mean density of 64 cells per neighborhood respectively). H) Kaplan-meier curves for patients whose biopsies indicate hot or cold fibrosis in OSRD analysis.

### An excitable pulse-generating circuit of T and B cells in the breast cancer microenvironment

We next considered two other cell types in the breast cancer microenvironment, T cells and B cells (Fig 4A). These cells are components of the adaptive immune system which play a major role in the cancer microenvironment and are crucial for immunotherapy (Laumont and Nelson 2023; Waldman, Fritz, and Lenardo 2020).

**Figure 4:**
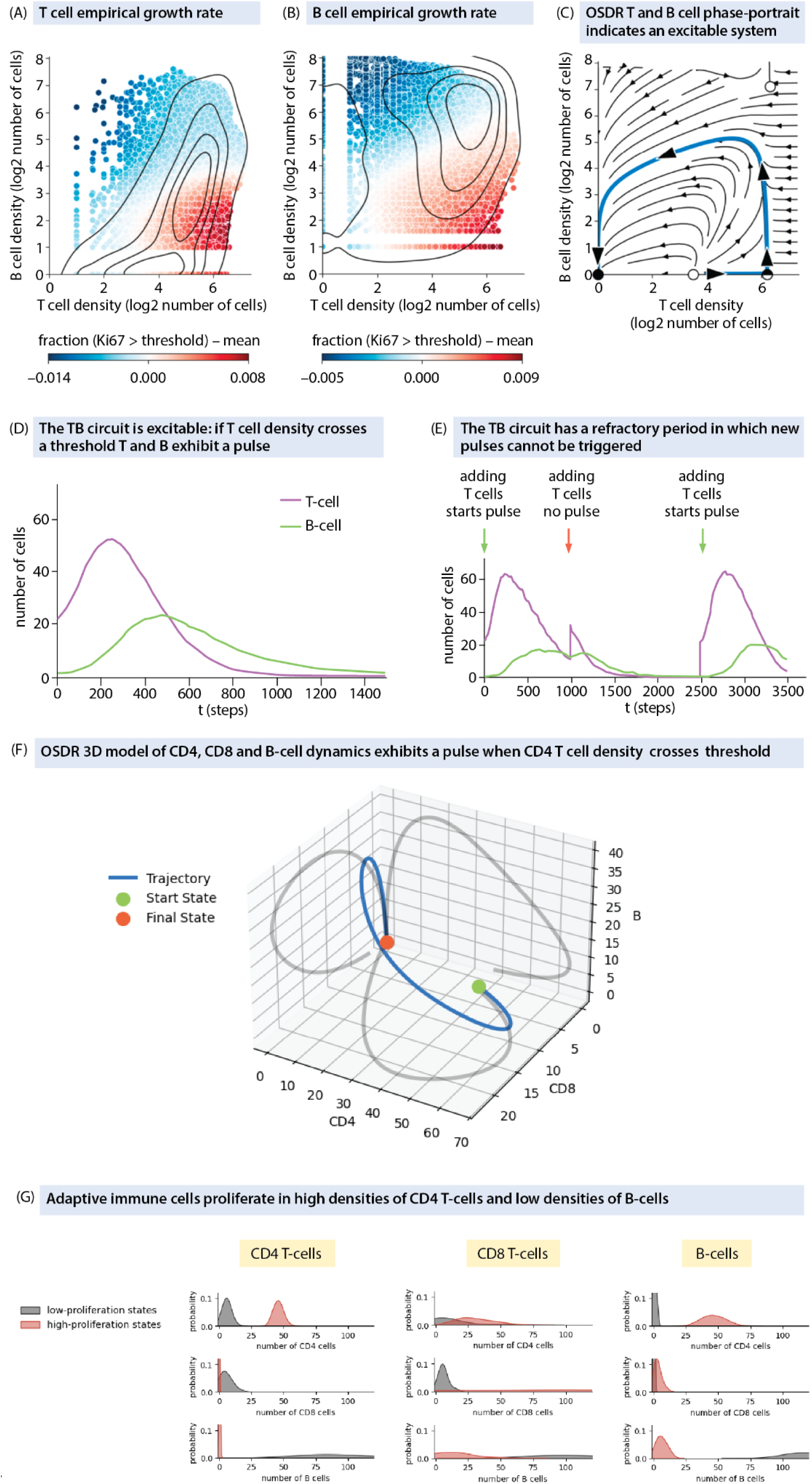
OSDR infers a pulse-generating excitable circuit of T cells and B cells in the breast cancer microenvironment. A) Ki67 marker as a function of neighborhood composition. Each point corresponds to one T-cell from the IMC dataset of (Danenberg et al. 2022). B) same for B cells. C) inferred phase portrait. The blue highlighted trajectory displays a large pulse of adaptive immune cells. D) Simulation of inferred population dynamics shows that high enough T cell levels trigger a pulse of T cells followed by B cells . E) Each pulse is followed by a refractory period, as evidenced in simulations in which additional T cells are added at different times (small vertical arrows). A new pulse is generated only when T cells are introduced after B cells from the previous pulse have declined. F) 3D phase portrait with CD4 T cells, CD8 T cells and B cells shows an immune flare with 2D projections in gray. G) Distributions of CD4, CD8 and B-cells in the 100 most proliferative and least proliferative states of each cell type. The most proliferative states are characterized by a high density of CD4 T-cells and a low density of B-cells.

We analyzed all neighborhoods in the Daneberg dataset with at least one T and/or B cell. This includes a total of 74,006 T cells and 27,643 B cells, with 1,069 and 213 cell division events respectively. Thus, about 1.4% of the T cells and 0.7% of the B cells showed division events, consistent with previous breast cancer data (Wu et al. 2021).

We estimated the dynamics between T and B cells using OSDR. The inferred phase portrait has a striking feature called excitability. It shows a stable fixed point at zero cells (Fig 4D). However, if T cells are raised above a threshold, they generate a pronounced pulse in which T cells rise, followed by B cells, and then both populations reduce to zero (Fig 4E). This pulse is similar to pulses observed in autoimmune diseases such as relapsing-remitting multiple sclerosis (Lebel et al. 2023).

Similar to the fibroblast-macrophage analysis above, the estimated dynamics were robust to resampling at the level of both cells & patients (S4 B,C,D) and were consistent under a wide range of death rates, neighborhood sizes and Ki67 thresholds (S4 E,F,G). The dynamics are not confounded by unmodeled cell types (S4 H).

The pulsatile adaptive immune response in the inferred model is shown in Fig 4E, where T cells rise, followed by B cells that inhibit them, and then both cell populations decline. The model further indicates that after a pulse there is a refractory period, where B cells are still high and T cells are low. During this period, a new pulse cannot be generated due to the inhibitory effects of the B cells (Fig 4F). A new pulse is possible only after the recovery period (Fig 4F).

To study the role of different T-cell subsets we fit a 3D model including CD4 and CD8 T-cell subsets as well as B-cells. The excitable dynamics apparent in the 2D model were reproduced in the 3D model. The 3D model revealed that a pulse is initiated when the density of CD4 T-cells crosses a threshold (Fig 4F). For all 3 cell types proliferation was highest in neighborhoods with a high density of CD4 T-cells and a low density of B-cells (Fig 4G). Proliferation was lowest at a high density of B-cells and low-density of CD4 cells (Fig 4G). We conclude that CD4 cells initiate the immune pulse whereas B-cells provide the negative feedback required to terminate the pulse.

We next considered subtypes of breast cancer from the Danenberg cohort. We compared ER+, HER2+, PR+ and triple negative breast cancer. Fibroblast-macrophage dynamics displayed hot and cold fibrosis fixed points in the phase portrait of all four subtypes (S3I). The T-B immune pulse was similar in the three receptor positive subtypes, ER+,PR+ and HER2+. However, the triple negative breast cancer phase portrait showed a difference where the pulse does not end in zero T and B cells, but instead in a fixed point where B cells remain stable (S4J). This result is consistent across breast cancer cohorts, with mixed cohorts (Danenberg et al. 2022; Fischer et al. 2023) displaying a collapse of both cells, and triple-negative cohort (X. Q. Wang et al. 2023) displaying B-cell stability. This result is consistent with the high rates of lymphocyte infiltration in triple negative breast cancer, including tumor infiltrating B cells at early stages of the disease (Toney et al. 2025; Stanton, Adams, and Disis 2016).

We also asked whether a full OSDR model with all cell types displays the same dynamics as the 2D models. We thus developed a model with the 6 major cell types in the Danenberg dataset - fibroblasts, macrophages, T cells, B cells, cancer cells and endothelial cells. We find that the 6D model recapitulates the dynamics of the 2D models when holding other cell types at their mean concentrations (S4 K,L). We conclude that the OSDR approach allows reconstruction of the dynamics of the full cell-cell interaction network in the samples.

### OSDR predicts the collapse of the tumor-cell population in treatment responders based on early-treatment biopsies

Cancer patients can go through many months of treatment without knowing if the tumor is growing or shrinking. In order to adjust treatment when it is ineffective, we need early signs of response. To address this challenge we applied OSDR to a clinical trial with three longitudinal biopsies - the NeoTRIPtrial <u>{Updating}</u>. A cohort of 279 triple negative breast cancer (TNBC) patients were randomly assigned to neoadjuvant chemotherapy (n=141) or chemotherapy + anti-PD-L1 immunotherapy (n=138) (Fig 5a). Treatment consisted of eight 3-week cycles.

**Fig 5:**
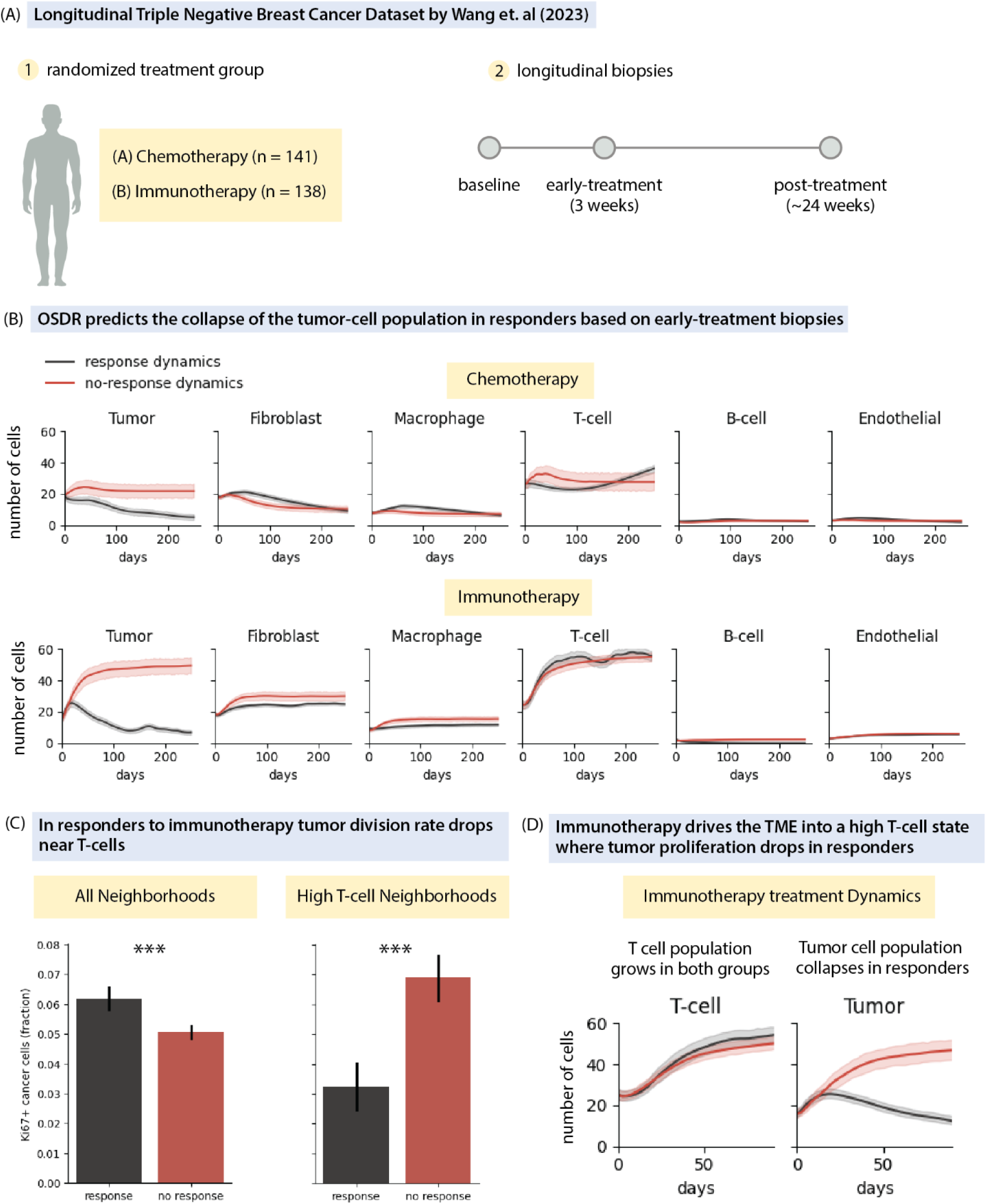
OSDR predicts the collapse of the tumor-cell population in responders based on early-treatment biopsies. A) Longitudinal triple negative breast cancer dataset by (X. Q. Wang et al. 2023). B) Longitudinal predictions for 6 cell types. OSDR models were fit separately to early-treatment biopsies from responders and non-responders in each treatment arm. The tumor cell population collapses in responders (dark gray lines) but is stable in non-responders (red). To isolate the effect of the TME dynamics, as opposed to initial tissue composition, trajectories were computed starting from the composition of all biopsies taken at the beginning of treatment. Plots display the average cell count over all trajectories. C) Fraction of dividing tumor cells in all neighborhoods and in neighborhoods with the top 10% of T cells. D) Zoom in on the first 90 days of predicted immunotherapy treatment response dynamics. T-cell density increases in both responders and non-responders, tumor cell population collapses only in responders.

Biopsies were collected at three timepoints: before treatment, 3 weeks into treatment (day of the second treatment cycle) and post-treatment (at surgical excision of the tumor following eight 3-week treatment cycles). Pathologists labeled the post-treatment biopsies as pathological complete response (pCR) or residual disease (RD).

We hypothesized that a response or lack of response to treatment should be evident in the early-treatment dynamics. We applied OSDR to week-3 biopsies from responders and non-responders in each treatment arm (total of four groups). We modeled dynamics of the six major cell types - fibroblasts, macrophages, tumor, endothelial, T- and B-cells. We then predicted the composition of the tissue over time. In order to detect differences resulting from dynamics, as opposed to differences resulting from initial tissue composition or tumor burden, we applied both responder and non-responder models to the same initial states - namely, we applied both models to every early-treatment biopsy. Fig 5b shows the predicted cell density over time, averaged over all starting states. In both treatment arms, OSDR predicts the collapse of the tumor-cell population in responders but not in non-responders. These results are robust to the choice of patients used to fit each model (S5A, P<10^-6^ for both treatment arms).

Our analysis shows that the collapse of tumor cells in responders is not a result of the initial tissue state, since we applied the models to all initial states. Moreover, the average proliferation rate of tumor cells at week 3 does not separate between responders and non-responders (S5B). In fact, the mean division rate of tumor cells in responders was higher than non-responders in the immunotherapy treatment arm (Fig 5C, P<10^-6^). Thus, the tumor cell population collapses as a result of interactions between cells over time and not initial tissue composition or average proliferation rates.

The predicted trajectories in immunotherapy show a sharp rise in T-cells (Fig 5B). This rise appears in both responders and non-responders receiving immunotherapy but not in patients in the chemotherapy arm. Because T-cell density increases in both responders and non-responders under immunotherapy, we hypothesized that T-cells have a different effect on tumor cells in these two groups. When considering all neighborhoods, the fraction of dividing cancer cells is higher in responders vs non-responders (Fig 5C left, P<10^-6^). But when we focus on neighborhoods with many T-cells (top 10% of T-cell counts) the fraction of dividing cancer cells is now 2 fold lower in responders than in non-responders (p<10^-7^) (Fig 5C).

We conclude that OSDR can predict treatment response using an early biopsy in both treatment arms (Fig 5D). It also captures the rise in T cells in immunotherapy but not chemotherapy (Fig 5D).

## Discussion

We presented the One-Shot Tissue Dynamics Reconstruction (OSDR) approach to discover cell population dynamics from a tissue omics snapshot. OSDR infers division rates as a function of neighborhood cell composition using a cell-cycle marker, and uses this to recover a phase portrait from a biopsy. We used OSDR to analyze spatial proteomics snapshots of human breast cancer tumor microenvironment (TME) samples (Danenberg et al. 2022; Fischer et al. 2023; X. Q. Wang et al. 2023). The estimated fibroblast-macrophage dynamics agree with dynamics measured directly from co-culture experiments in breast cancer conditioned medium, showing a flow towards fixed points of hot and cold fibrosis, as well as self-sustaining macrophages. OSDR also infers the unanticipated possibility of excitable immune pulses in cancer immunosurveillance - T cells and B cells in the breast TME form a pulse-generating circuit, in which T cells rise, followed by inhibitory B cells causing decline of both populations. This suggests a temporal view of the tumor microenvironment with flares of adaptive immunity.

OSDR also predicts collapse of the cancer cell populations in responders but not in non-responders from early biopsies in a clinical trial on breast cancer chemotherapy and immunotherapy. Our approach opens the possibility of obtaining cell population dynamics, and hence tissue composition over time, from snapshots of patient biopsies.

### The OSDR approach

OSDR can be applied to any experimental modality that provides measurements of cell location, type and division marker levels. Such modalities range from standard quantitative immunostaining to state-of-the-art spatial transcriptomics and proteomics (Keren et al. 2019; Giesen 2014; Janesick et al. 2023). Our results suggest that OSDR works well with data on the order of thousands of cells of each type and above. It is possible to implement OSDR using different markers for division or death.

OSDR can in principle enable an individualized dynamical model for a given patient based on a single biopsy sample. The present study used 0.25 mm^2 breast cancer samples, and used hundreds of samples from different patients to obtain large cell numbers. One can obtain the required number of cells from a single biopsy, provided that it is large enough (e.g. multiple sections from a biopsy of a few cubic millimeters totaling an area of 0.2 cm^2). This is feasible using current technology (Keren et al. 2019; Giesen 2014). One can then ask how the dynamical behavior of the sample correlates with prognosis and treatment response.

In this study we inferred division probabilities using logistic regression. The features in the regression were based on the number of cells of each type within a fixed radius. We show that these features are strong predictors of a cell’s division rate (Fig 3C, S2 B,C). Future studies may develop more sophisticated models for this task in order to utilize additional spatial features, markers and interactions.

Another interesting extension is to incorporate prior knowledge on cell motion within the spatial simulations of OSDR. For movement within the tissue, this could be done by applying a neighborhood-dependent spatial translocation to each cell at every time step in the simulation. Flux from external sources can also be modeled by placing new cells in the tissue. As an example, for immune cells this approach could include system-level dynamics involving translocation between tumor, lymph nodes and circulation, as well as local cell-cell interactions.

The cell-circuit models inferred by OSDR can be used to identify key cell-cell interactions that drive the dynamics. These key interactions are potential targets for interventions to guide tissue composition towards desired states. The effect of such interventions can be modeled by perturbing parameters in the OSDR equations that describe each interaction. As inspiration, we note recent studies on fibrosis by (Adler et al. 2020; Miyara et al. 2023; S. Wang et al. 2023) in which a cell-circuit approach identified the myofibroblast autocrine loop as a critical component, and indicated that inhibiting this autocrine loop can collapse myofibroblast populations.

Experimental inhibition of this autocrine loop reduced fibrosis in mouse models for myocardial infarction (Miyara 2023) and NASH liver cirrhosis (Wang 2023). OSDR can propose similar targets for multi-cellular circuits. For example, OSDR predicts that increasing B-cell removal rate will produce higher T-cell levels. This is a nonlinear effect resulting from the negative feedback of B cells on T cells.

### Hot fibrosis in the breast tumor microenvironment

We used OSDR to infer the dynamics of cancer-associated fibroblasts and macrophages in the breast TME. The inferred phase portrait shows that fibroblasts alone (cold fibrosis) or macrophages alone can support themselves. When both cell types are present, their numbers increase to approach a hot fibrosis fixed point where both cell types support each other. These dynamics agree with in-vitro measurements of mouse mammary fibroblasts and BMDM-derived macrophages cocultures in cancer conditioned medium. OSDR also infers that cancer cells support the transition from cold to hot fibrosis (Fig 3G, S3 K,L), a state associated with poor survival (Fig 3F).

### Inferred pulse-like excitable T cell and B cell activation

We also studied the dynamics of T and B cells in the breast TME using OSDR. These adaptive immune cells are crucial for cancer immunosurveillance and for immunotherapy. Most accounts of immune-surveillance of advanced cancer implicitly consider it as a steady-state process (Weinberg 2014), in which T cells become exhausted. In contrast to this static picture, we infer a dynamic process with pulsatile T and B dynamics. The phase portrait suggests that CD4-T cells, once they cross a concentration threshold, induce their own proliferation and that of CD8 T-cells and B cells, whereas B cells have an inhibitory effect on the growth of the three cell types.

These inferred interactions align with immunological principles. T cells promote their own activation and proliferation, as well as that of B cells, via cytokines such as IL-2, IL-4 & IL-21 (Murphy and Weaver 2016; Butler 2012). Regulatory roles of B cells in the TME have also been documented (Wolf H. Fridman et al. 2022; Engelhard et al. 2021; Laumont et al. 2022; Murakami et al. 2019; Cai et al. 2016; Shalapour et al. 2017; 2015; Yuen, Demissie, and Pillai 2016). Such regulatory B cell effects have been observed in human breast cancers and associated with poor prognosis (Ishigami et al. 2019). Regulatory B-cell activity is, in part, induced by cytokines released by T-cells such as IL-21 (Yoshizaki et al. 2012).

Together, these interactions can generate pulses of activity when activated T cells cross a concentration threshold. Such pulses or flare-ups are observed in autoimmune diseases (Lebel et al. 2023; Oleinika, Mauri, and Salama 2019; Yoshizaki et al. 2012). A rise and fall of adaptive immune cells with time is also a hallmark of pathogen response (Murphy and Weaver 2016). The inferred timescale of the pulses in the present study, on the order of weeks, roughly agrees with the flare duration in the autoimmune and pathogen contexts.

The T and B cell pulsatile circuit also predicts a refractory period after each pulse. During the refractory period, a new pulse of T cells cannot be started due to high levels of inhibitory B cells. Only after B cells have declined can a new pulse be generated. This refractory period is similar conceptually to that found in neuronal action potentials, also an excitable system (Hodgkin and Huxley 1952).

A T cell refractory period can potentially bear on the timing of cancer immunotherapy. It may be beneficial to time immunotherapy doses so that they don’t overlap the refractory period of the previous dose. Removing such ineffective doses can help minimize immune-related adverse events which currently cause treatment termination in up to 30% of patients (Martins et al. 2019). More generally, the existence of such T and B cell pulses can conceptually add a temporal dimension to research on the TME, with flares of adaptive activity.

Adaptive immune flares can also address the pleiotropic effects of B cells observed in tumors. On the one hand depleting B cells in mice improved outcome in several cancer models (Qin et al. 1998; Shah et al. 2005; Brodt and Gordon 1978; Barbera-Guillem et al. 2000; Lee-Chang et al. 2013). On the other, B cells in human tumors (primarily in tertiary lymphoid structures) are associated with improved prognosis in many cancers (Laumont et al. 2022; Wolf H. Fridman et al. 2022, 20; Wolf Herman Fridman et al. 2020). One possible explanation is that observing high B cell counts in tumors indicates that T cells have crossed a sufficiently high level for a flare of immune attack; depleting B cells may reduce their inhibition and sustain the T cell pulses for longer durations.

Triple negative breast cancer samples showed a variation on the excitable system, in which B cells remain high after the pulse. This aligns with the immunogenic nature of triple negative breast cancer, displaying high rates of tumor infiltrating lymphocytes (Toney et al. 2025; Stanton, Adams, and Disis 2016).

### OSDR can predict response dynamics in response to chemotherapy and immunotherapy

The response to cancer treatment is often heterogeneous. Some patients respond while others do not. Ineffective treatment not only risks unnecessary side-effects, but also comes at the expense of alternative treatments that could potentially work. Thus, we would like to know if treatment is working as early as possible.

We applied OSDR to a triple-negative breast cancer clinical trial where biopsies were taken soon after the start of treatment (week 3 of 24) and again at the end of treatment. The trial included two treatment arms where patients received either chemotherapy or a combination of chemotherapy and immunotherapy. Using the early biopsies OSDR accurately predicted the collapse of the tumor cell population in responders. This result was reproduced in both treatment arms and was robust to the subset of patients used for fitting the model (S5A).

OSDR provides insight into the dynamics of immunotherapy. Initially, there is a low density of T-cells and the proliferation rate of tumor cells is similar in both responders and non-responders. Then T cells rise in both responders and non responders. The difference between the two groups emerges only after T cells rise, where cancer cells collapse in responders but not in non-responders.

OSDR picks up this difference by virtue of a neighborhood effect in the early biopsy - in future responders, tumor proliferation is low near T-cells, but not in future non-responders. As a result, when T-cell levels rise the tumor cell population begins shrinking.

The low tumor proliferation near T-cells in future responders may thus serve as a functional marker of response to checkpoint immunotherapy. Similarly, high tumor proliferation near T-cells could be an early marker for identifying a subset of patients harmed by immunotherapy termed “hyper-progressors” (Champiat et al. 2017; Adashek et al. 2020; Leake 2023; Sehgal 2021; Kato et al. 2017). In these patients, the tumor proliferates more rapidly following immunotherapy.

### Limitations and Caveats

Application of OSDR requires both domain knowledge and awareness of model limitations. First, OSDR estimates dynamics at a given point in time. It does not model the changes in the dynamics themselves. As such, OSDR will be inaccurate if dynamics significantly change. For example, in the cohort by (X. Q. Wang et al. 2023) cell division rates drop considerably over the 24 weeks of treatment (S5C). This could be a cumulative effect of chemotherapy. Prior knowledge or samples from multiple time points can be used to establish the time-scale of the changes in dynamics. Rapid changes in tissue composition could also have an effect that history rather than current neighborhoods dominate the dynamics (S2F). In all cases, plots such as Fig 3C, S2B,C should be used to test that neighborhood compositions predict proliferation rates.

The current implementation approximates the death rate for each cell type as independent of the neighborhood, equal to the mean division rate in the sample. Establishing reliable markers for cell death rate may improve model estimates in cases where regulation of death rate by neighboring cells dominates proliferation.

When sampling tissues it is important to obtain a large variance of neighborhood compositions. By sampling a large tissue section or multiple biopsies, one can observe different tissue regions in different stages of the dynamic processes (S5D).

To employ OSDR, one needs to consider confounders. First, when using data from multiple patients one needs to check that a result is stable across different subsets of patients (e.g stage, genotype, tumor size). Reproducing the dynamics across patient subgroups ensures we do not overinterpret multiple events from different tissues as a trajectory. A second type of confounder could occur spatially within the tissue, e.g hypoxia or inflammation. This can be tested by including terms for endothelial, immune and tumor cells, which serve as proxies for hypoxia and inflammation. These terms are automatically included in full OSDR using all cell types. If key cell subtypes are crucial for the dynamics but are not detected by the experimental markers it might be impossible to test this type of confounder.

OSDR captures cell division and removal within the tissue, so predictions can deviate from observations when transdifferentiation or migration play a dominant role in the cell population dynamics. We can identify cases where transdifferentiation or migration terms are required to explain observations by plotting the sources and sinks of unexplained components of the dynamics, as in figures S4 M-P. For example, in the case of T- and B-cells OSDR predicts a single stable point with zero cells of each type. An external source of cells is required to explain the observation of tissues with positive densities of each cell type. Accordingly, figures S4 M-P suggest that influx of lymphocytes drives tissues with no lymphocytes to the point of initiation of an immune flare. Thus, observed high-densities of lymphocytes are explained by local proliferation (i.e by OSDR) rather than external influx. In Fig. 4 D,E we simulate this migration component by adding cells to the tissue. Future implementations of OSDR could directly model transdifferentiation or migration components.

In the present work we compared OSDR predictions against pre-post therapy biopsies from human patients, over a 24-week course of treatment (Fig 5). Direct measurement of TME dynamics in vivo can provide another opportunity for validating dynamics on a finer timescale in animal models. Possible methods include mouse models of tumor development (which have less inter-sample heterogeneity), pulse chase experiments and intravital microscopy (Entenberg, Oktay, and Condeelis 2023).

### Summary

We presented the OSDR approach to infer cell population dynamics in a tissue based on spatial snapshots from biopsies. We demonstrate the utility of OSDR in understanding the breast cancer microenvironment. We validated OSDR by comparison to macrophage and fibroblast dynamics in vitro, and used OSDR to infer T and B cell pulses in the breast tumor microenvironment, adding a temporal dimension to cancer immunosurveillance. OSDR predicted collapse of cancer cells in treatment responders versus non-responders and provided insight on the dynamics of immunotherapy treatment. OSDR could help discover cell population dynamics in other tissues and conditions based on biopsies - including developmental processes and pathological processes such as autoimmunity or fibrosis. Tissue dynamical models provided by OSDR can inform mechanisms of pathology and provide targets for treatment aimed at changing tissue composition to reach therapeutic goals.

## Methods

The number of cells of a given type in a tissue section can change by division or death, moving in or out of the section (flux) or transdifferentiation (Weinreb et al. 2018). If we restrict ourselves to modeling cell-types who do not transdifferentiate at appreciable rates such as T-cells and Macrophages and analyze tissues in which flux is negligible relative to rates of death or division or add flux terms post-hoc, we remain with the following equation for the rate of change of a population of cells of type *i*:

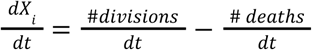

OSDR aims to transition from static observations of cell division or death in a tissue into rates. The key insight is as follows - if we obtain a marker for cell division (or death), and in each cell division the marker remains above a defined threshold for a time period *dt*, then all observed divisions occurred within the last *dt* hours. Thus:

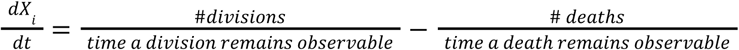

A cell’s rate of division or death is influenced by the signals that it receives from its environment, its access to nutrients, its genetics and factors such as physical contact with other cells. We call this complete set of factors the cell’s “neighborhood”. This definition sets an ideal, and we denote the particular set of features used to approximate this ideal as *N*(*x*), where *x* is some cell in the tissue.

If we consider cells with identical neighborhoods, a fraction of them will be dividing. This fraction is higher if the neighborhood induces a high rate of division and vice versa. We thus view the observations of division or death as random events whose probabilities are determined by the cell’s neighborhood. Thus, for cell *x* of type *i*, the distribution of the observation *O_i,t_*(*x*) of a division or death event is modeled as follows:

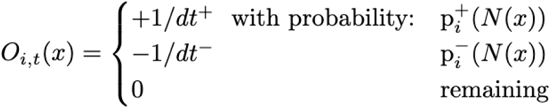

Where p ^+^, p_i_ ^-^ are the statistical inference-models for division or death, and dt^+^, dt^-^ are the durations of observed division or death markers respectively. In this study, dt^+^ is defined as 1 time unit (roughly a few hours, see (Uxa et al. 2021)), and the approximation of the death rate as the mean division rate is defined using the same time units. Thus, in this implementation dt^+^=dt^-^=1. We divide by the durations so that we get the following simple formula for the (stochastic) change in the number of cells in each timestep:

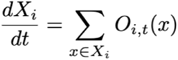

Tissues are heterogenous, and the diverse cellular compositions in different regions result in various directions of change. In order to analyze the change at a certain *state*, rather than the change in complete tissues, we compute the expected change with respect to an initial condition where cells share the same neighborhood (mathematically equivalent to a mean-field approximation of a large tissue with lots of cell motion and growth factor diffusion, Methods):

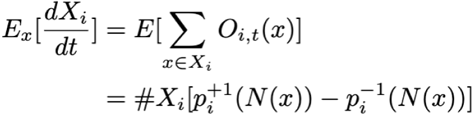

In this study, features are based on the number of neighboring cells of each type within a certain predefined radius. For further discussion on this choice of features see S2F. The neighborhoods can thus be represented as a vector of cell densities:

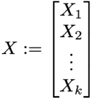

As a result, we can interpret the models as components of a set of ordinary differential equations defined over a state-space of cell densities:

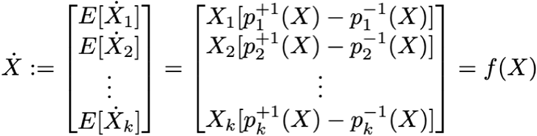

Note that OSDR is inference-model-agnostic, while we employ logistic regression here, other statistical inference models can be adopted within this framework. In addition, this approach can be applied to data acquired by any technology that enables classification and spatial localization of discrete cells, together with robust measurement of a set of markers for division (and possibly death).

### Model Inference Algorithm

<u>Input:</u>

Cell level annotations: 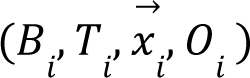 for cell *i* = 1,… *N*

- *B_i_* - an ID for the biopsy sample cell *i* came from. *B_i_* is used to formalize that a cell can only influence cells from the same tissue.
- *T_i_* ∈ *T* - cell type for cell *i* (e.g *T_i_* = “Fibroblast”), and let *T* be the set of types.
- For any *t* ∈ *T* let *n_t_* be the total number of cells of type *t*. We shall also assume some order on the cells of type *t*, such that: *t*[*j*] = *i* for any index *j* ∈ {1,…, *n_t_*} and the original index *i* ∈ {1,… *N* }.
- 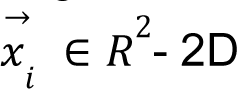 spatial coordinates for cell *i*.
- *O_i_* ∈ {0, 1} - binary observation of division

<u>Algorithm:</u>

For each cell-type *t* ∈ *T*,:

1. Compute a table of features 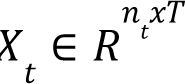.

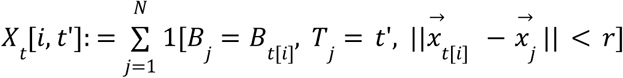 Here *t*[*i*] is the original index of the i’th cell of type *t*. The value at row *i* and column *t*’, is a count of all cells that came from the same tissue as cell *t*[*i*], that have type *t*’ and are within a distance *r*. Row *i* in *X* is our representation for the “neighborhood” of cell *t*[*i*] - the counts of all cell-types in its proximity. Denote this vector by 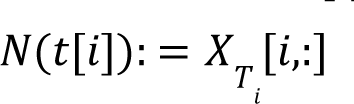
2. Perform feature transformations on *X*, such as adding interaction terms. (transformations are selected through a separate process of cross validation)
3. Define the binary labels *y_t_* ∈ {0, 1}*^n_t_^*:

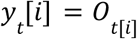
4. Fit a multivariate logistic regression model *p*^+^_*t*_ for division rate based on the features and labels *X_t_*, *y_t_*.
5. Define death rate: *p*^−^_*t*_ = *mean*(*y_t_*)

<u>Output:</u>

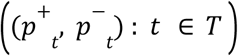

### Ki67 Thresholds

We used previous data to establish a Ki67 threshold for cell division events. Uxa et. al, 2021 demonstrated that Ki67 levels peak towards the G2/M phase of the cell cycle, with preserved kinetics (up to scale) across two human and one mouse cell line. We adjust Ki67 levels to correct for different scaling and division rates between cell lines (S1 A,B) by selecting Ki67 values above a noise threshold *T_n_*, subtracting *T_n_* and dividing by the Ki67 standard deviation. We choose *T_n_* = 0. 5 mean isotopic counts because this is the typical magnitude of experimental noise in this dataset (S1C). This produces distributions with similar shape (S1D) in accordance with the cell line results of Uxa et al. We define a cell division by the normalized Ki67 above a division threshold *T_d_*. Example fractions of dividing cells are provided in S1E. The resulting model estimates are robust to *T_d_* values between 0 and 1 on this adjusted scale (S3E, S4G)

### Choice of spatial proteomics technology

Out of the currently available spatial-omics technologies, multiplexed protein imaging(Giesen 2014; Keren et al. 2019) was the most suitable for our setting. Current barcode-based spatial-transcriptomics (Tirado-Lee, n.d.) aggregate cells in spots of 55 micron diameter so they don’t associate transcript levels with single discrete cells. Also, because the distance between spot centers is 100 micrometers, if we place a cell in the center of a spot, we measure less than 1/9 of the area immediately surrounding it (area of a circle of radius 27.5 micrometers, within another of radius 72.5). Fluorescent based approaches like MERFISH allow modeling single cells but they only recover a small fraction of every transcript in the tissue. This is a barrier for basing the analysis on levels of a single transcript - Ki67. IMC provides measurements within well-defined cells, reliable measurement of Ki67 and, in terms of cost, makes datasets in the order of magnitude of hundreds of thousands of cells feasible. Classical staining methods should also be feasible for analyzing specific pairs of cells. For example, for analyzing fibroblasts and macrophages, we could use 4 markers: fibroblast and macrophage markers, a cell nucleus marker, and Ki67.

### Enforcing Maximal Density

We assume there cannot be a net positive flux at the maximal observed density of a cell type. We apply a correction in the form of:

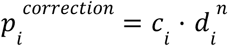

Where d_i_ is the density of cell i, and the power n controls the steepness - high n implies the correction applies primarily to higher densities (S1F). The constant term c_i_ is defined as the minimal correction ensuring non-positive flux at max density (D_i_). We compute division minus death rates at the maximal density of one cell, over all possible values of the second cell:

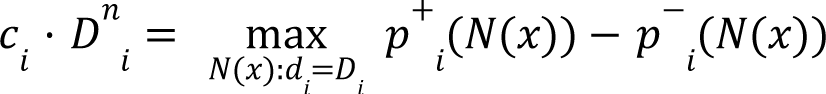

In practice, the state-space region near maximal density for both cell types is unpopulated by cells, so we use values up to the 95% quantile density for each type. This is more robust to artifacts due to extrapolation in areas with minimal data and produces smaller corrections (S1G). For the FM model this correction is trivial because the estimated net flux is negative at high densities, for the TB model the correction applies only to T-cells (S1G,H).

### Population vs Neighborhood Dynamics

In OSDR we first estimate cell-level models for division and death probabilities (Methods, Model Inference Algorithm). We then use the cell-level models to analyze tissue dynamics in one of two ways.

We call the first approach “population-level dynamics”. We start with a spatial tissue section. We then produce a temporal sequence of tissue snapshots as follows. We first compute the probabilities of division and death for each cell, taking into account the composition of each cell’s neighborhood. We then use the probabilities to sample for each cell - division, death or none. We place each new cell in the neighborhood of the dividing cell. We remove cells that died (S1H).

One nuance in this process relates to cells at the edge of the tissue. When a cell near the edge of the tissue divides, its daughter cell might be placed outside the tissue. To keep tissue dimensions fixed we remove cells placed out of bounds. This biases the neighborhoods of cells near the edge of the tissue. To correct this bias, we rescale the cell counts of cells near the edge of the tissue: 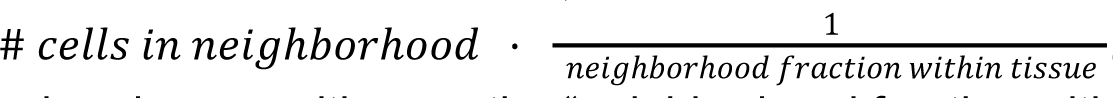. This is an unbiased estimator of neighborhood composition, as the “neighborhood fraction within the tissue” is also the probability of keeping a daughter cell following division.

We can also add cell-movement to this sampling procedure. We implemented a random walk by sampling a gaussian translation to each cell at each time step. Note that a large diffusion coefficient can disperse cells out of tissue boundaries (S1I), while a small diffusion coefficient can cause regions in the tissue to be effectively isolated (S1J). More elaborate modes of cell-movement include attraction towards a chemokine source, adhesion to nearby cells and migration of cells from outside the tissue. Studying the effects of different modes remains outside the scope of this current work.

To study a sequence of tissue sections we can produce a graph of the number of cells over time (e.g Fig. 4E,F). Because tissues can have different dimensions, we find it useful to rescale the number of cells to the area of a neighborhood. Thus, the y-axis displays:

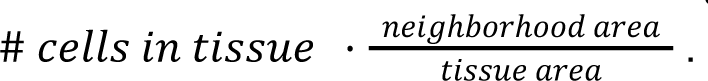

One limitation of plotting the average density is that we don’t observe the complete distribution. Notably, the average density doesn’t necessarily coincide with the mode. S1K shows a simulation where the mode locates the stable fixed-point of the system, whereas the average density is strongly biased.

Thus, to analyze population-dynamics we need to specify an initial spatial tissue configuration and a diffusion coefficient. We must also consider the distribution of neighborhood densities across the tissue. The second approach for analyzing tissue dynamics doesn’t demand such choices.

We call the second approach “neighborhood-level dynamics”. Here we analyze an isolated neighborhood where all cells have the same neighbors. We then analyze an ordinary differential equation (ODE) obtained by computing the average direction of change of each cell type (Methods). To analyze this ODE it is convenient to plot phase-portraits for 2D models (e.g Fig 2B,C,D, 3F, 4C), and trajectories for higher dimensions (e.g Fig 5B).

Mathematically, the neighborhood dynamics can be derived from the population dynamics through the law of large numbers. Denote the tissue area as *A*, and fix a neighborhood area *a*. Let *X* be a vector with the number of cells of each type in the tissue. The change in the number of cells of type *i* is given by sampling a division or death event for each cell: 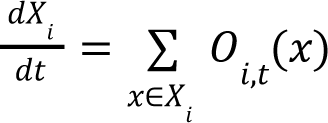, where *O*(*x*) = 1 with probability *p*^+^, *O*(*x*) =− 1 with probability *p*^−^ and otherwise *O*(*x*) = 0 (recall, *N*(*x*) is a vector of the number of cells of each type in the neighborhood of cell *x*). If we assume that the rates of cell motion and cytokine diffusion are large relative to the timescale of cell proliferation we can assume that, effectively, 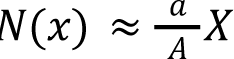. In this case, for all cells *x* the probabilities of division and death are the same 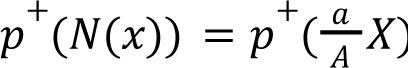, 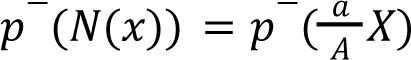. Now, if we scale the equation by 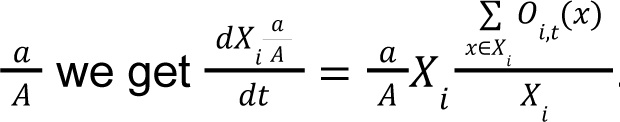. If we take a limit on the size of the tissue, while fixing the density of cells 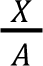, then for all *i*, *X_i_* → ∞ and the empirical mean of the independent and identically distributed variables *O*(*x*) approaches their mean. We get: 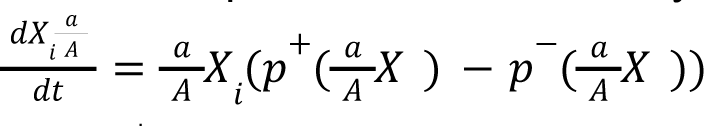. We get the neighborhood dynamics equation: 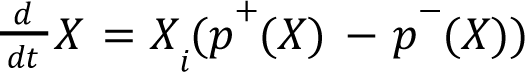 by replacing notation from 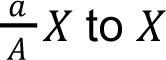

Generally, population and neighborhood dynamics don’t have to agree. The population model is stochastic and discrete, while the neighborhood model is deterministic and smooth. We constructed examples to highlight two important differences between the population and neighborhood approaches.

The first type of discrepancy results from collapse of a cell-population. In the deterministic neighborhood dynamics, a cell-population will always flow in the average direction of change. Thus, if a cell-population has a positive stable fixed-point, it will deterministically flow towards it. On the other hand, in the stochastic population-dynamics, a cell-population can move against the average direction of change. If due to a random fluctuation, all cells in a population die, the population won’t recover (similar to a gambler’s ruin). This can produce large differences between neighborhood and population dynamics. A collapse is more likely to occur for cells with a low-density fixed point. It is also more likely without diffusion. Adding even a little bit of diffusion allows neighborhoods with higher densities to “rescue” neighborhoods that collapsed. S1L shows that the macrophage population eventually collapses without diffusion. Adding a bit of diffusion produces similar steady states under both models (S1L).

A second type of discrepancy results from the initial tissue configuration. Population-dynamics depend on the initial tissue configuration, whereas neighborhood-dynamics do not. Certain particularly “adversarial” tissue configurations, coupled with low cell-motion, can produce large discrepancies. For example, under neighborhood dynamics, the T- and B-cell model produces a flare from a location of (4,1) on the TB phase-portrait (Fig 4C). Under population-dynamics, a flare also occurs when we initialize a tissue by placing cells at random (S1M, 1st example). But we can organize the same number of cells such that no flare occurs. For example, we can place T-cells at one side of the tissue and B-cells at the other. In this case, the B-cell population collapses because the B-cells don’t have T-cell neighbors (S1M, 2nd example). Another option is placing all B-cells in the same place, creating a high density of B-cells locally. In this case the low density of B-cells at the tissue-level doesn’t reflect their high density locally (S1M, 3rd example). In both examples, we used our understanding of neighborhood dynamics to design a tissue configuration that produces a discrepancy.

### Validating OSDR by simulating known dynamical models (Fig 2E)

We simulate spatial data based on a known dynamical model (i.e predefined functions p^+^ and p^-^) using the following procedure:

1. Sample a random initial number of cells for each cell type.
2. Sample a random spatial position in the tissue for each cell.
3. For n steps:

a. Compute for each cell the probability of division or death based on its current neighborhood.
b. Sample an event of division, death or none based on the computed probabilities
c. Remove dead cells and place a new cell next to each dividing cell (here, location of new cells is sampled uniformly within the neighborhood. It’s straightforward to incorporate knowledge about cell motility at this stage, if available).

This produces a tissue section whose density and spatial distribution is biased by the dynamics, as we assume is the case for real tissues. We repeat the procedure 100 times from various initial densities and sample with replacement 50K cells evenly from all tissues. From this pool we then sample 1K, 5K, 10K or 25K cells for the model fits.

Because empirical distributions in our data had tails towards lower cell-densities, we sampled the initial cell densities from a beta distribution biased towards lower densities (parameters 2,4 scaled to a maximal value of 7). Model parameters were selected to produce division rates in ranges that resemble real data (mostly within 1%-6%). The fraction of divisions was approximately 2% for all models. We selected the number of simulation time steps as that required to produce distributions qualitatively similar to real data.

To evaluate model fits we tested whether the correct location and type of stable or semi-stable fixed points were recovered. To account for discretization error, if a stable point on an axis was recovered, and an unstable point was located within 1 cell from that point, we count it as a semi-stable. Such a point effectively has no basin of attraction. The same precaution should apply to interpretation of model fits in general.

For a detailed analysis of the simulations of each ground truth model see S2C,D,E,F,G.

### Plotting Ki67 levels as a function of neighborhood composition

Figures 3D,E and 4A,B display the measured Ki67 > Threshold as a function of neighborhood composition. To transform the cell-level binary division events into rates, we compute a local average and subtract the mean division rate. For the local average, we used a gaussian smoothing kernel over the cell-density state space, providing a non-parametric plot of the division rate. The kernel’s gamma parameter controls the degree of smoothing. We plot a value of gamma that produces contours of similar complexity as those of the estimated parametric models.

### Estimating magnitude of error due to unmodeled terms such as cell migration

We apply the Fokker-Planck equation to the motion of tissues through the state-space of cell densities. Each tissue is analogous to a particle moving through a space whose coordinates are defined by the densities of each cell type. For example, the location *x* could be the density of fibroblasts and macrophages (i.e *x* = (*F*, *M*)). The velocity through this space is determined by the rate of change in each cell type. This velocity has a deterministic component composed of *v⃗*(*x,y*), the division minus removal rates we previously estimated using division markers, and *u⃗*(*x,y*), which includes unmodeled terms such as cell migration and differentiation from stem cells. The velocity also includes a stochastic component which reflects fluctuations velocity of each cell type.

Applying the Fokker-Planck equation to our setting:

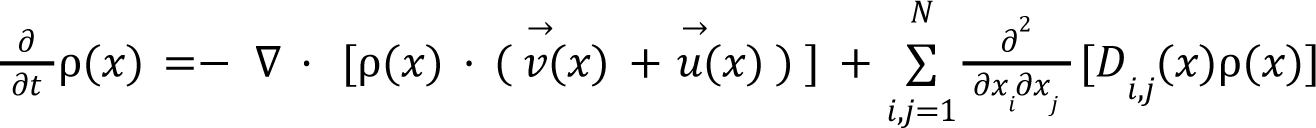

Notation:

∇ · is the divergence operator.

ρ(*x*) - density of tissues at location *x* in the state-space

*v⃗*(*x*) + *u⃗*(*x*) - velocity of a neighborhood in the state-space.

*v⃗* (*x*) - division minus death rate field, estimated from data.

*u⃗* (*x*) - velocity field due to unmodeled terms such as cell migration (will be estimated now).

*D*(*x*) - diffusion coefficient matrix. We assume fluctuations are proportional to the number of cells (e.g changes in nutrient availability modify growth rates of existing cells, rather than adding or removing cells at a constant rate).

Formally, 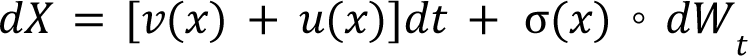. Where *dW_t_* is an N-dimensional brownian motion and σ(*x*) = σ · (*X*,…, *X*)^*T*^ multiplies each component independently. Overall, for *i* ≠ *j D_i,j_*(*x*) = 0 and for 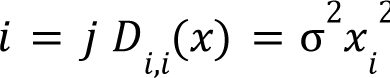. The diffusion term becomes: 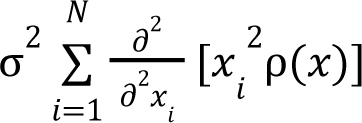. We henceforth denote the scalar 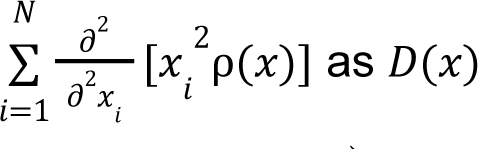. The equation becomes:

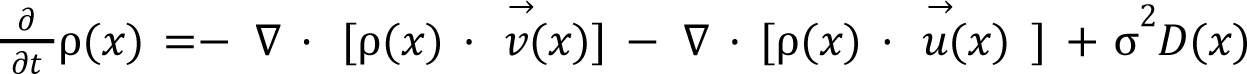

For a large sample of patients, we can assume that sampling the same tumors within a few days would produce approximately the same distribution. This implies that 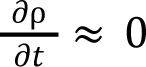. Thus:

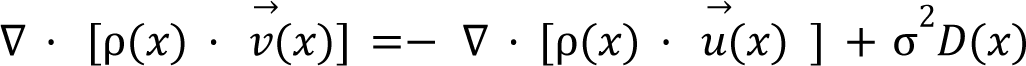

The left hand side is inferred from data for all *x*. Recall, *v⃗*(*x*) is estimated from division markers and we estimate ρ(*x*) by computing the location of each patient’s tissue in the state-space followed by kernel density estimation. We then numerically evaluate the divergence ∇ · (ρ(*x*) · *v⃗* (*x*)) using finite differences.

By plotting ∇ · (ρ · *v⃗*) we can visualize the net contribution (in units of change in density per unit time) of the missing terms. In the special case where σ ≈ 0, we directly obtain the divergence of the error (e.g migration) field as ∇ · (ρ · *u*) =− ∇ · (ρ · *v⃗*). Plotting ∇ · (ρ · *v⃗*) thus identifies the sources and sinks of the error field, as well as the magnitude of error. Note, for interpretability we divide the rates by the maximal ρ so that units are in [fractions of maximal density / Δt].

To quantify the contribution of diffusion versus deterministic terms (migration etc.) we solve an ordinary least squares problem with a single parameter σ:

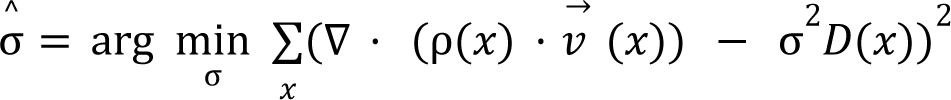

The variable *x* is taken over a discrete 2D grid. Solving for σ this way implies that the deterministic error velocity *u*(*x*) shouldn’t include components that could be explained by stochastic diffusion.

## Supplementary

### List of supplementary figures

● Supplementary 1 - Method:
  ○ S1A - Empirical Ki67 distributions have right tails
  ○ S1B - Varying fractions of cell with Ki67 > 0 or Ki67 > noise among different cell types
  ○ S1C - Typical magnitude of noise of IMC protein markers
  ○ S1D - Adjusted Ki67 distributions have similar shapes
  ○ S1E - Fraction of dividing cells (adjusted Ki67 > threshold of 0.5)
  ○ S1F - maximal density correction power-term n determines steepness
  ○ S1G - maximal density correction in the TB model - separating the original and correction components of the phase portrait (power n=8)
  ○ S1H - Population-dynamics produce a sequence of tissues by sampling division and death events for cells in the tissue
  ○ S1I - A high diffusion coefficient can disperse cells out of tissue boundaries
  ○ S1J - A low diffusion coefficient can create “isolated” regions in the tissue:
  ○ S1K - Example where average cell-density underestimates the stable fixed-point, while the mode is exact
  ○ S1L - Collapse of cell populations in the stochastic population-dynamics model can produce large discrepancies between neighborhood and population dynamics
  ○ S1M - Certain initial tissue configurations can produce significantly different trajectories
● Supplementary 2 - Model Estimation:
  ○ S2A - Cell-type definitions used in each dataset
  ○ S2B - Calibration plots for all cell types, compared with calibration on same data with permuted divisions
  ○ S2C - Logistic regression model log-likelihood ratio p-values and example model summary
  ○ S2D - Example summary of model fit
  ○ S2E - model performance as a function of neighborhood radius
  ○ S2F - The ability of current neighborhood to predict gradients and growth factors that determine proliferation, based on separation of timescales between signaling and proliferation dynamics
  ○ S2G - overview of 2-cell simulations
  ○ S2H - Simulation 1 - One fixed point
  ○ S2I - Simulation 2 - Two fixed points
  ○ S2J - Simulation 3 - Three fixed points
  ○ S2K - Simulation 4 - Four fixed points
● Supplementary 3 - Fibroblast-macrophage dynamics
  ○ S3A - Model selection using cross-validation suggests a 2nd order polynomial
  ○ S3B - The Fibroblast-Macrophage phase-portrait is robust to cell-level and patient-level bootstrap resampling
  ○ S3C - The Fibroblast-Macrophage phase-portrait is robust to patient-level bootstrap resampling
  ○ S3D - Fibroblast-Macrophage vector field without smoothing
  ○ S3E - Number of cells and divisions as a function of neighborhood size and Ki67
  ○ S3F - The Fibroblast-Macrophage phase-portrait is robust to choice of Ki67 threshold and neighborhood size
  ○ S3G - The Fibroblast-Macrophage phase-portrait is robust to perturbation of the death estimate
  ○ S3H - Fibroblast-Macrophage dynamics are consistent when modeled together with a third cell types
  ○ S3I - Fibroblast Macrophage Dynamics are conserved across various patient subgroups
  ○ S3J - Cox proportional hazards model for survival as a function of tumor and macrophage density shows a significant correlation between macrophage density and poor survival
  ○ S3K - The FM phase-portrait exhibits cold and hot fibrosis across 3 independent datasets
  ○ S3L - Phase-portraits under increasing densities of tumor cells in 3 datasets
  ○ S3M - Estimating magnitude of error due to unmodeled terms: Empirical position of tissues in state-space
  ○ S3N - Estimating magnitude of error due to unmodeled terms: Estimated density of tissues across the state-space
  ○ S3O - Estimating magnitude of error due to unmodeled terms: Estimated divergence of the error field (without diffusion)
  ○ S3P - Estimating magnitude of error due to unmodeled terms: Variance of error explained by diffusion
● Supplementary 4 - T-cell B-cell dynamics
  ○ S4A - Model Selection
  ○ S4B - The T-cell, B-cell phase-portrait is robust to cell-level bootstrap resampling
  ○ S4C - The T-cell, B-cell phase-portrait is robust to patient-level bootstrap resampling
  ○ S4D - T-cell B-cell vector field without smoothing
  ○ S4E - The T-cell, B-cell phase-portrait is robust to perturbation of the death estimate
  ○ S4F - Ki67 threshold determines fraction of dividing cell
  ○ S4G - The T-cell, B-cell phase-portrait estimate is robust to choice of Ki67 threshold and neighborhood size
  ○ S4H - T & B cell dynamics are consistent when modeled together with various cell types and under various levels of the 3rd cell
  ○ S4I - T & B cell Dynamics are conserved across various patient subgroups
  ○ S4J - T- and B- cell dynamics across breast cancer subtypes
  ○ S4K - FM dynamics are reproduced as components of a 6D model fit to all cells in the data
  ○ S4L - The T- and B-cell phase-portrait is reproduced as a component of a full 6D model with fibroblasts, macrophages, tumor cells, endothelial cells, T- and B-cells
  ○ S4M - Estimating magnitude of error due to unmodeled terms: Empirical position of tissues in state-space
  ○ S4N - Estimating magnitude of error due to unmodeled terms: Estimated density of tissues across the state-space
  ○ S4O - Estimating magnitude of error due to unmodeled terms: Estimated divergence of the error field (without diffusion)
  ○ S4P - Estimating magnitude of error due to unmodeled terms: Variance of error explained by diffusion
● Supplementary 5 - OSDR predicts response to chemotherapy and immunotherapy
  ○ S5A - Prediction is robust to patient bootstrap resampling
  ○ S5B - Early-treatment tumor cell average division rates are similar between responder and non-responder treatment groups
  ○ S5C - division rates change markedly over the course of treatment
  ○ S5D - Within and between patient variability are on similar scales

## Supplementary 1 - Method

### S1A - Empirical Ki67 distributions have right tails

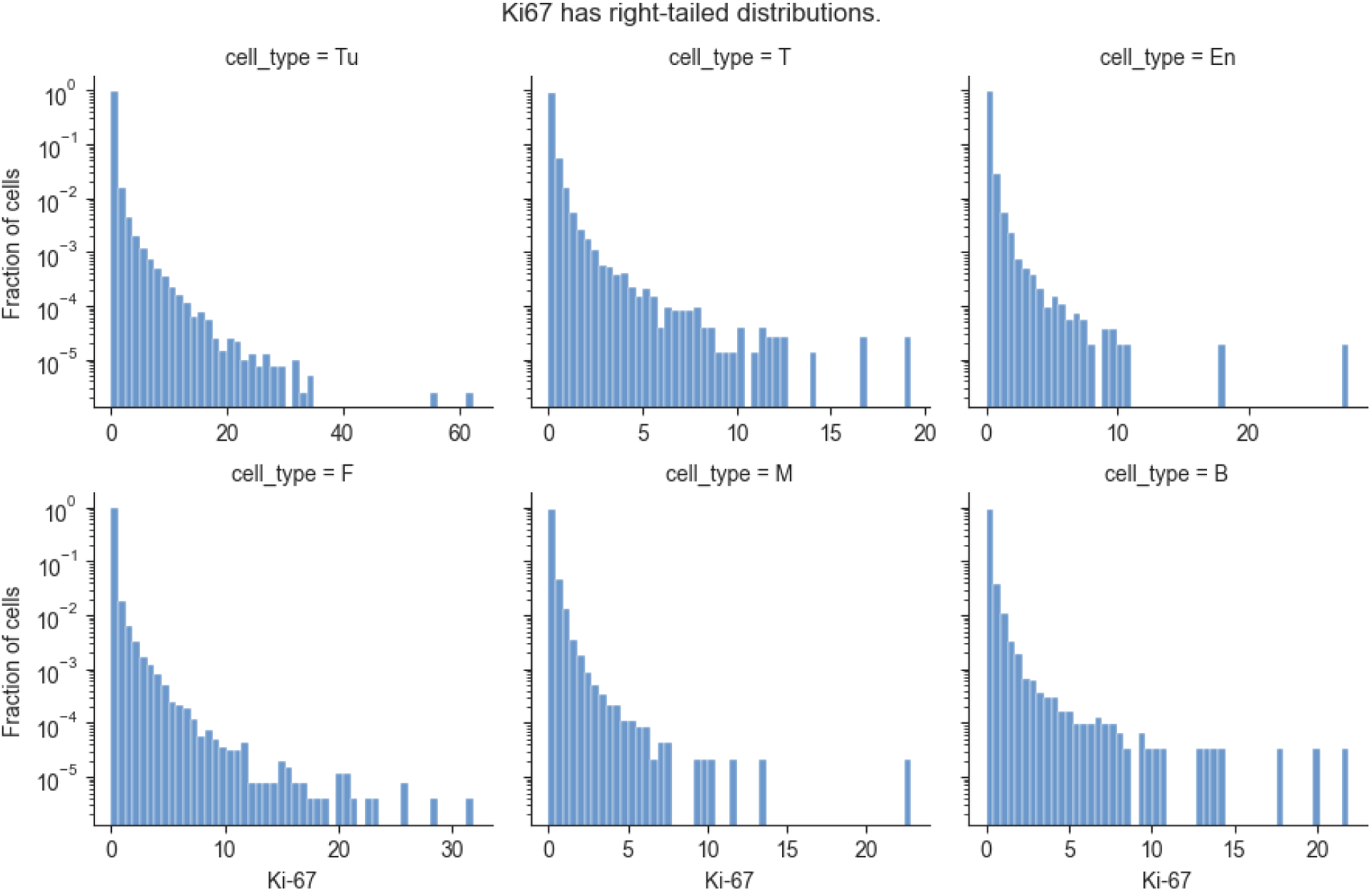

### S1B - Varying fractions of cell with Ki67 > 0 or Ki67 > noise among different cell types

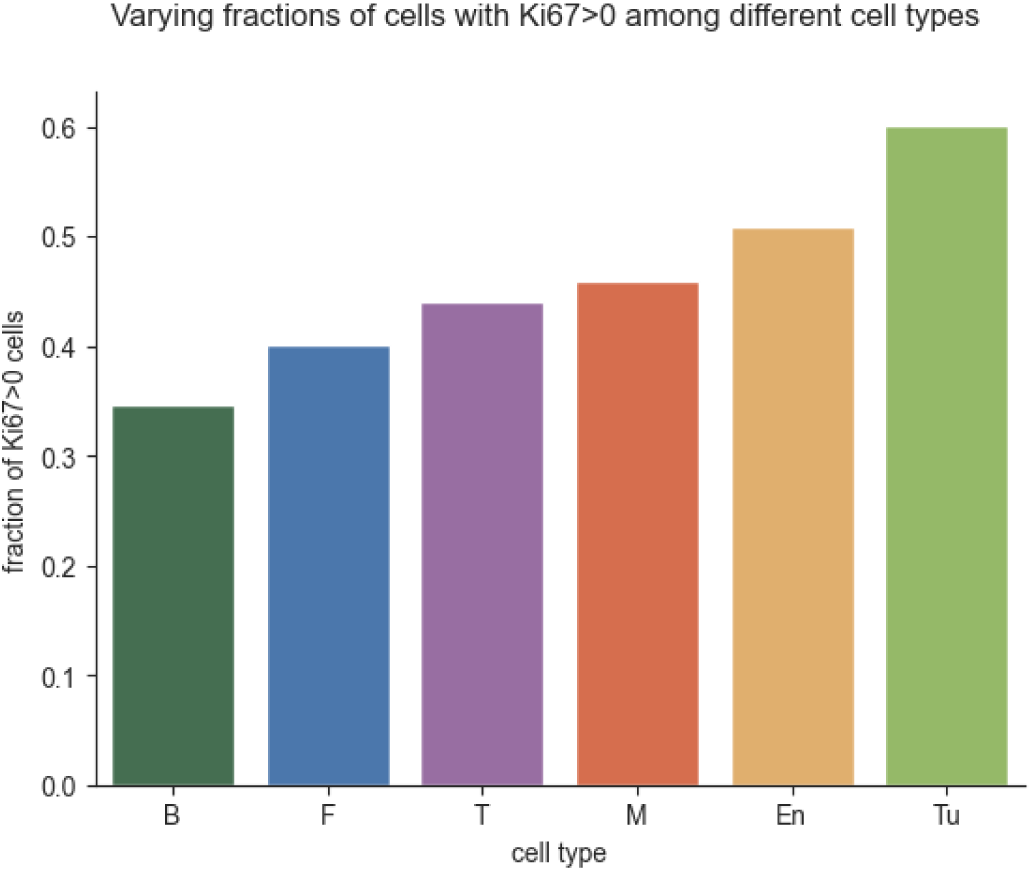
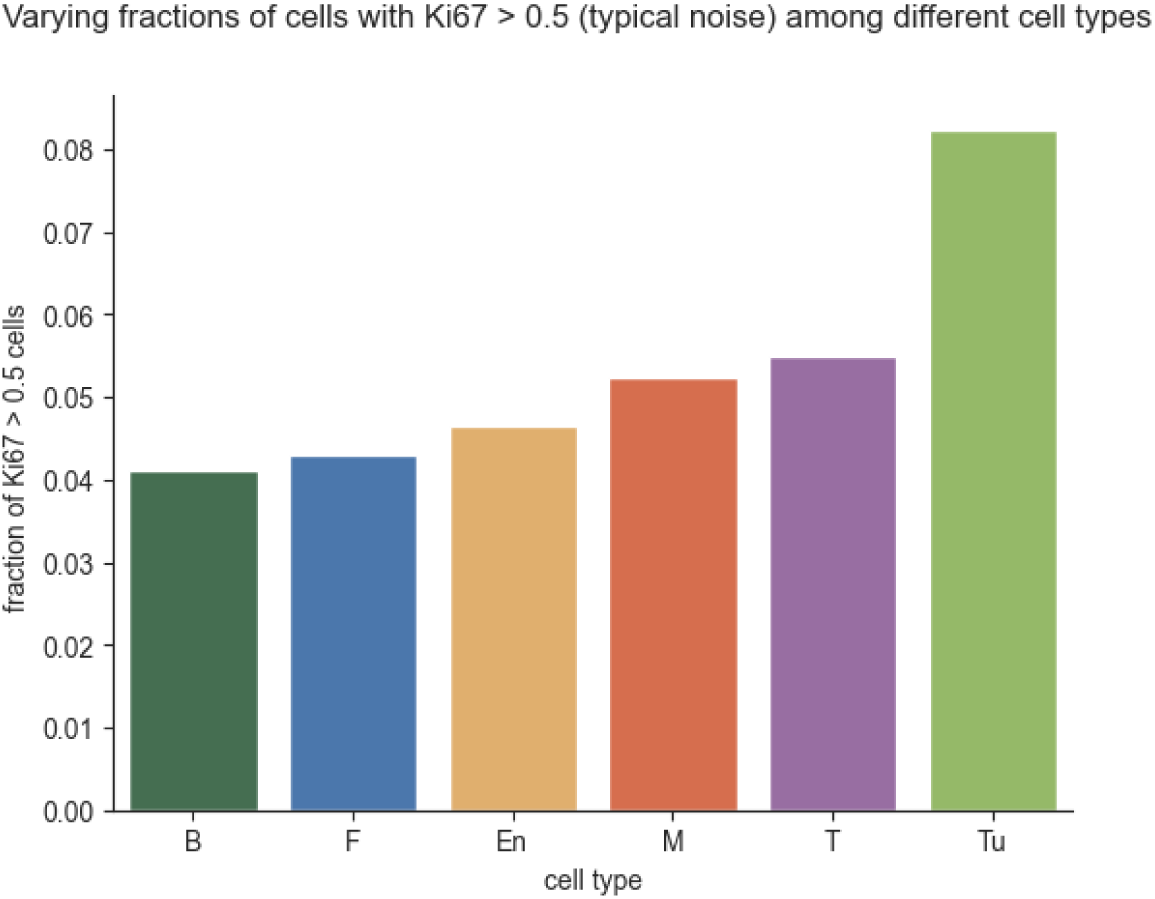

### S1C - Typical magnitude of noise of IMC protein markers

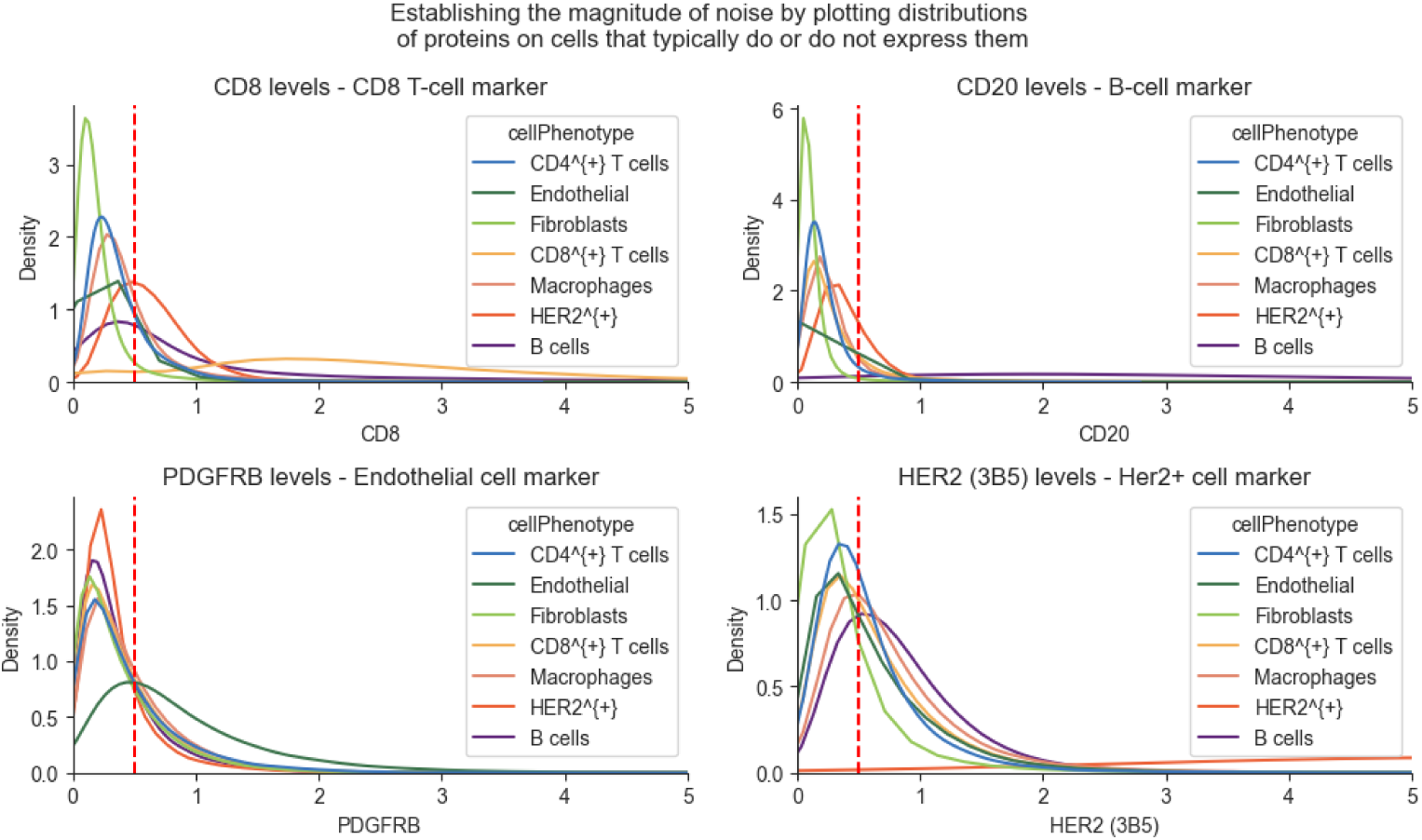

### S1D - Adjusted Ki67 distributions have similar shapes

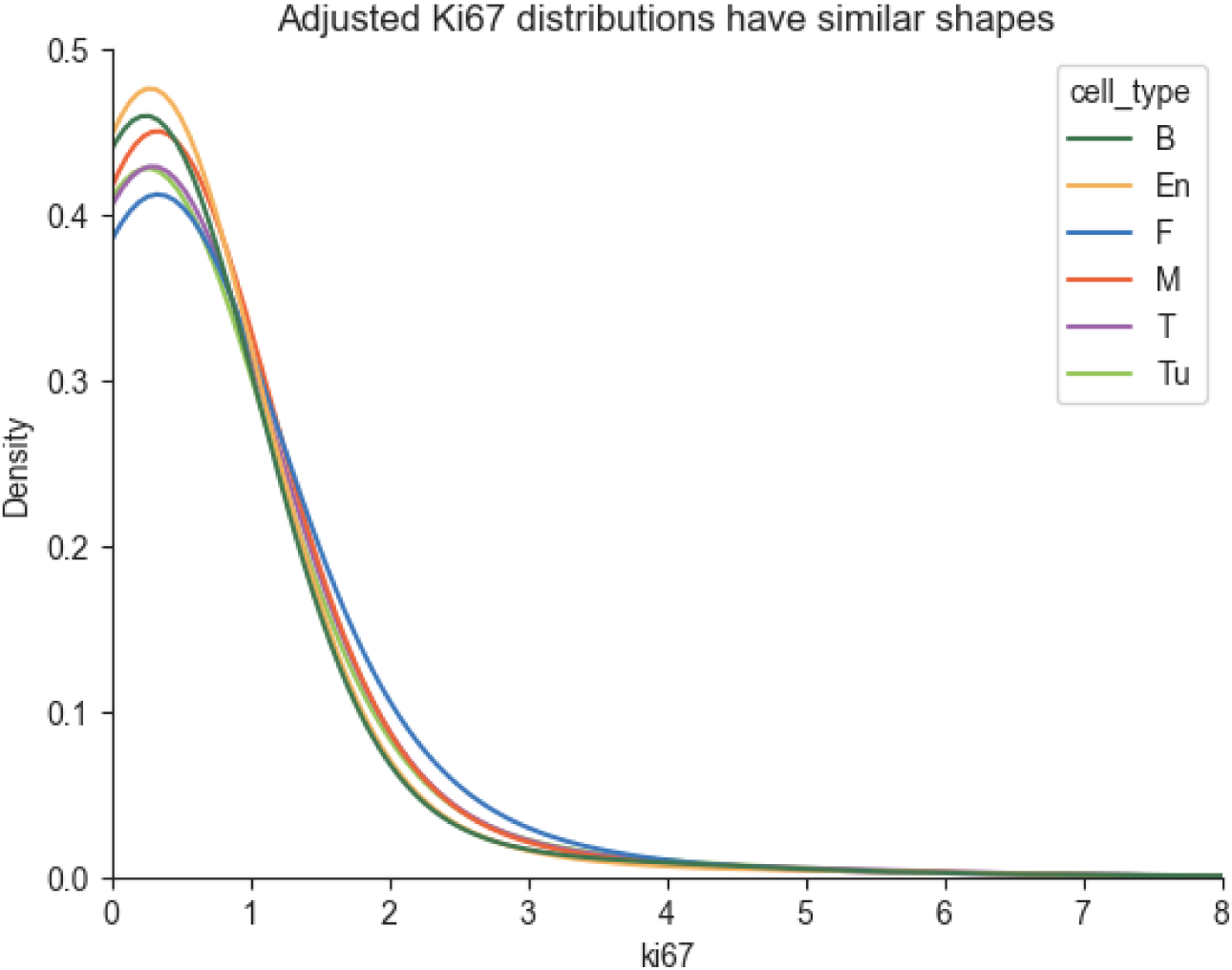

### S1E - Fraction of dividing cells (adjusted Ki67 > threshold of 0.5)

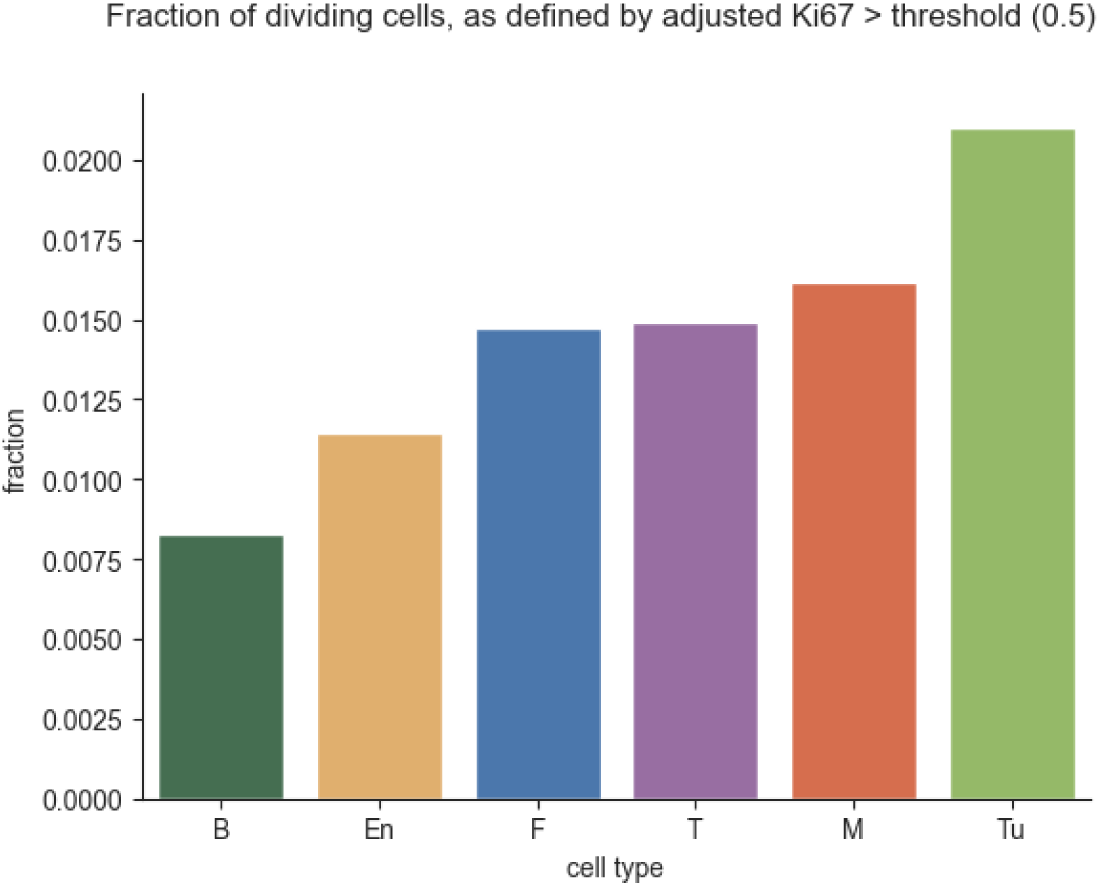

### S1F - maximal density correction power-term n determines steepness

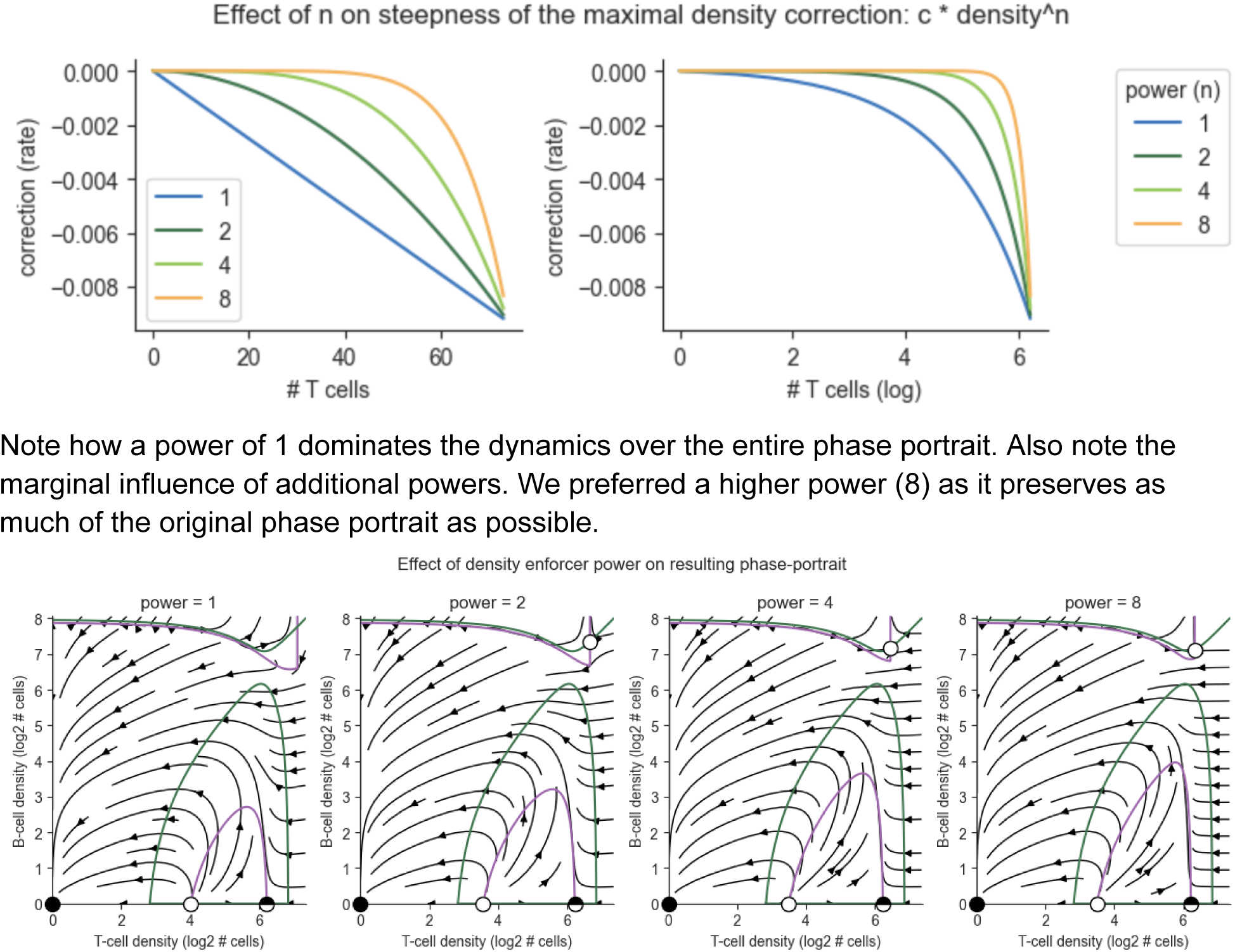

### S1G - maximal density correction in the TB model - separating the original and correction components of the phase portrait (power n=8)

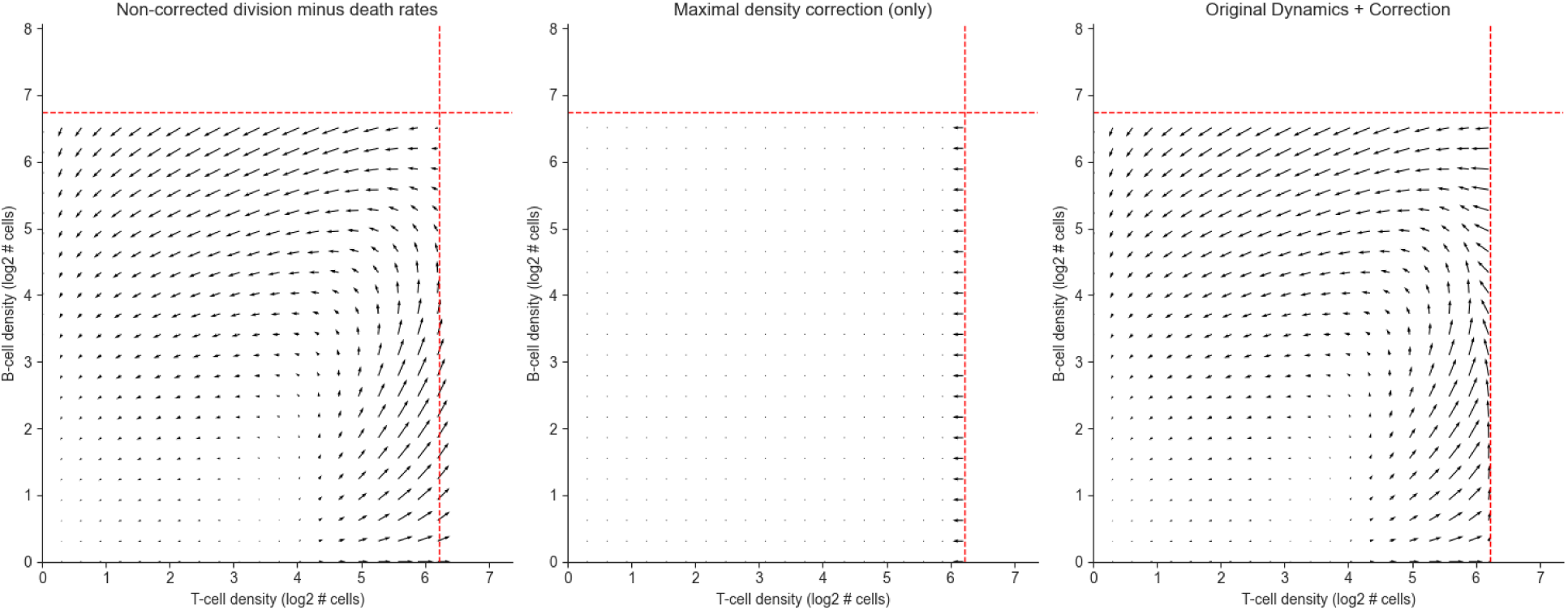

#### Population vs Neighborhood Dynamics

The following sections use the fibroblast-macrophage model fit to the dataset by (Fig 3, Danenberg et al. 2022).

### S1H - Population-dynamics produce a sequence of tissues by sampling division and death events for cells in the tissue

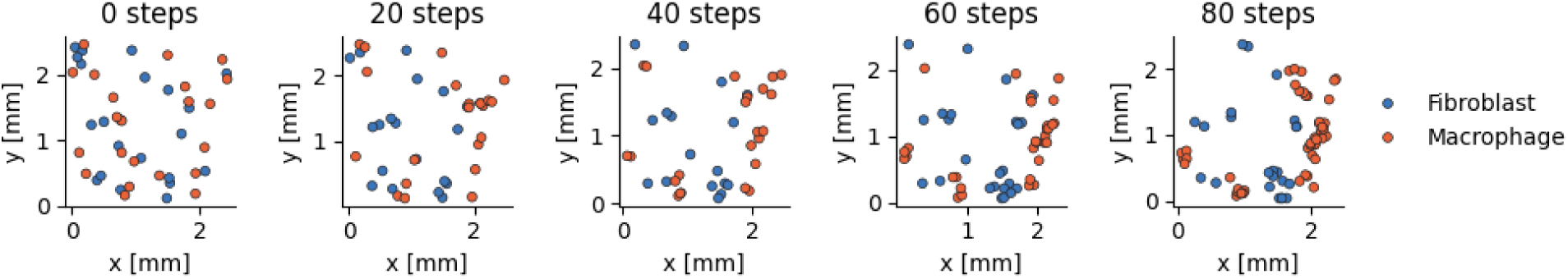

### S1I - A high diffusion coefficient can disperse cells out of tissue boundaries

Here we set the diffusion coefficient to 100 microns. That is, we apply a 2D gaussian translation to each cell, where the standard deviation in each axis is 100 microns.

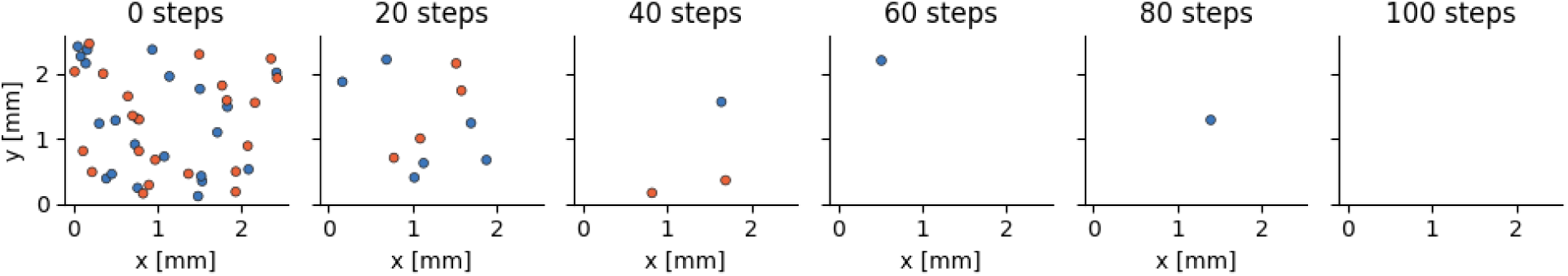

### S1J - A low diffusion coefficient can create “isolated” regions in the tissue

Here we set the diffusion coefficient to zero. In some regions all cells died, creating blank patches. The tissue appears as isolated clusters of cells. Note there is still some spatial dispersion because new cells are placed in a random location within the dividing cell’s neighborhood.

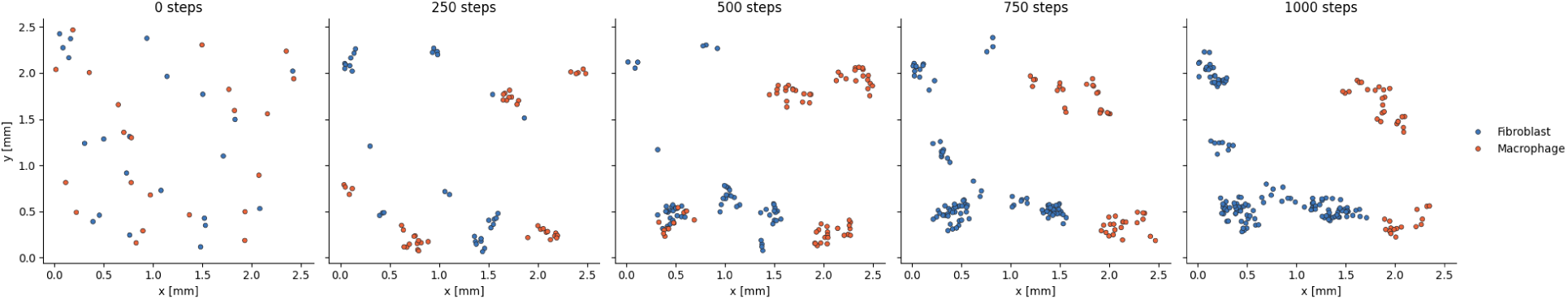

### S1K - Example where average cell-density underestimates the stable fixed-point, while the mode is exact

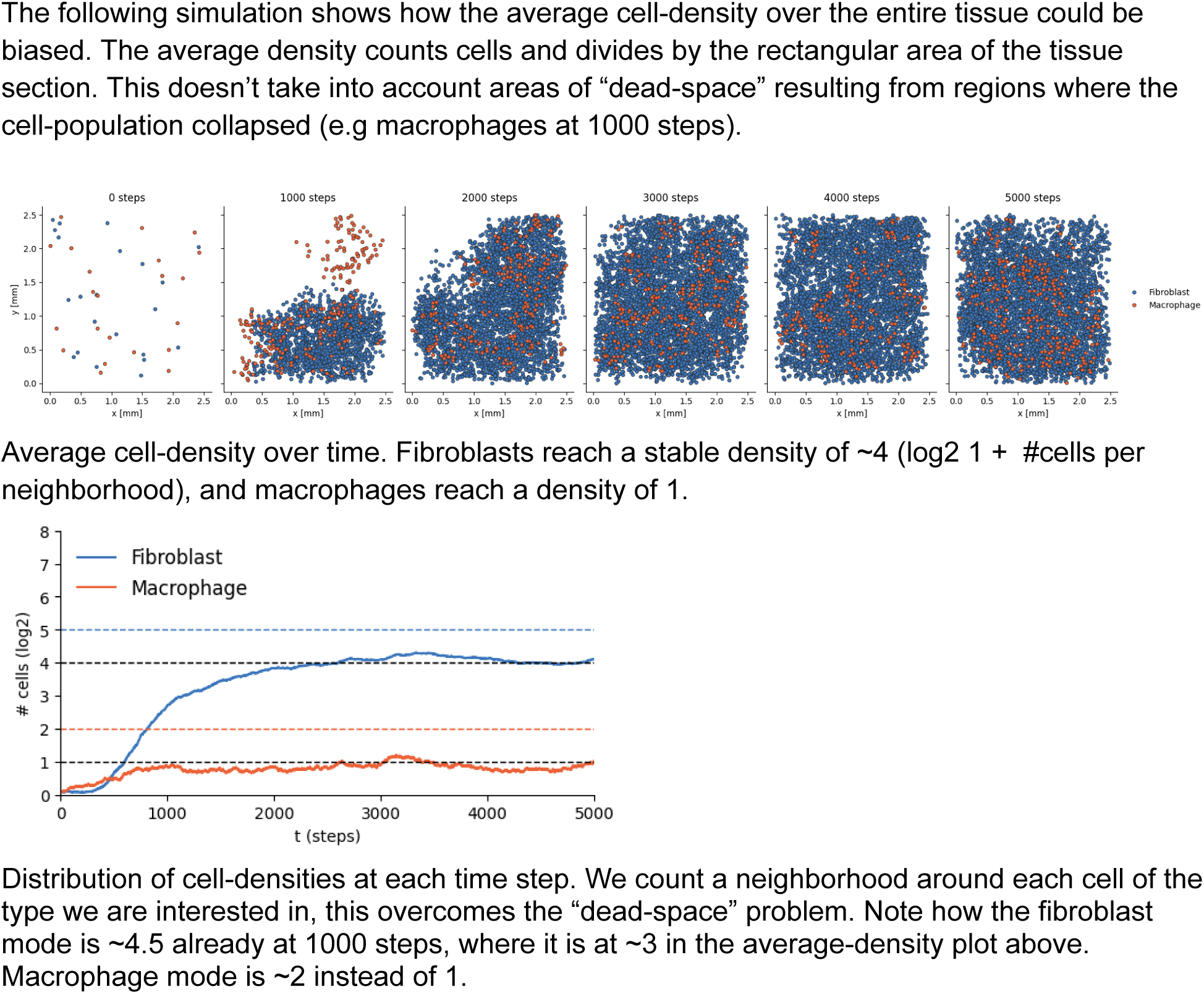
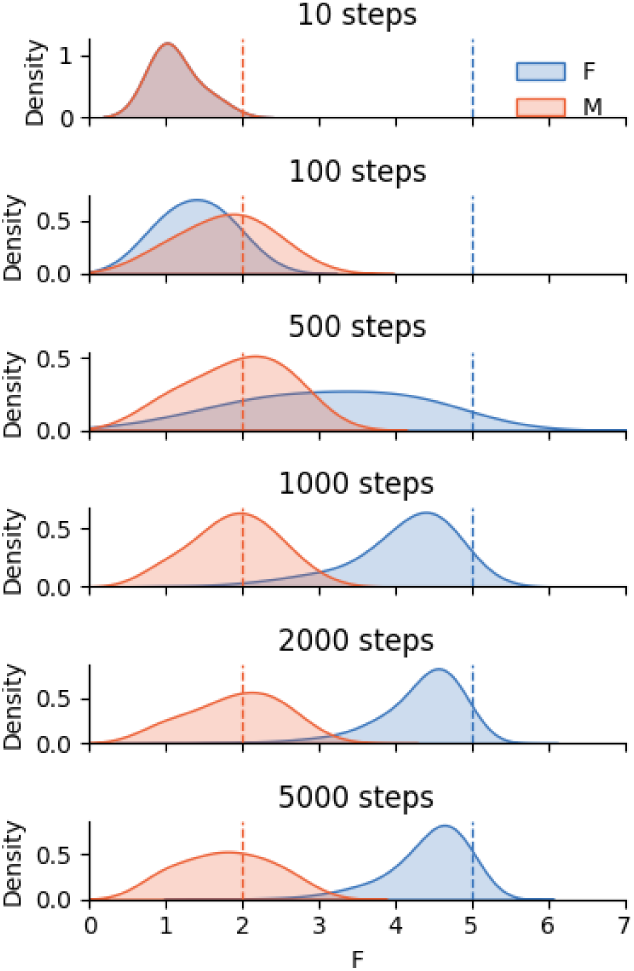

### S1L - Collapse of cell populations in the stochastic population-dynamics model can produce large discrepancies between neighborhood and population dynamics

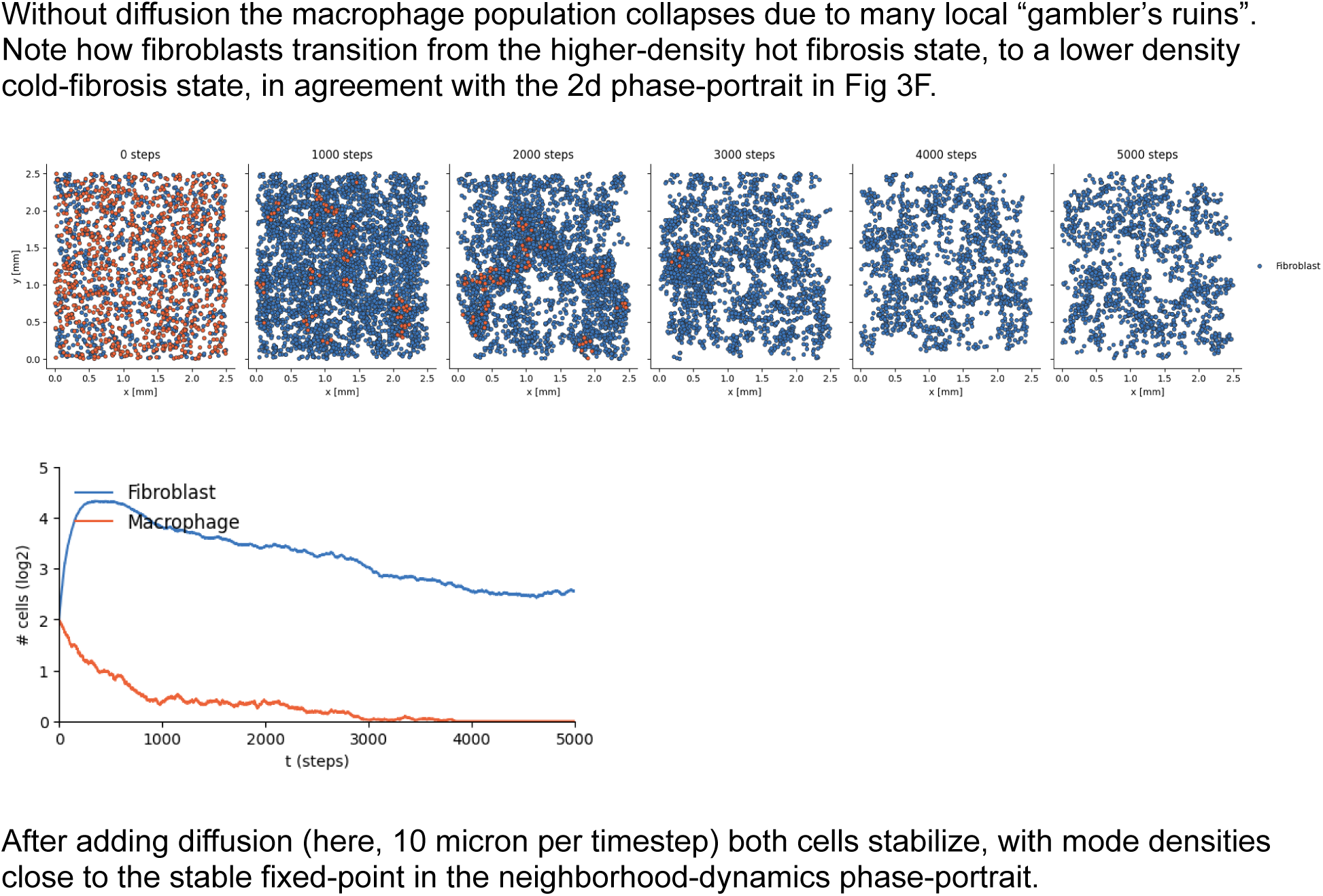
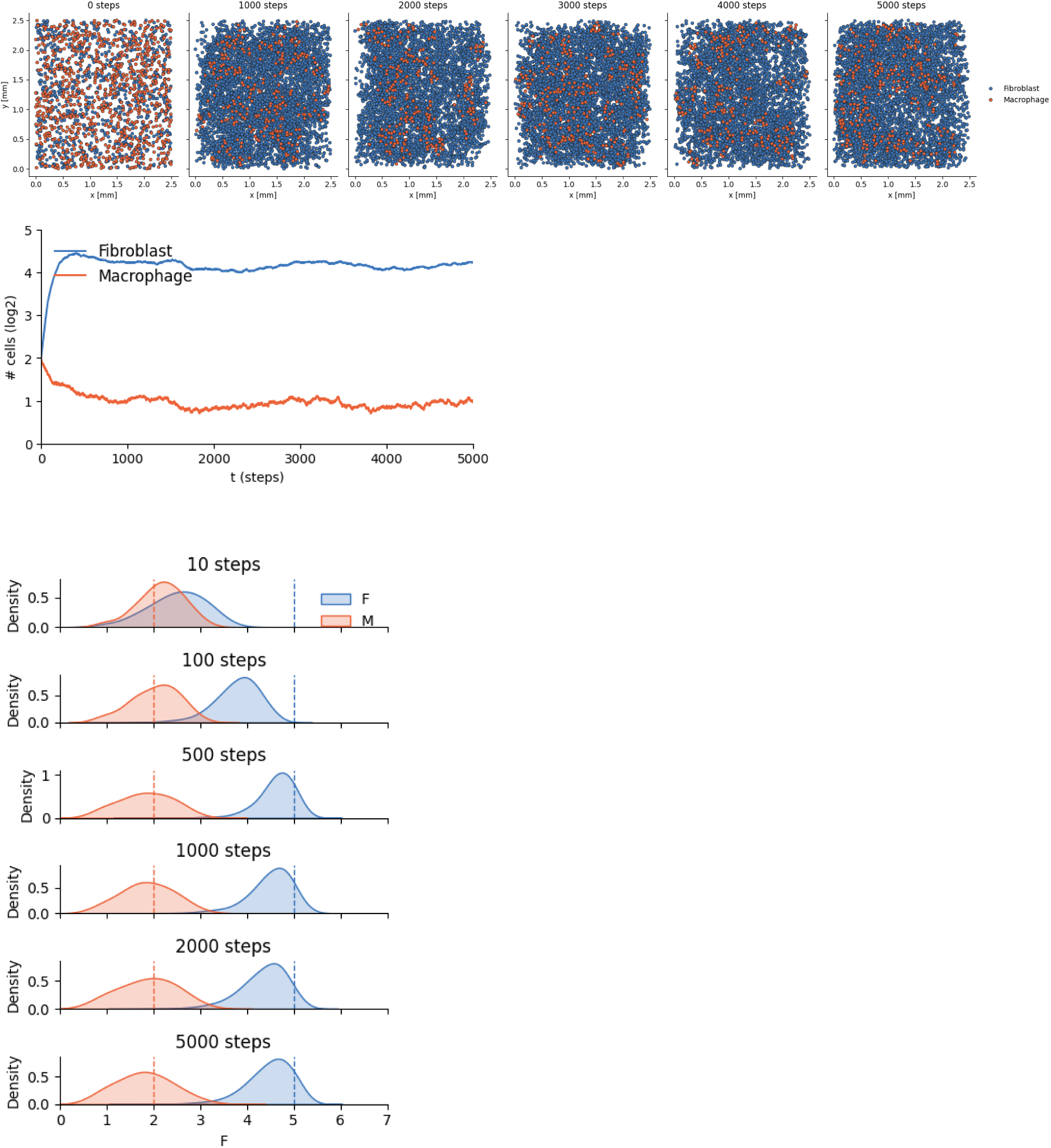

### S1M - Certain initial tissue configurations can produce significantly different trajectories

The TB circuit from figure 4D produces a large flare starting from a random placement of cells in a density corresponding with (T=4.5,B=1) in the phase-portrait.

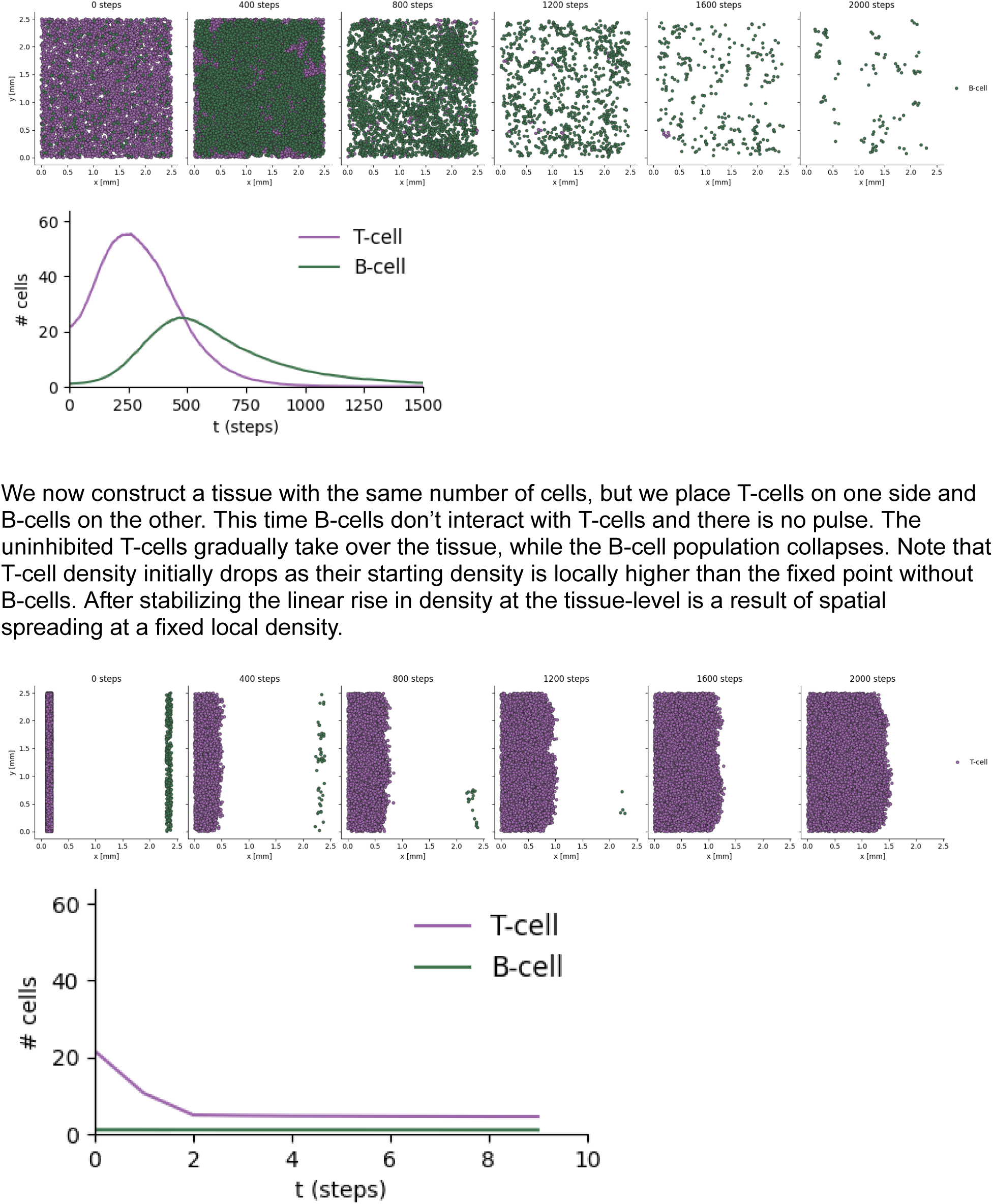
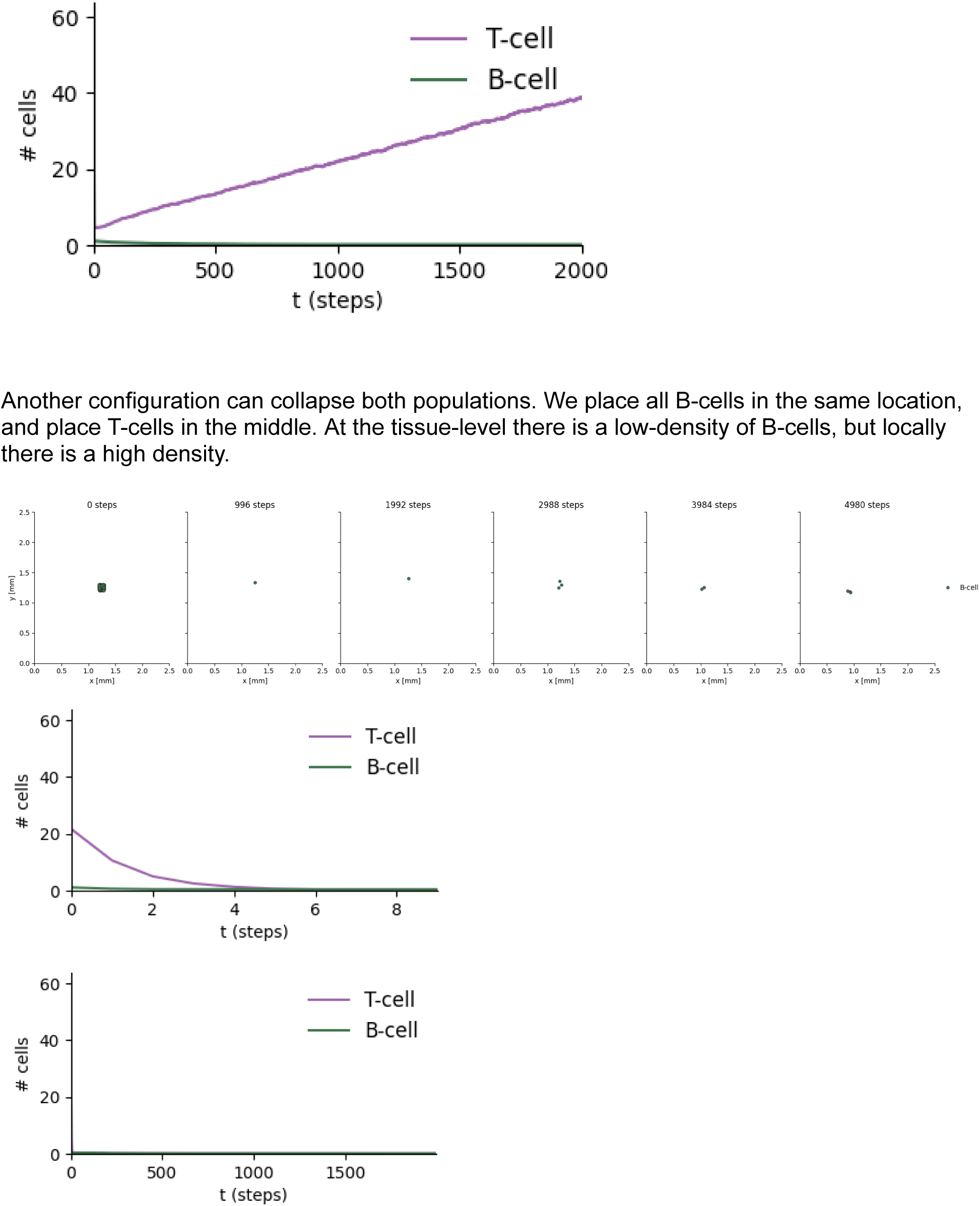

## Supplementary 2 - Model Estimation

### S2A - cell-type definitions used in each dataset

#### Dataset by Danenberg et. al

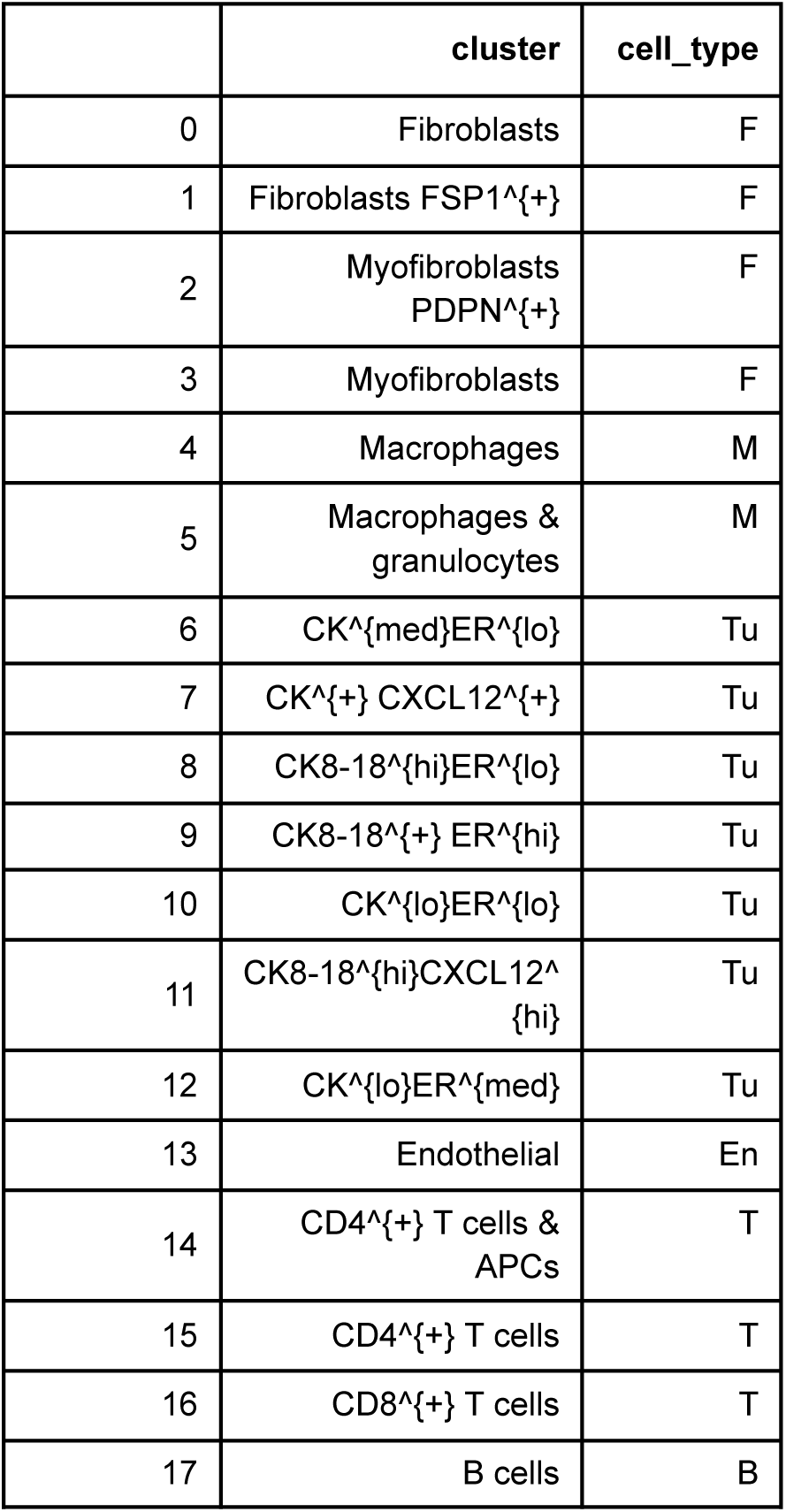

#### Dataset by Fischer et. al (cluster mapping to cell type)

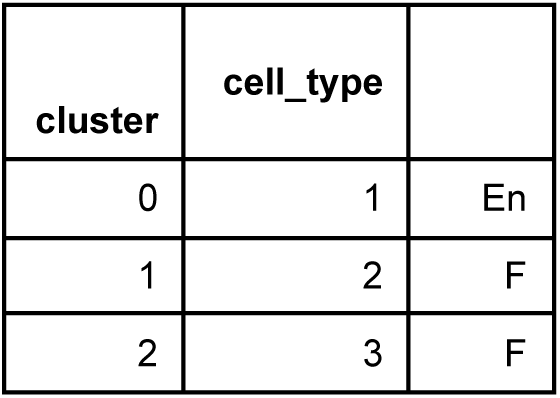

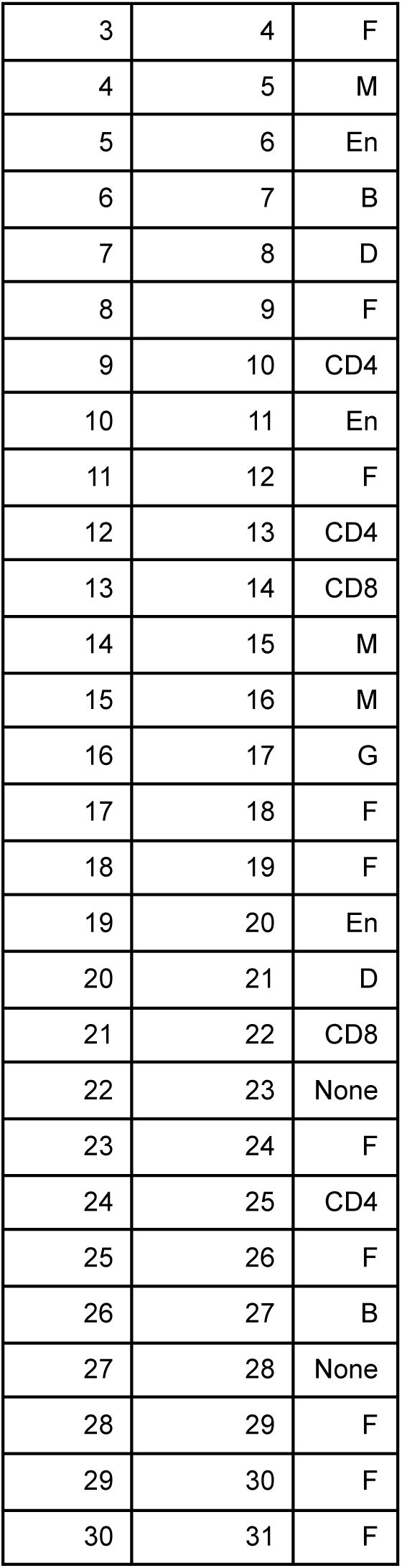

#### Dataset by <u>{Updating}</u>

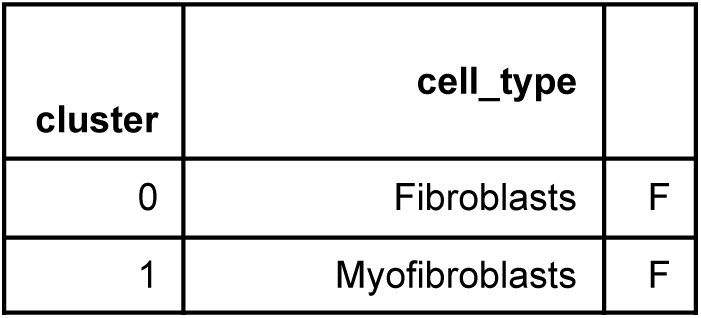

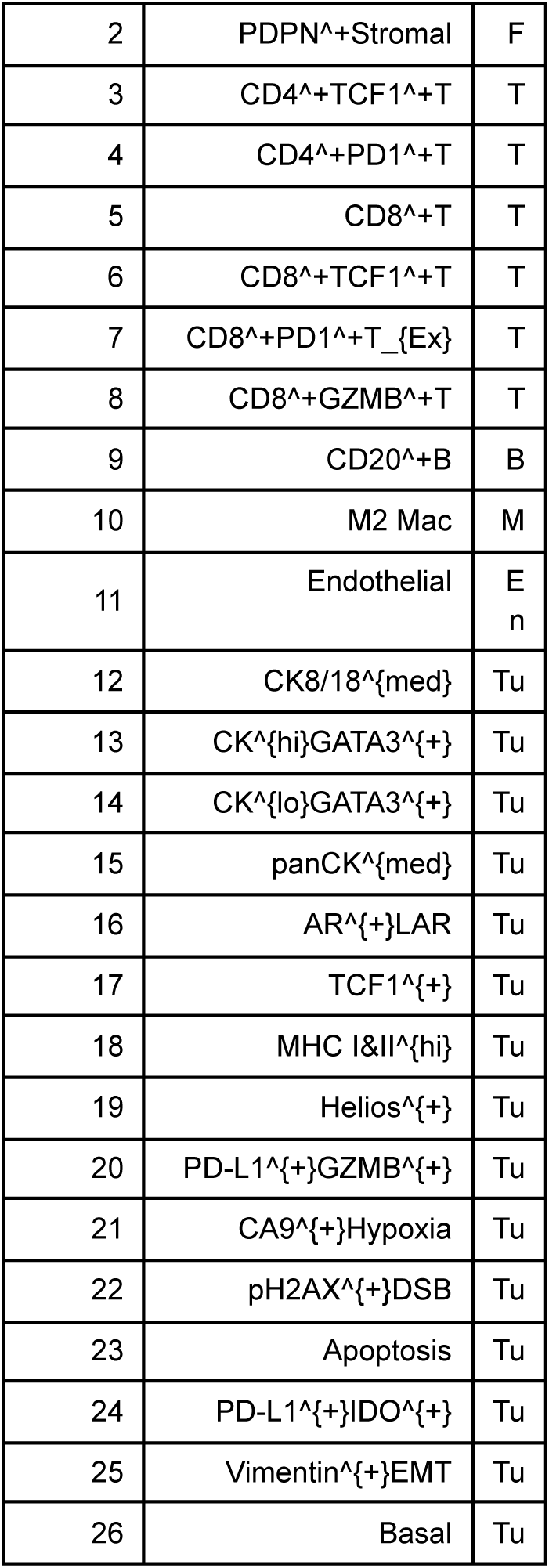

### S2B - Calibration plots for all cell types, compared with calibration on same data with permuted divisions

Each green dot aggregates 4000 cells, red lines are min & max true probabilities. The x-axis location of a dot is the model’s average predicted probability for a cell in that subset. The y-axis location is determined by the fraction of cells that are truly dividing, as determined by a Ki67 measurement > threshold.

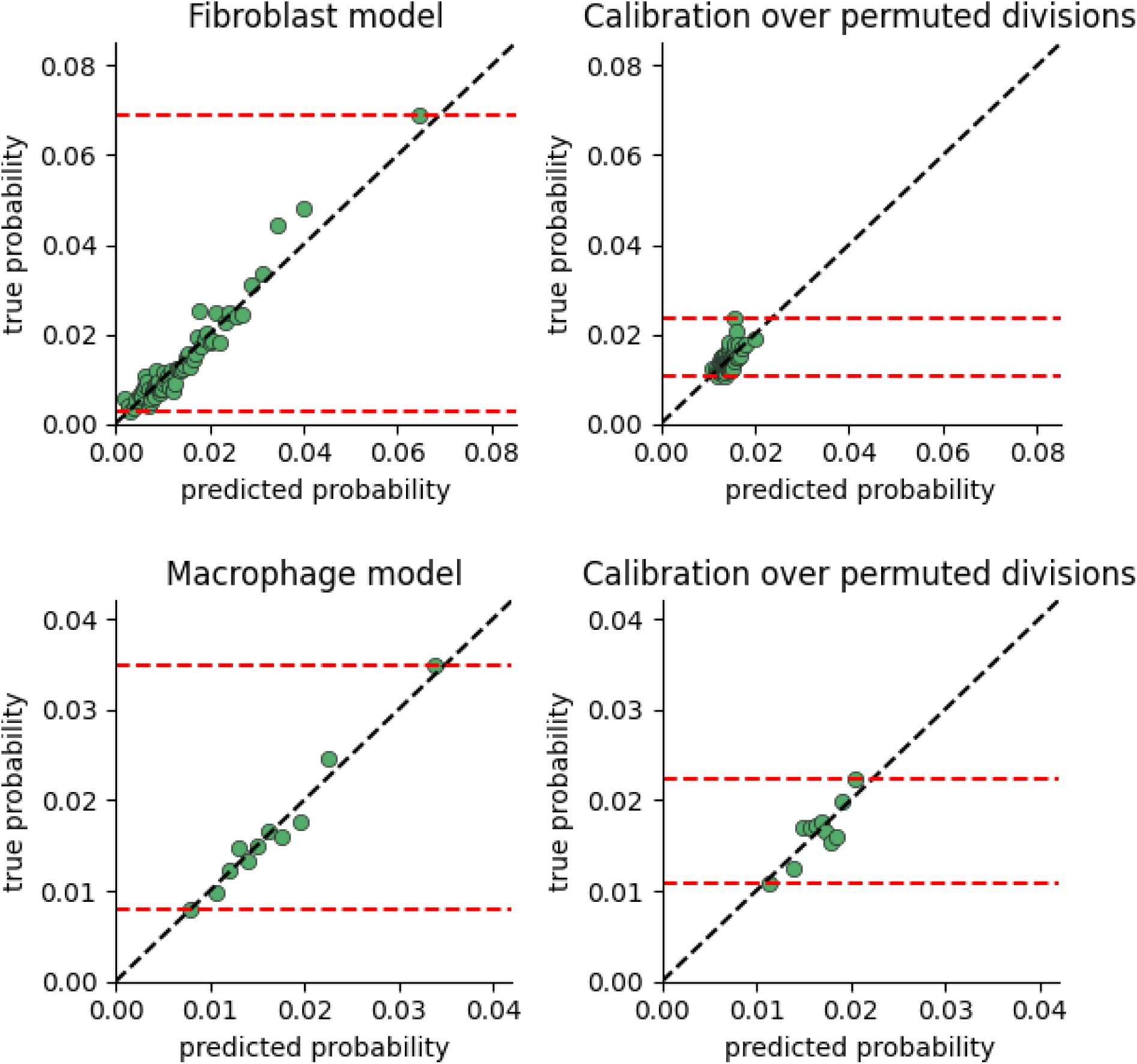
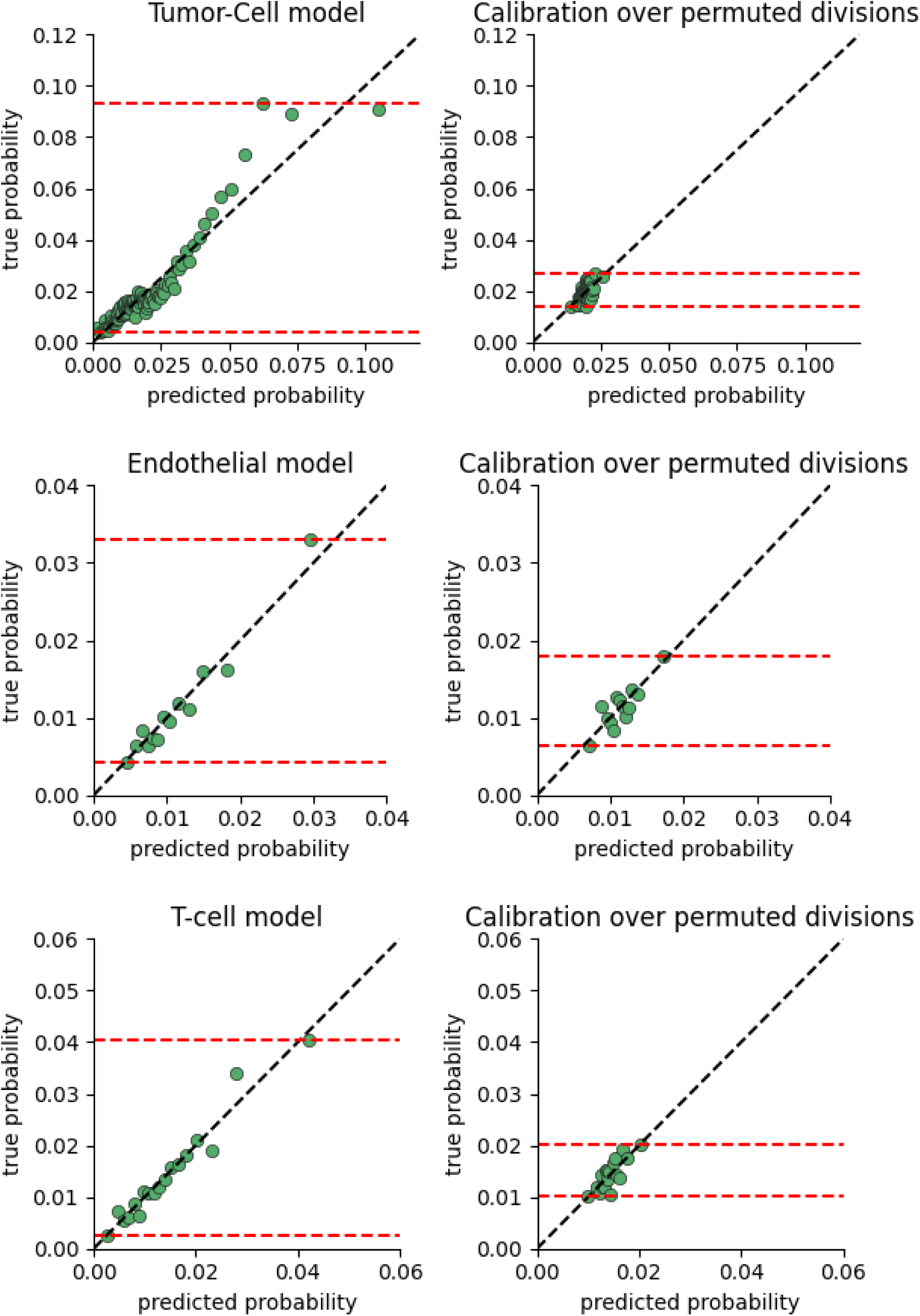
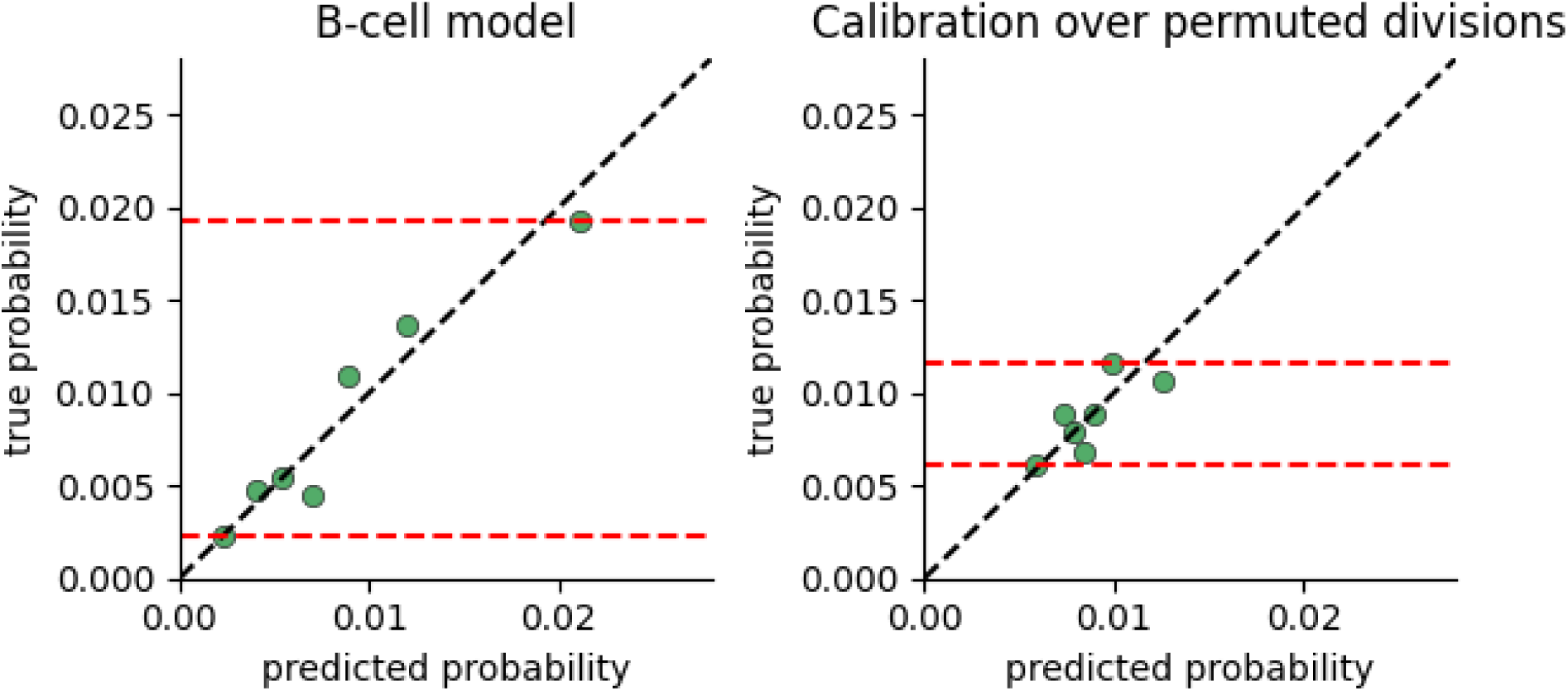

### S2C - Logistic regression model log-likelihood ratio p-values and example model summary

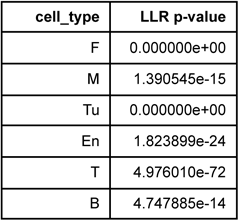

### S2D - Example summary of model fit

The variable for each cell type is log(1 + cell count). We include up to 2nd order interactions (i.e all 2nd order polynomial terms are included). We present the fibroblast proliferation model fit summary below.

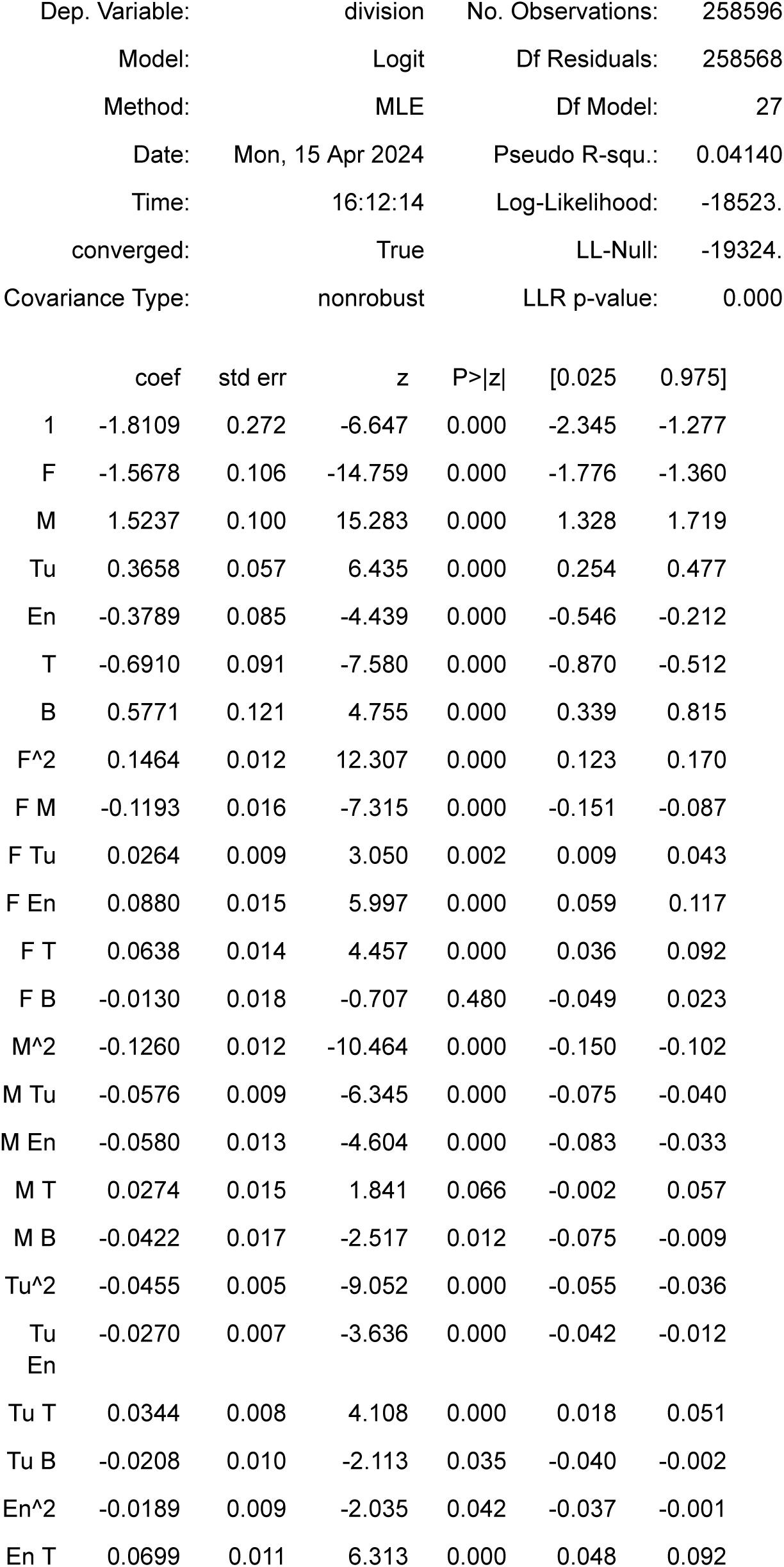
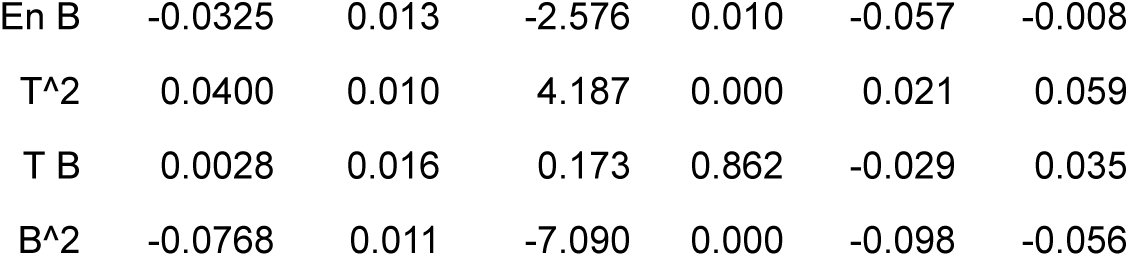

### S2E - model performance as a function of neighborhood radius

To test at which radius neighbors become less predictive we fit 6D models to the 6 main cell-types in the dataset by Danenberg et. al. We fit models to neighborhood sizes in 10 micron increments across the published range of interactions (Oyler, Yaniv 2017). We plotted the log-loss (negative log-likelihood, lower is more predictive) for each cell-type and radius (figure below). We find that neighborhood radius has a negligible effect on the loss. Also, estimated phase-portraits maintained their topology under a wide range of neighborhood radii (S3E, S4F). We conclude that the statistical relationship between neighborhood composition and division rate, as well as the derived dynamics, are robust to choice of neighborhood radius. This could be due to correlation between nearby neighborhood compositions.

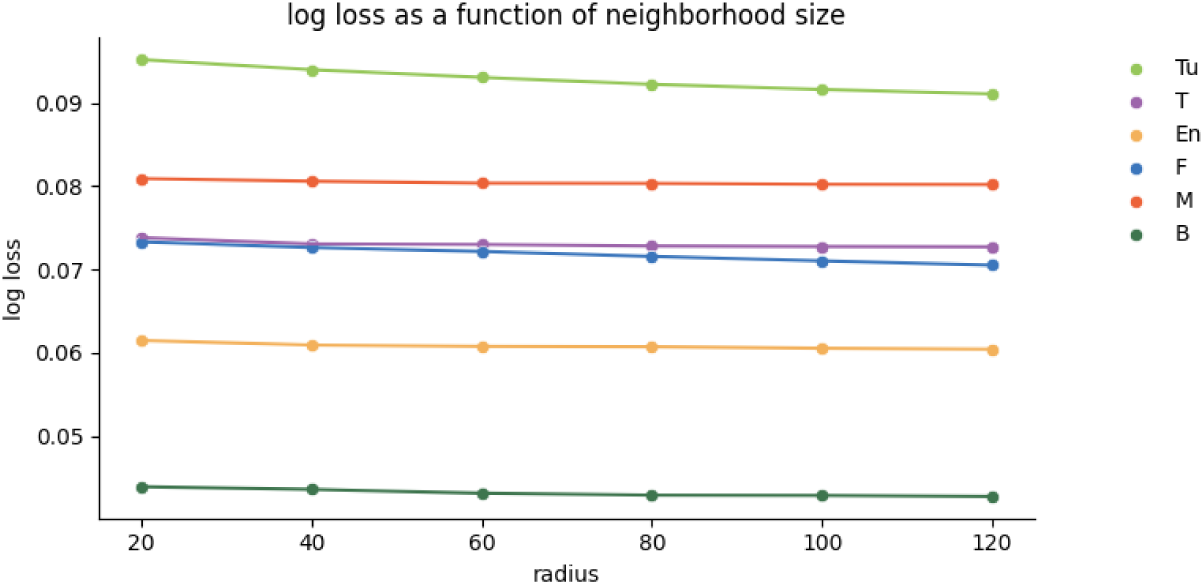

### S2F - The ability of current neighborhood to predict gradients and growth factors that determine proliferation, based on separation of timescales between signaling and proliferation dynamics

The features used in OSDR are based on counts of neighboring cells. These features are empirically predictive of cell division rates in the breast cancer cohorts (Fig3A, S2A). Here we provide some background for this choice of features, as well as possible settings where the current tissue composition might not be predictive of cell division rates.

Cells modulate the proliferation of nearby cells through signaling proteins such as cytokines and growth factors. The half-life of a signaling protein determines the time it takes to reach a steady state concentration. For instance, most cytokines are known to have a short half-life, on the order of minutes to hours determined in many cases by endocytosis in receiver cells (Adler 2018; Oyler-Yaniv et al. 2017).

On the other hand, the timescale of changes in tissue composition is typically much slower. For example, published doubling times in breast-cancer are in the range of months to years (Förnvik et al. 2016). Even in the most rapidly developing breast cancers, tumor mass only increases by 1% per day. In such systems we can assume that growth factor concentrations are near quasi-steady-state.

To illustrate this principle we analyze a simplified mathematical model describing a single cell type secreting a growth factor *g*. Each cell secretes the growth factor at a fixed rate α, and the growth rate is removed at a rate γ:

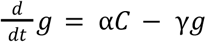

The solution shows that growth factor concentration approaches its steady state exponentially fast with a half-time of 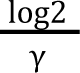 equal to the half life of the growth factor:

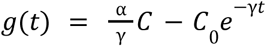

At (quasi) steady state, 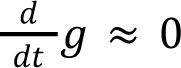 and hence the growth factor is proportional to the number secreting cells 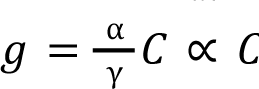. In settings where tissue composition changes very rapidly at a similar timescale to that of growth factor degradation, current tissue composition might not be predictive of cell division rates.

See (Adler 2018) for a study on more elaborate mechanisms of growth factor control.

As noted in the text, neighbor counts also reflect a cell’s chance to directly contact another, capturing processes such as contact inhibition or signaling via direct contact. Counts of some cell types might also influence factors like tissue inflammation or hypoxia, which can influence cell proliferation rates. Effects such as hypoxia in the core of a tumor can be captured by low proliferation of a cancer cell surrounded by other cancer cells. In contrast, cancer cells with non-cancer neighbours are at the edge of the tumor. Effects of inflammation may be captured by neighboring macrophages or lymphocytes, given that the range of many cytokines is limited by endocytosis and is hence on the order of 100 microns (Oyler-Yaniv et al. 2017). In effect, the present use of a neighborhood can plausibly capture spatial effects of gradients with some accuracy.

### S2G - overview of 2-cell simulations

In sections 2D,E,F,G we present results from simulations used to test feasibility of the OSDR approach and the required sample size for recovering various aspects of a ground truth model, including location and type of fixed points. The four models considered exhibit the four possible organizations of stable fixed points for two cells, such that each cell is stable either in the presence of the other cell or not.

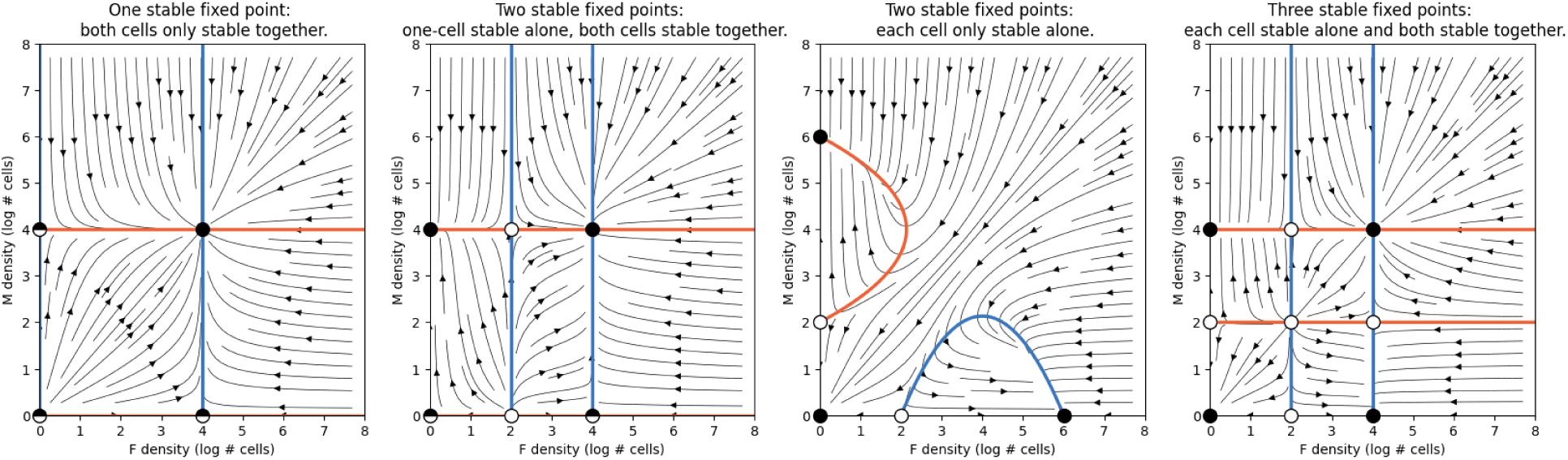

The following simulations show that some aspects are recovered with few cells (as little as 1K), such as a stable point off-axis or presence of any kind of stability on an axis. Discerning between types of stability on the axes requires more cells. At 25K cells all aspects are recovered with high accuracy.

### S2H - Simulation 1 - One fixed point

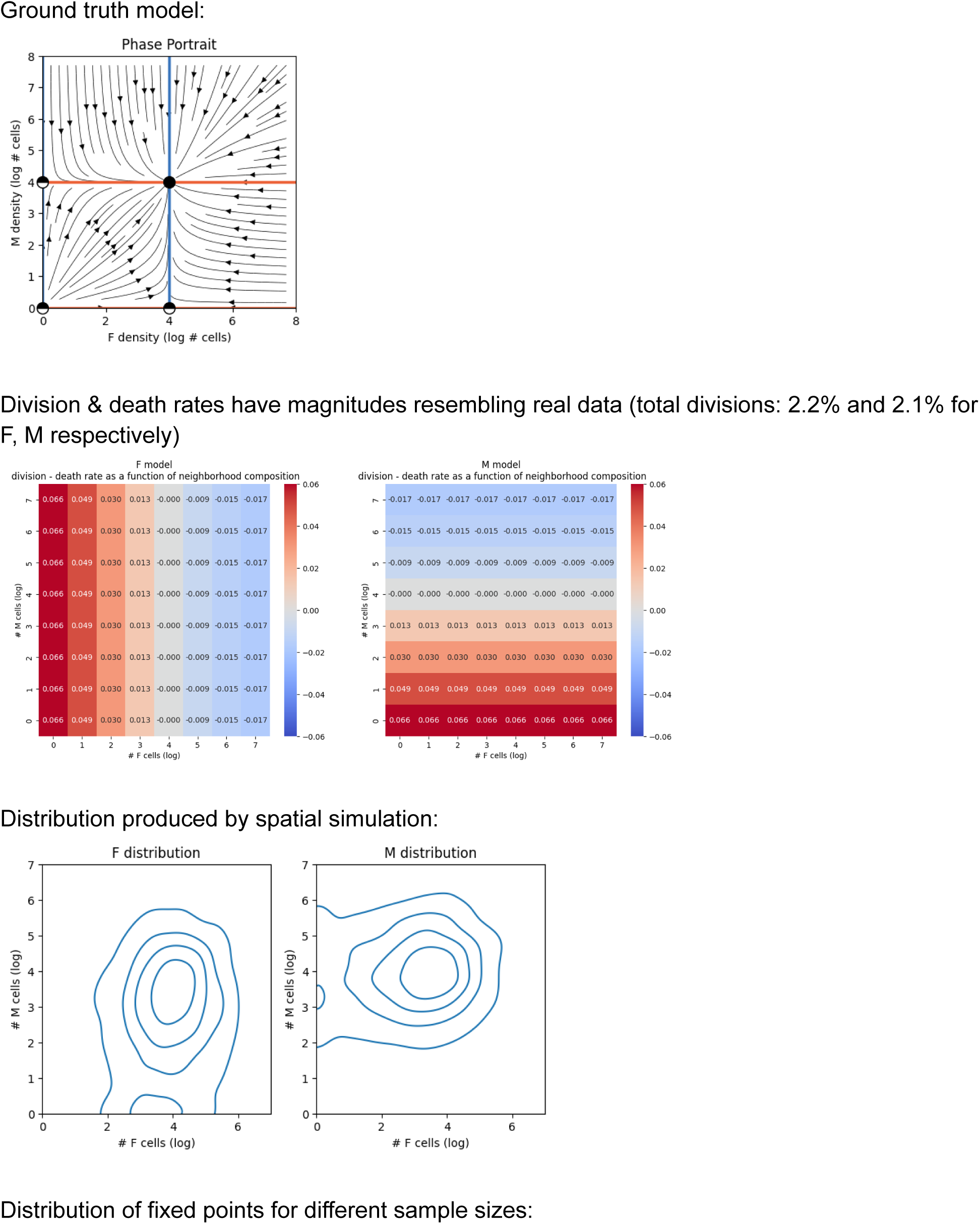
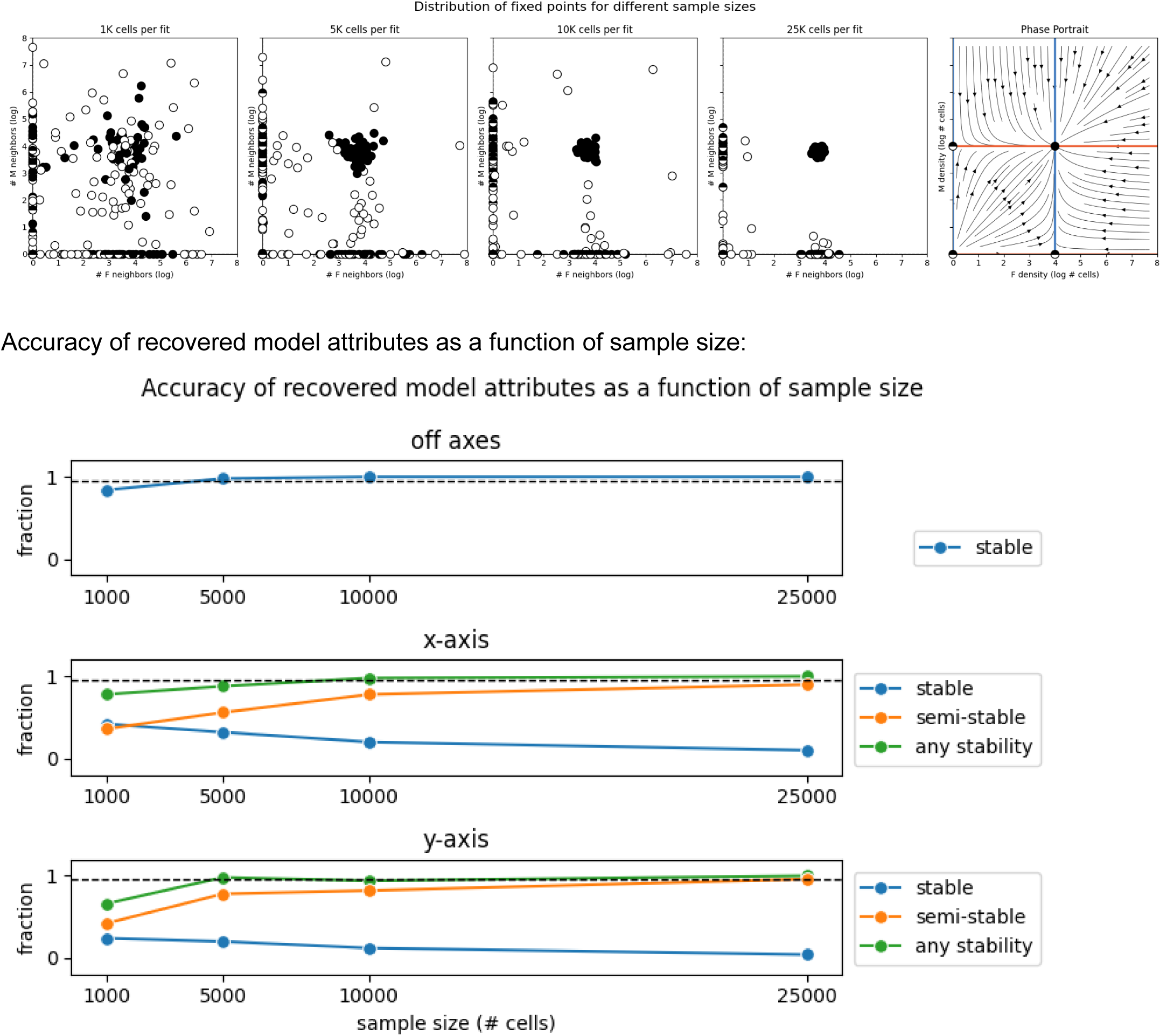

### S2I - Simulation 2 - Two fixed points

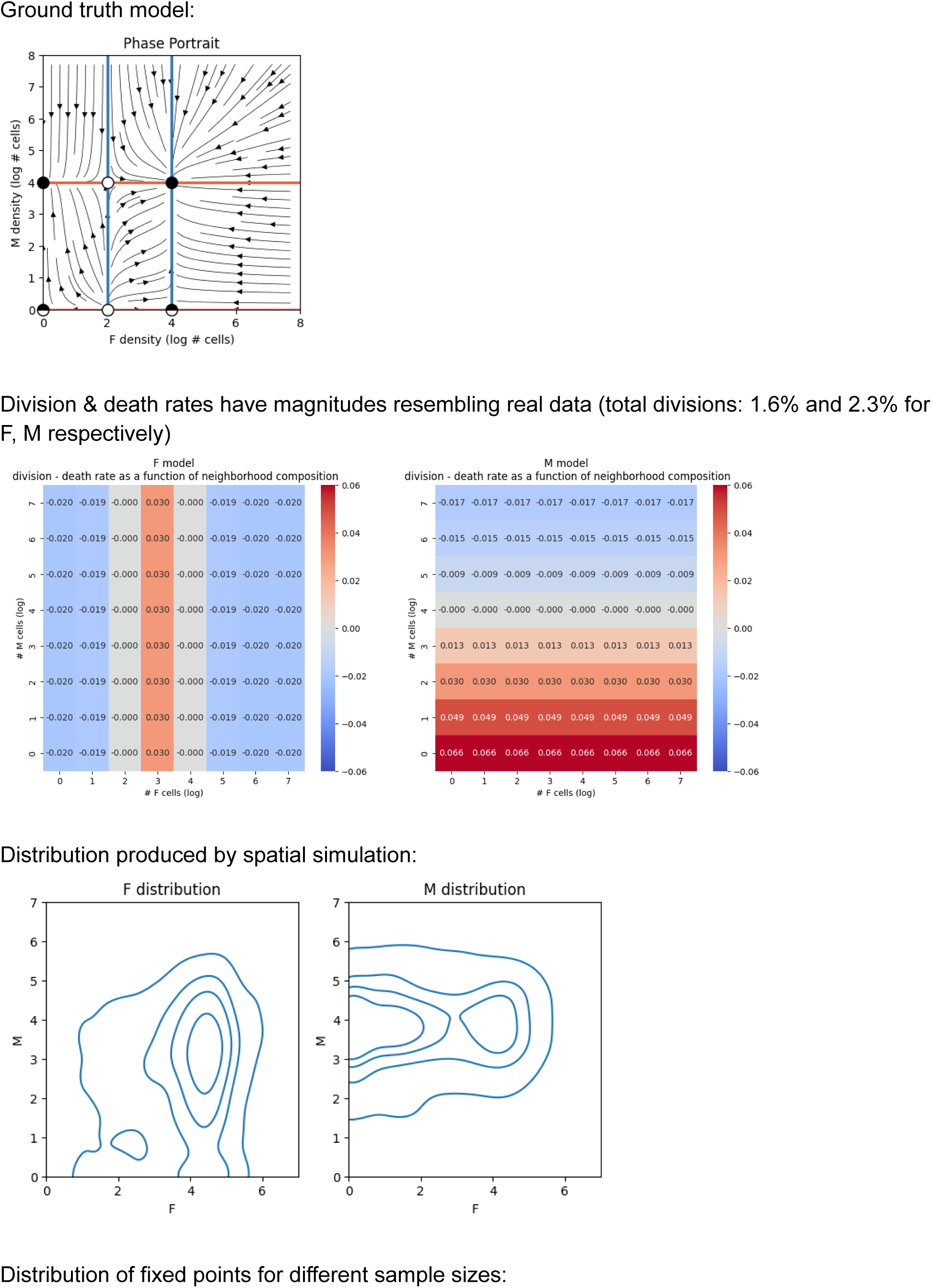
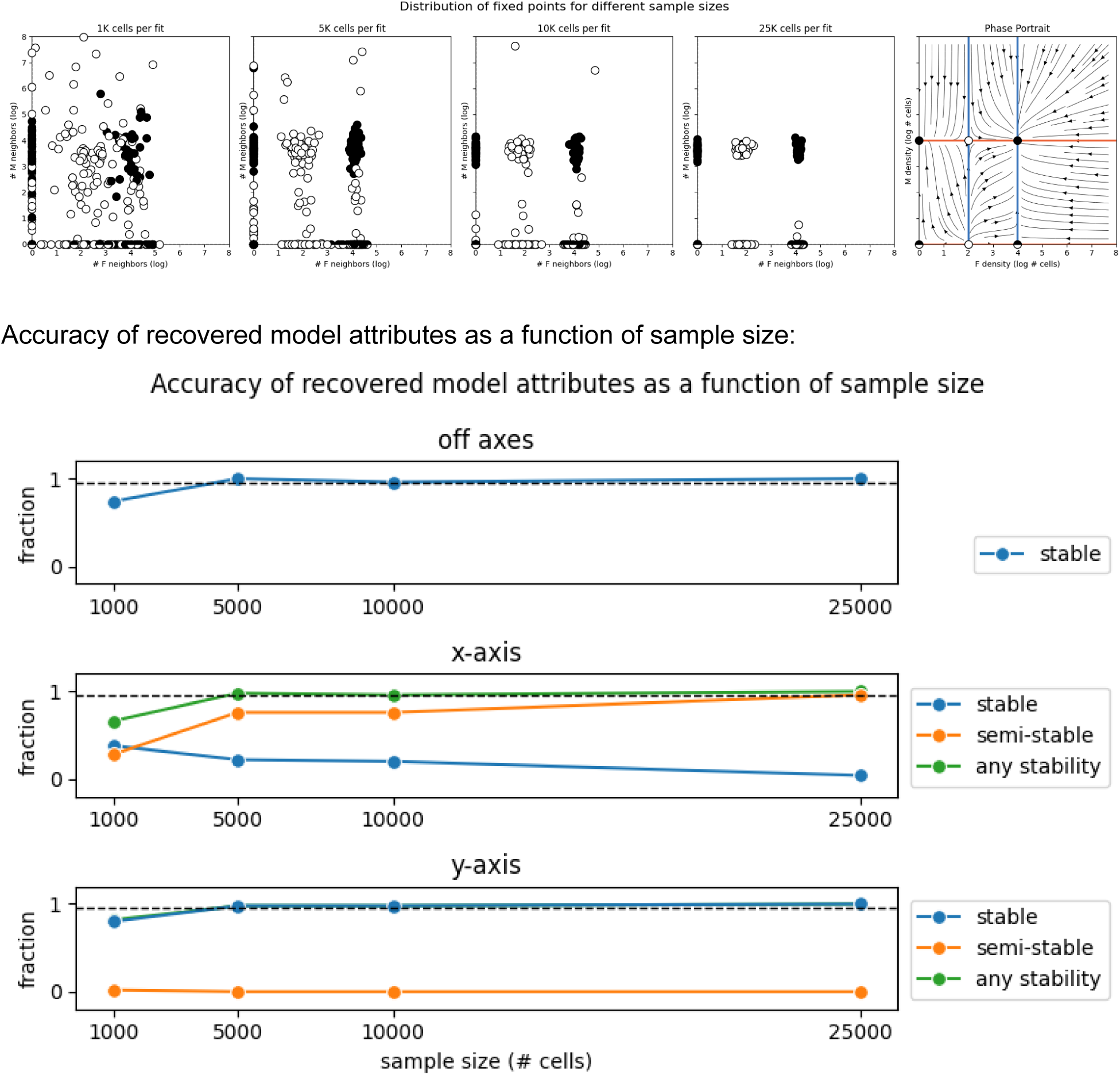

### S2J - Simulation 3 - Three fixed points

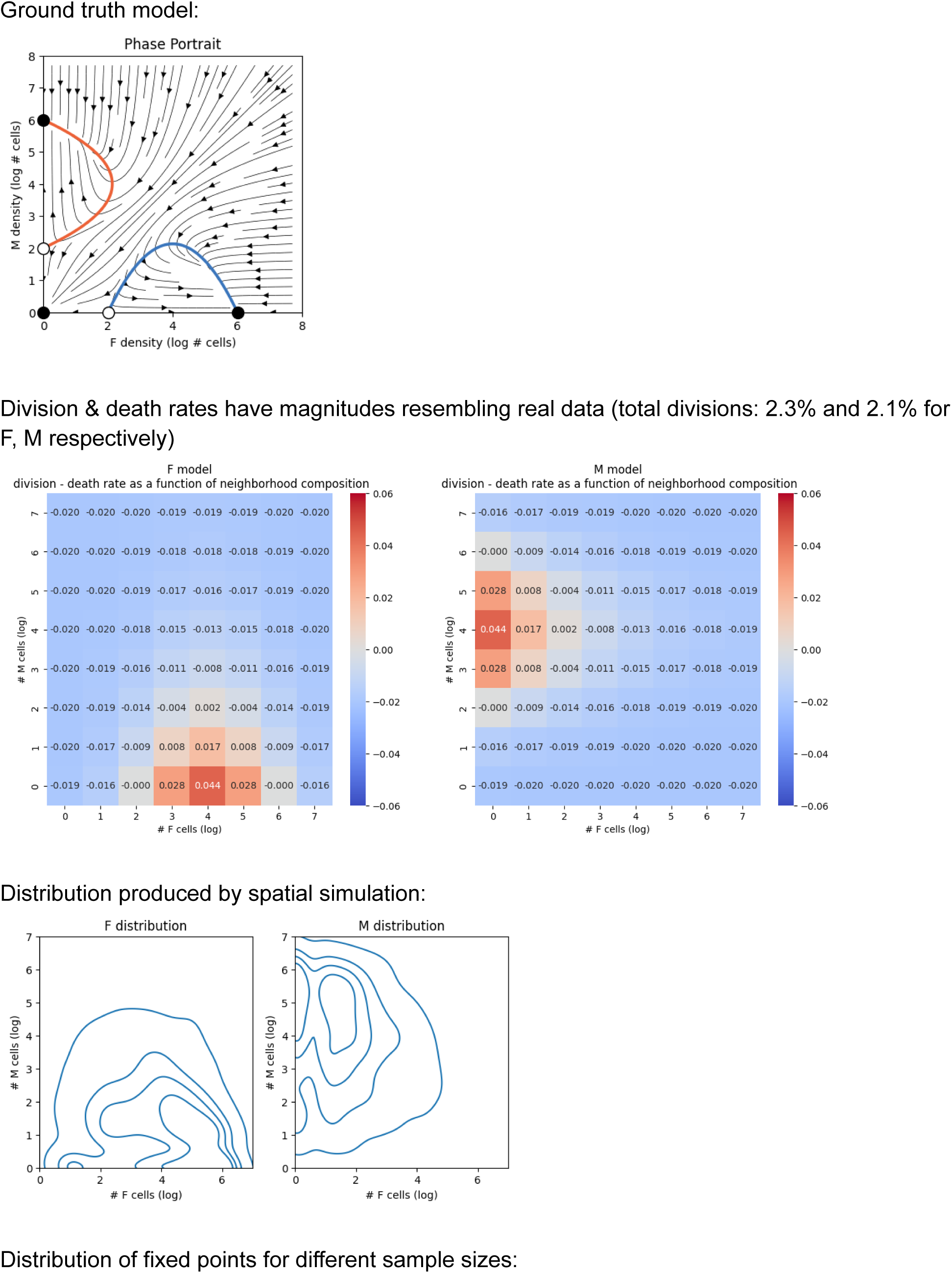
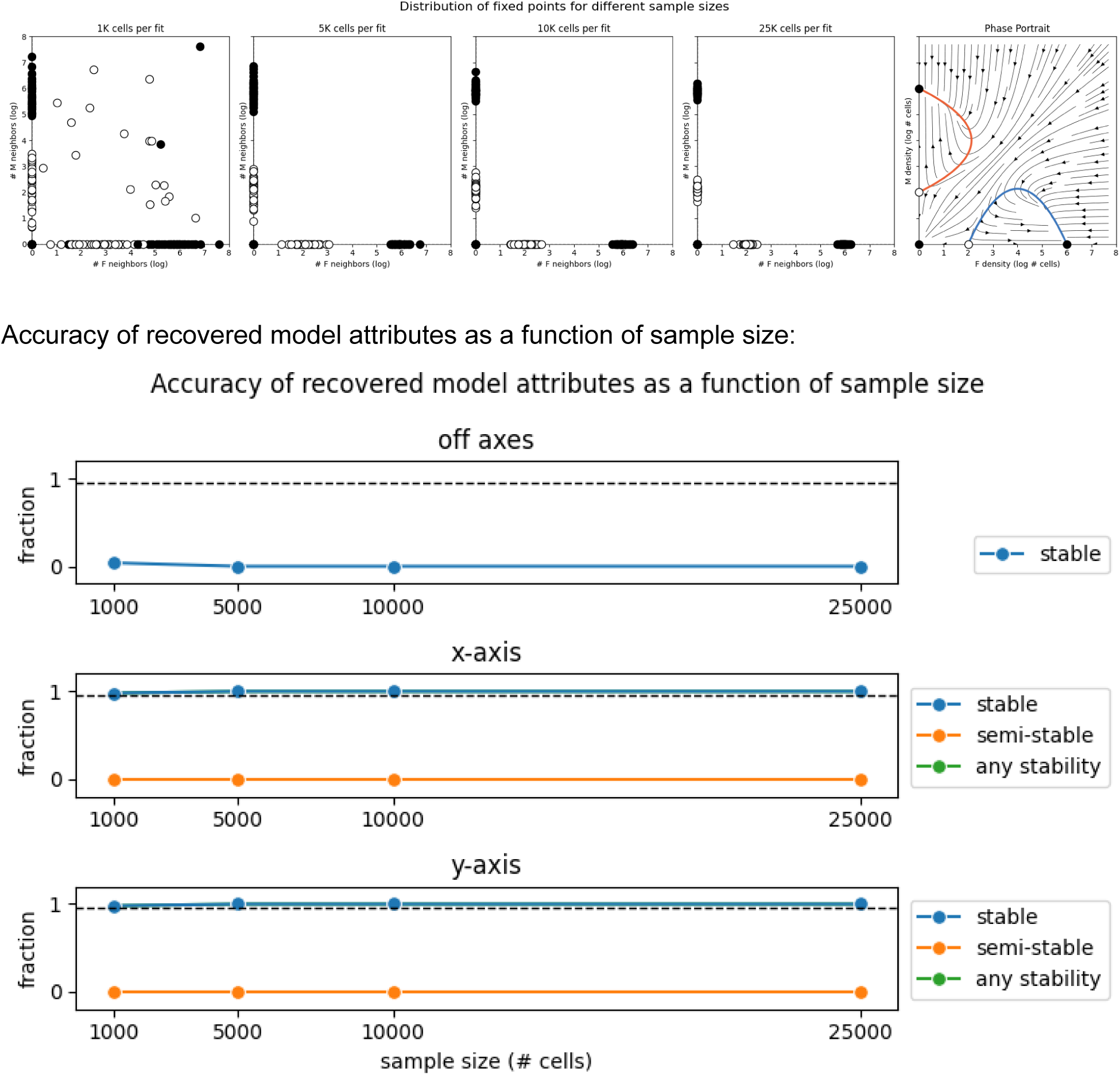

### S2K - Simulation 4 - Four fixed points

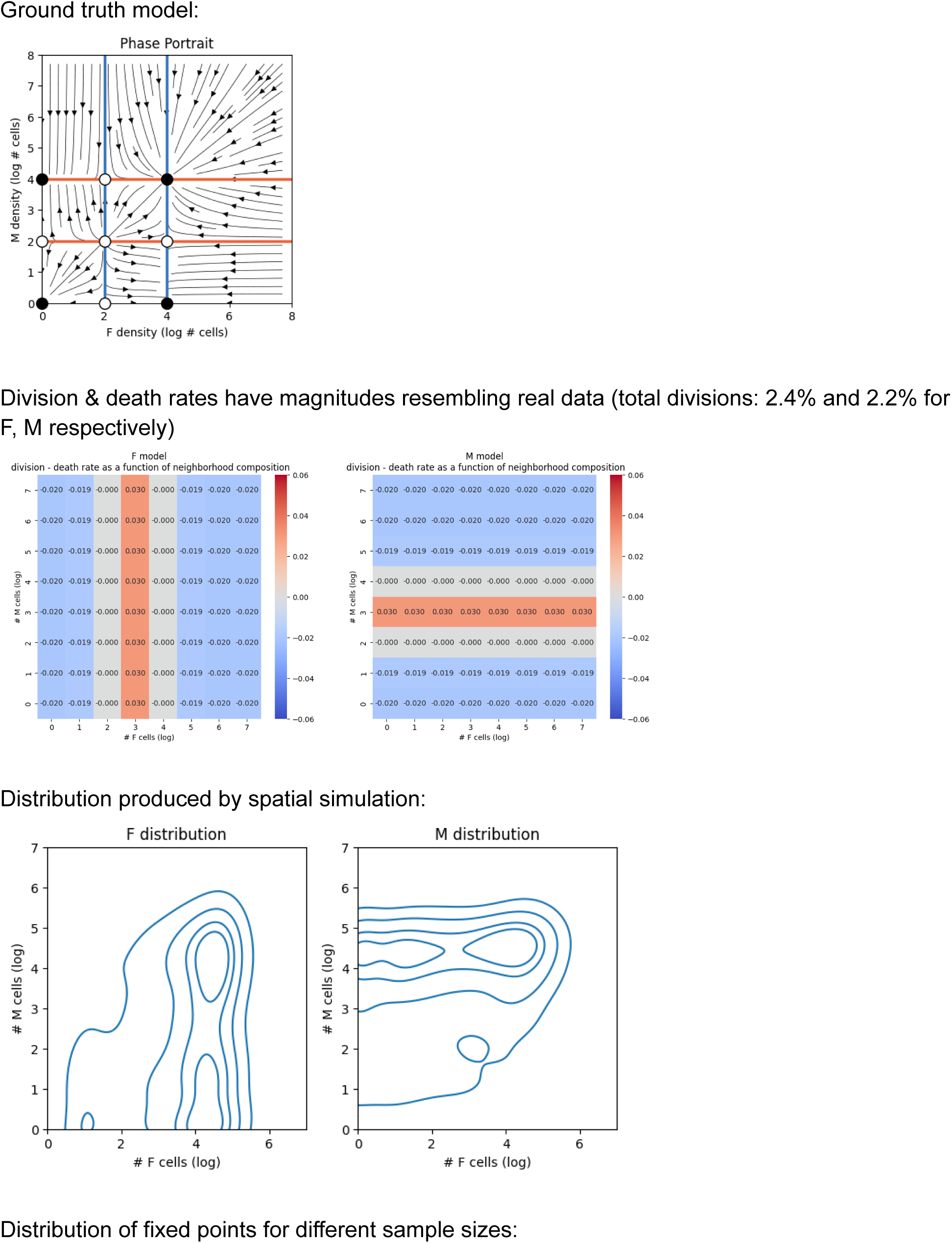
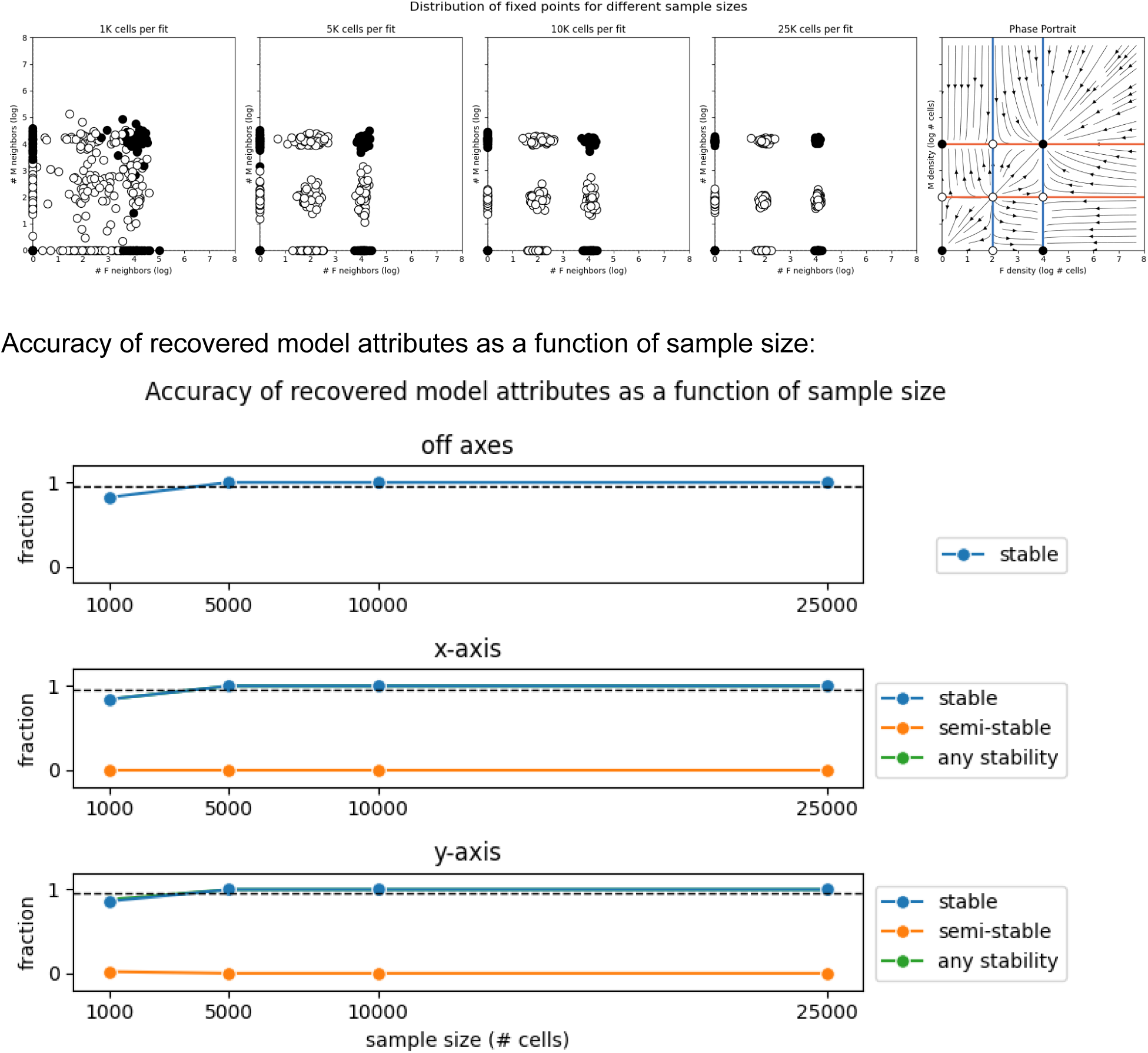

## Supplementary 3 - Fibroblast-macrophage dynamics

### S3A - Model selection using cross-validation suggests a 2nd order polynomial

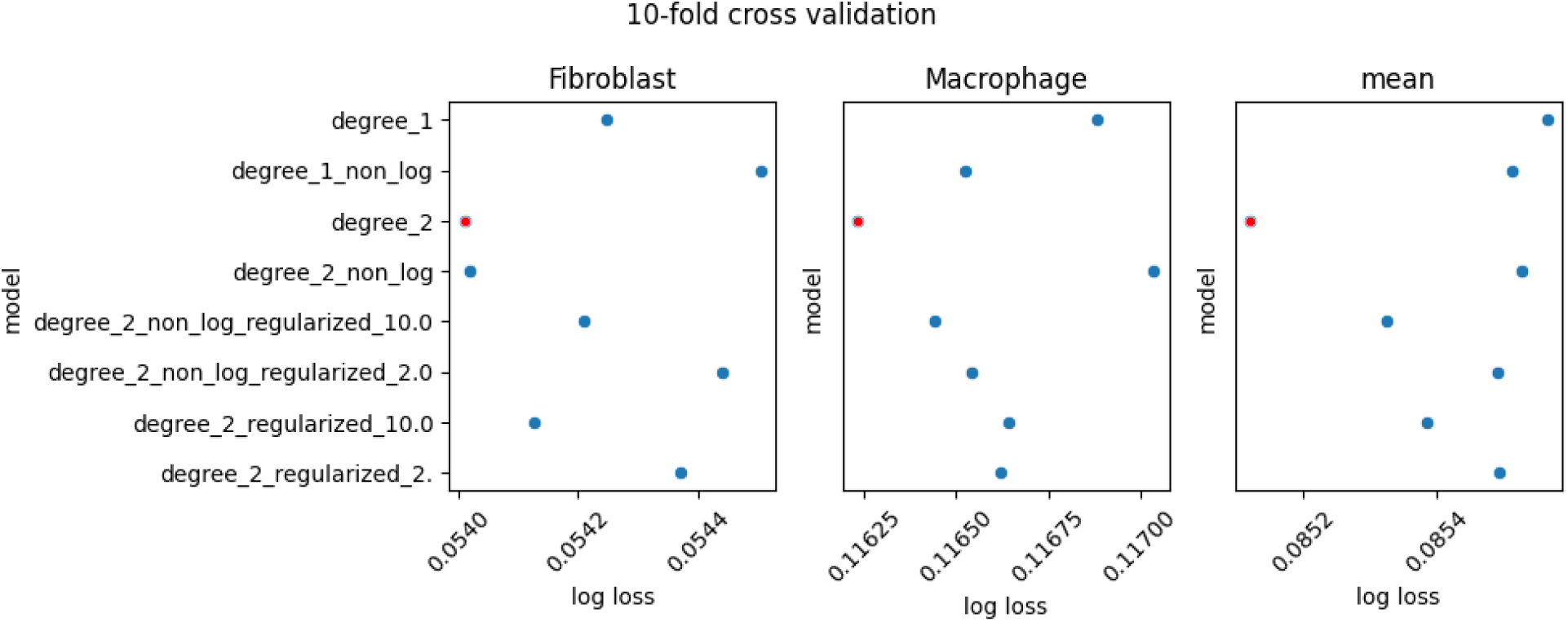

### S3B - The Fibroblast-Macrophage phase-portrait is robust to cell-level bootstrap resampling

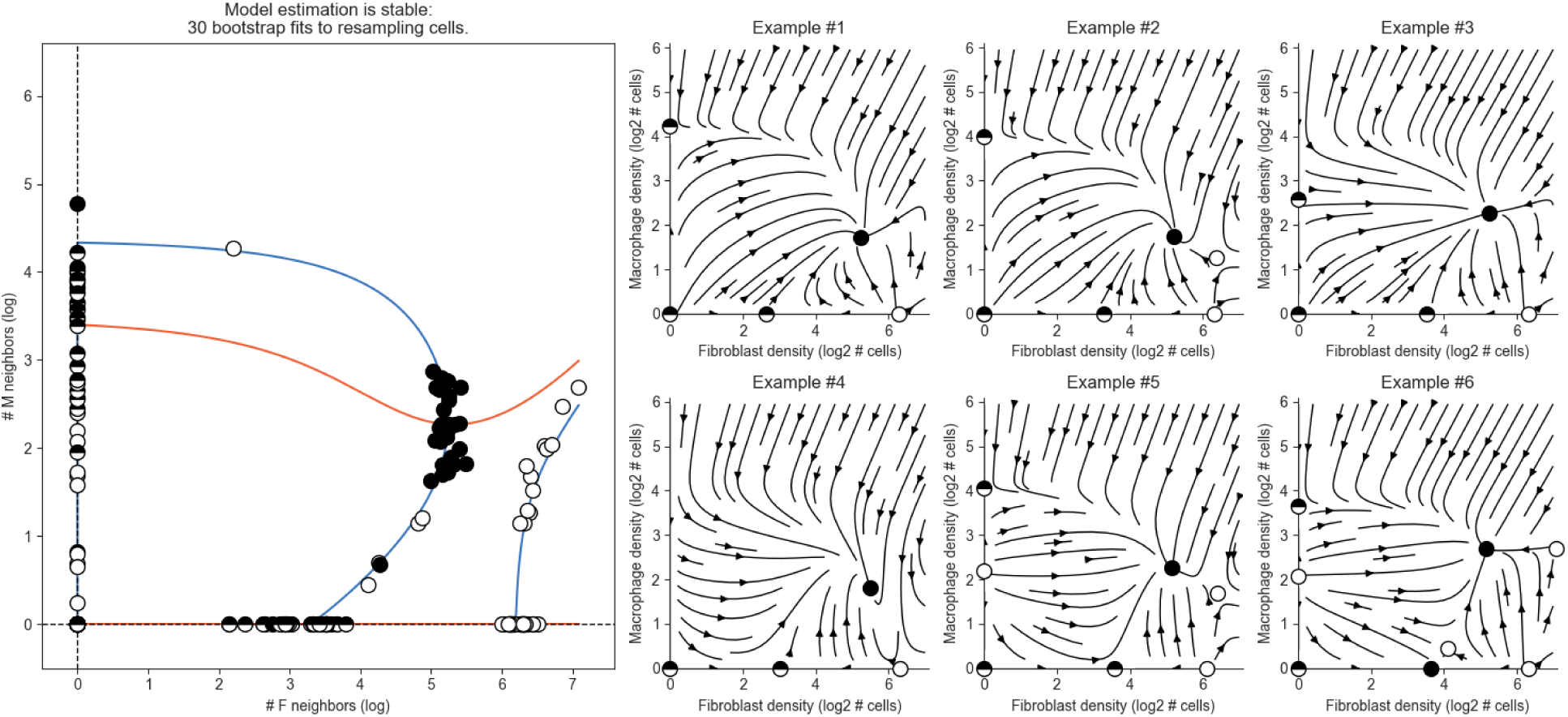

### S3C - The Fibroblast-Macrophage phase-portrait is robust to patient-level bootstrap resampling

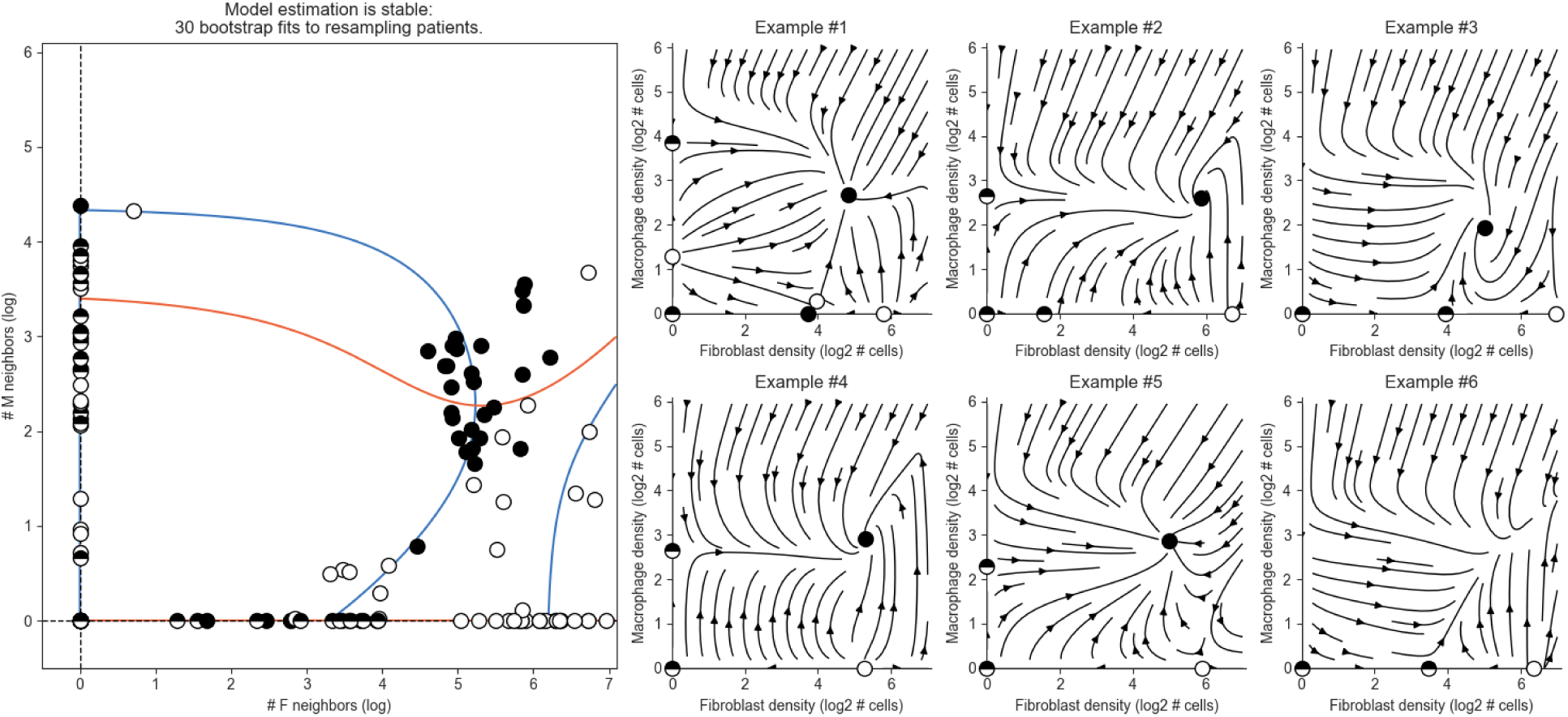

### S3D - Fibroblast-Macrophage vector field without smoothing

We plot the fibroblast-macrophage vector field over a finite grid. Arrow sizes correspond with magnitude of change in <u>absolute</u> numbers of cells. Arrows in gray are velocities from 10 bootstrap samples. See table XXX for velocities at all neighborhood compositions.

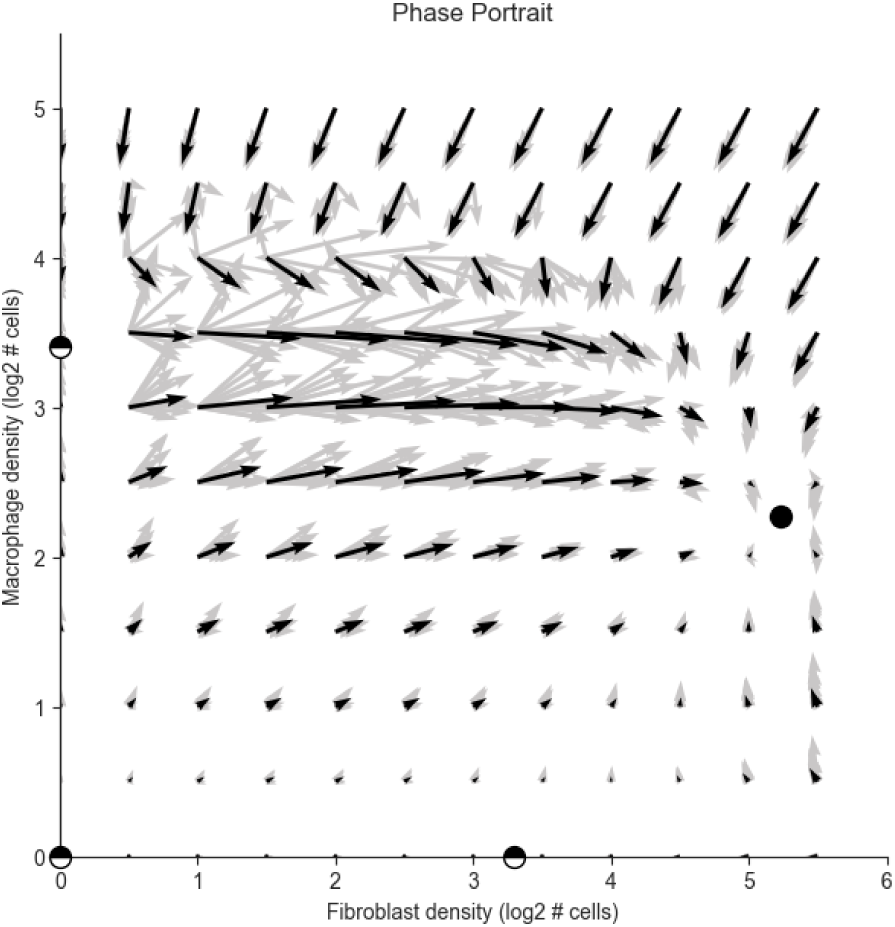

### S3E - Number of cells and divisions as a function of neighborhood size and Ki67

Because we exclude fibroblasts or macrophages with neighboring adaptive immune cells, the number of cells included in the analysis depends on the neighborhood size. We plot the number of cells and divisions for each combination. In S3F we show that analysis is robust to this choice. Cell numbers drop with neighborhood size because larger size increases probability of T and B cells, and thus exclusion of that neighborhood.

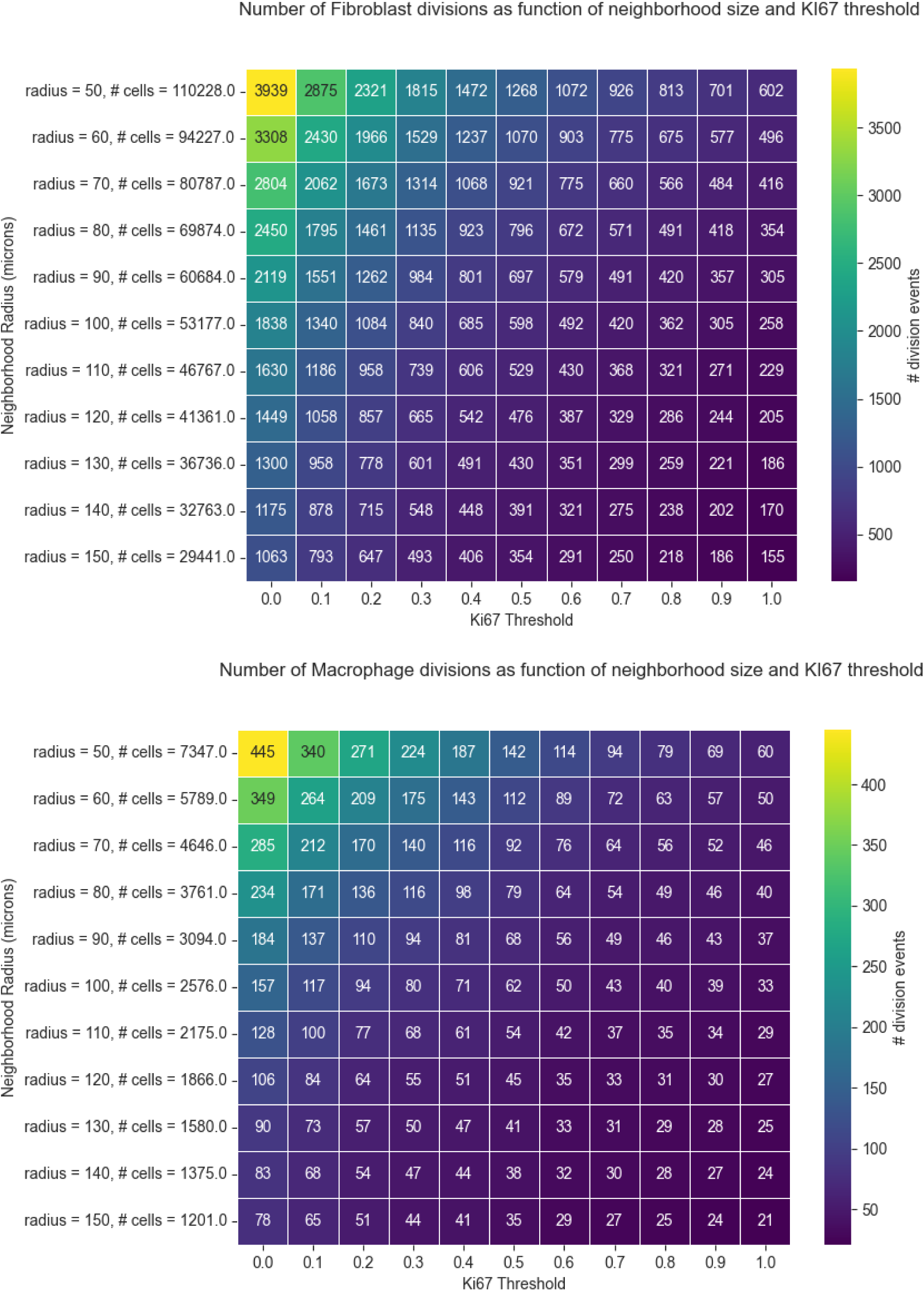

### S3F - The Fibroblast-Macrophage phase-portrait is robust to choice of Ki67 threshold and neighborhood size

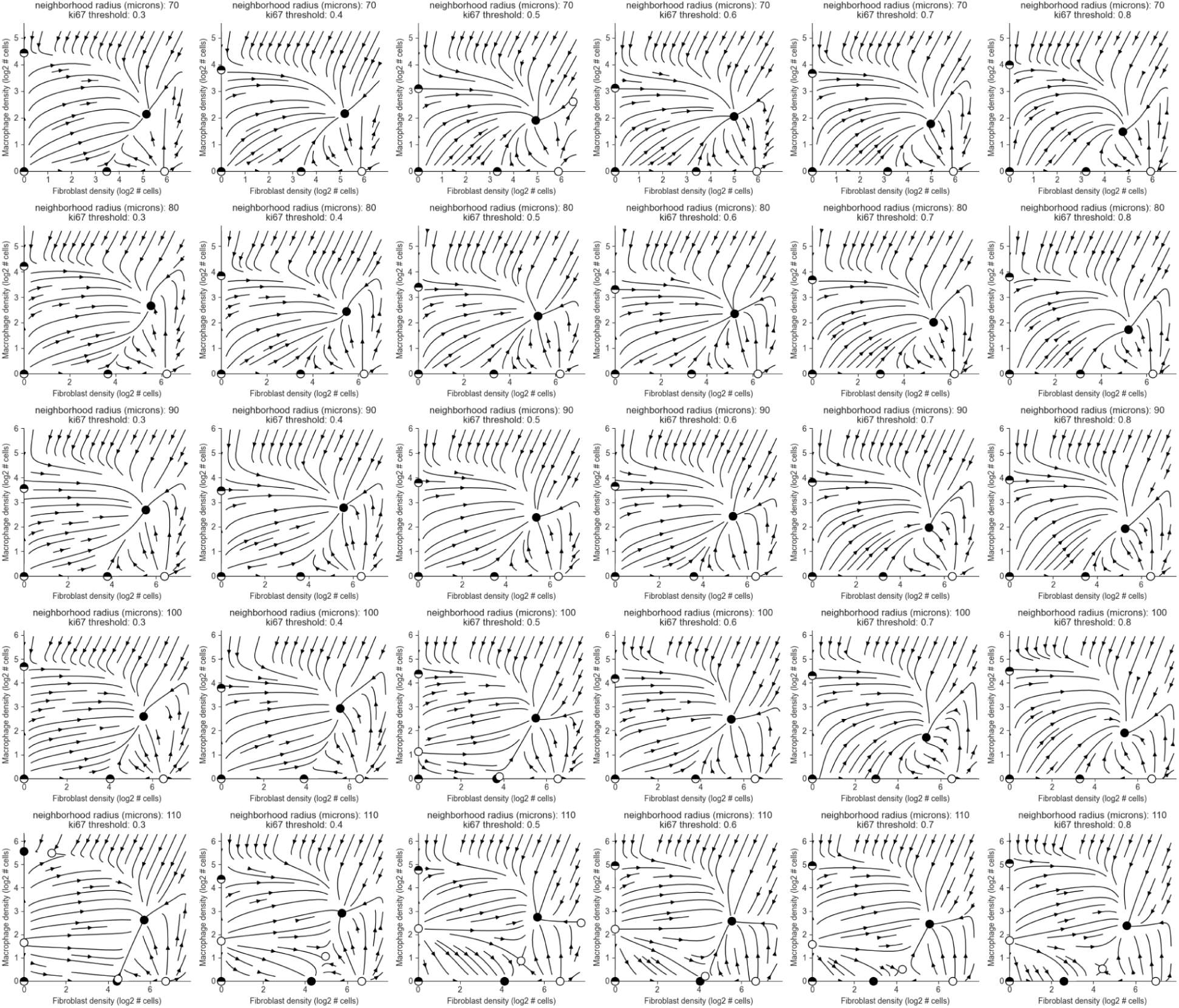

### S3G - The Fibroblast-Macrophage phase-portrait is robust to perturbation of the death estimate

Here we plot the phase-portrait against different constant death rates. The constant used throughout this study is equal to the mean division rate. This approximation is exact at steady state. We show that dynamics are largely preserved even for constants that produce a doubling or halving time of 2 weeks for a cell population.

The computation of the doubling time is as follows: say a cell has a mean division rate of 0.02 per time step. For a death rate that is 70% of the division rate, at each time step we gain (0.02 - 0.02*0.7) x the number of cells. Note this is an exponential growth.

We get a doubling time of:

T = log(2) / log(1 + 0.02 - 0.02*0.7) = 115 steps

This is ∼2 weeks for a timestep of 3 hours.

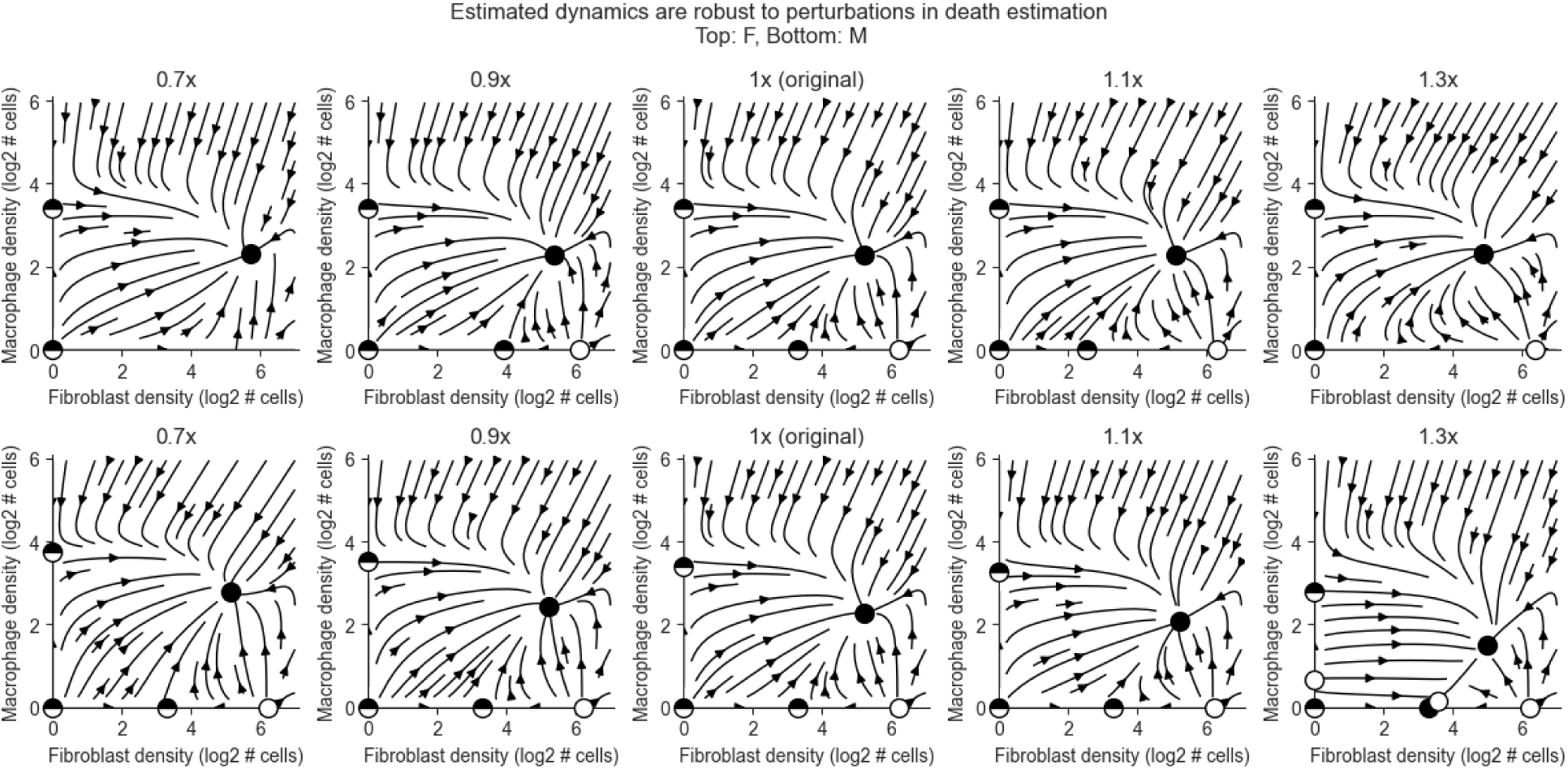

### S3H - Fibroblast-Macrophage dynamics are consistent when modeled together with a third cell types

Tumor and Endothelial cells were present in the neighborhoods in the main analysis but weren’t included in the model (recall, neighborhoods with adaptive immune cells were excluded). Here, we fit a model including 3 cell types each time, then dynamics are plotted for values ranging from the 20th to the 80th quantile density of the 3rd cell type.

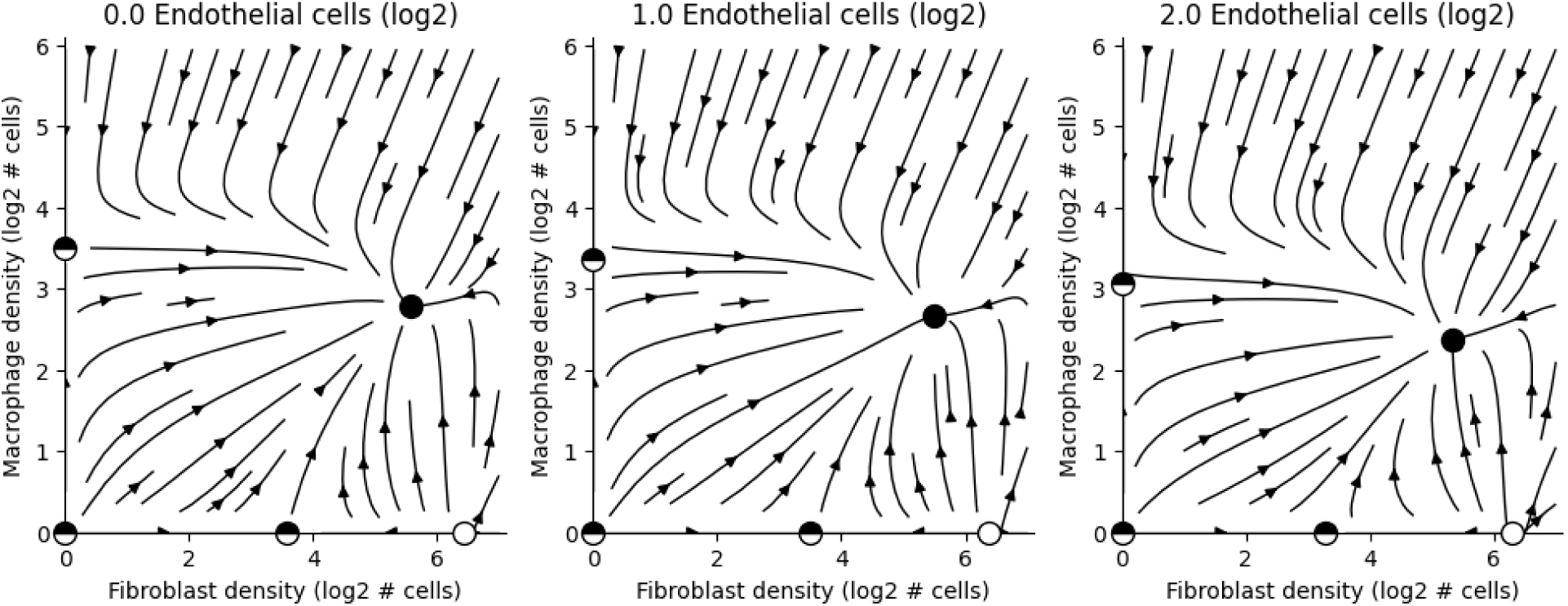
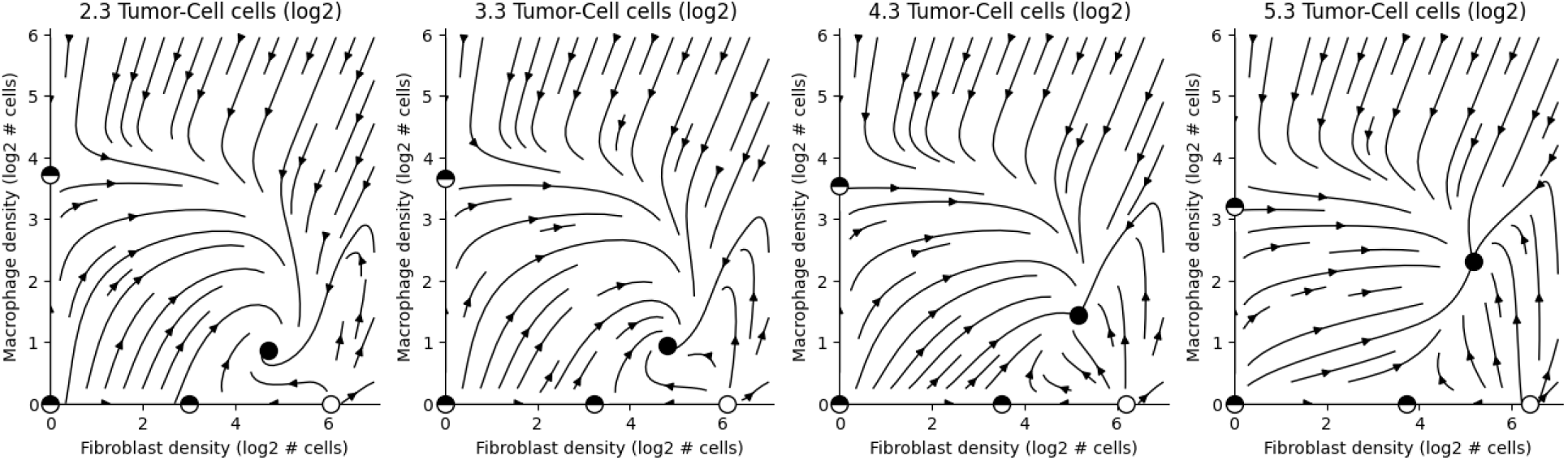

### S3I - Fibroblast Macrophage Dynamics are conserved across various patient subgroups

We fit a model independently to subgroups of patients. The complete cohort had only 4K macrophages so partitioning the cohort leaves just a few thousand per subgroup. As such we partitioned into two groups in all cases, even if more subgroups could be warranted. We note the number of samples in parentheses in order to make more explicit when the quality of a phase-portrait likely results from a small sample size. Despite partitioning this initially low sample size in half or less, many elements of the original phase portrait are preserved.

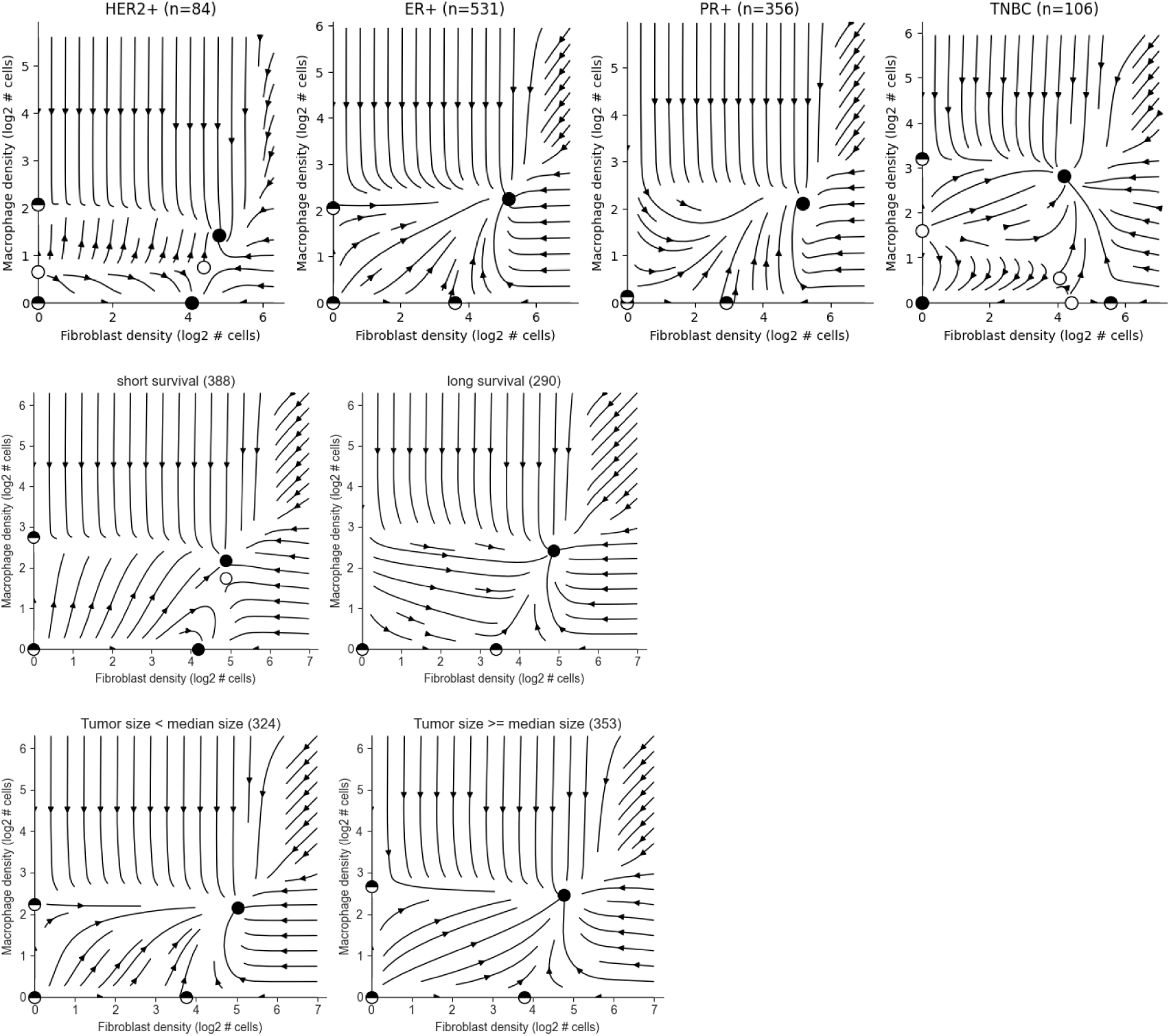
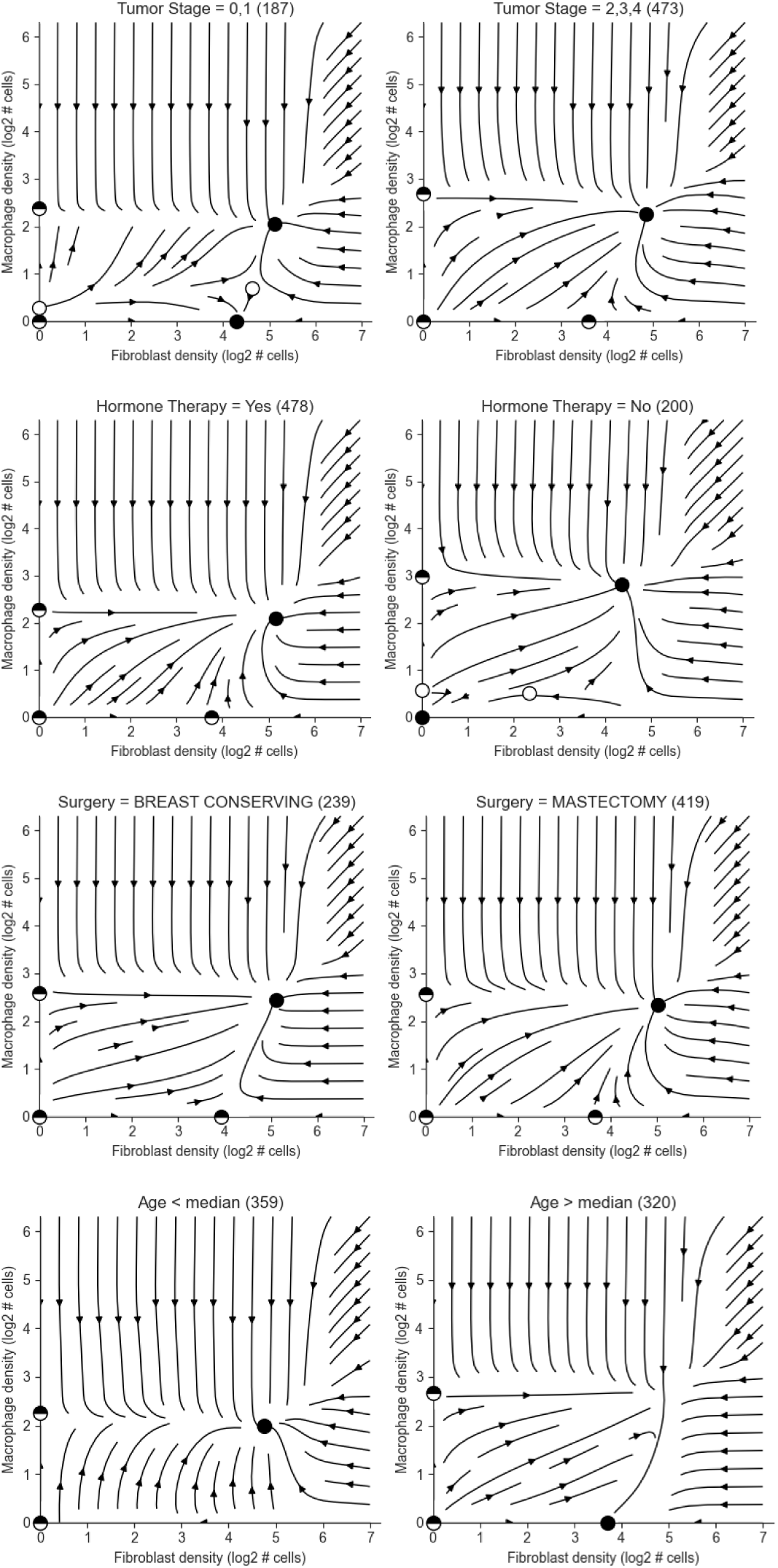

### S3J - Cox proportional hazards model for survival as a function of tumor and macrophage density shows a significant correlation between macrophage density and poor survival

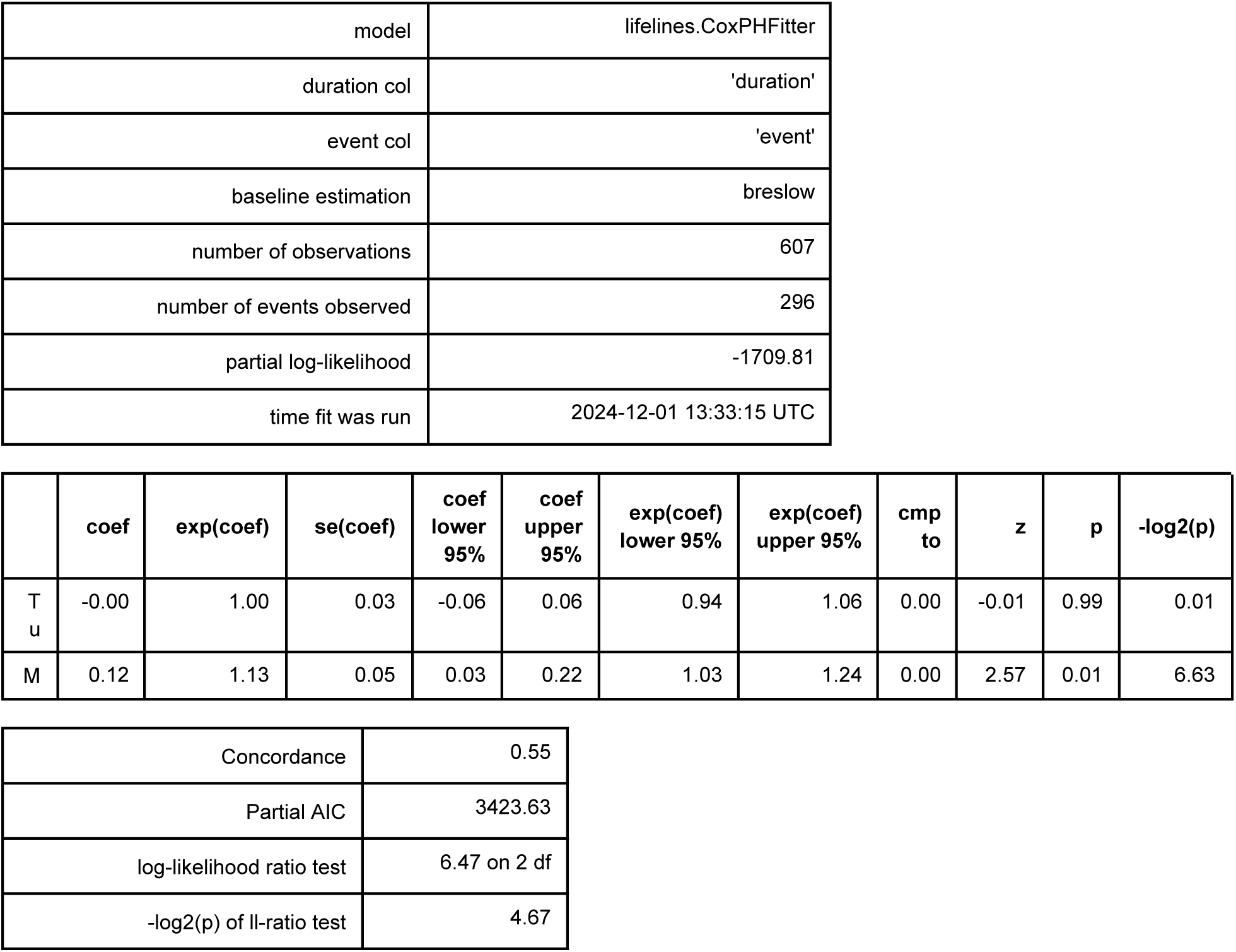

### S3K - The FM phase-portrait exhibits cold and hot fibrosis across 3 independent datasets

Note that without controlling for the level of tumor cells, the Fischer dataset displays only a cold-fibrosis stable point. When modeling tumor cells and fixing their value we see both cold and hot fixed points (S3M). As the tumor controls the type of fixed point, a different fraction of healthy sections across the datasets could produce such differences.

**The FM phase-portrait in 3 independent datasets:**

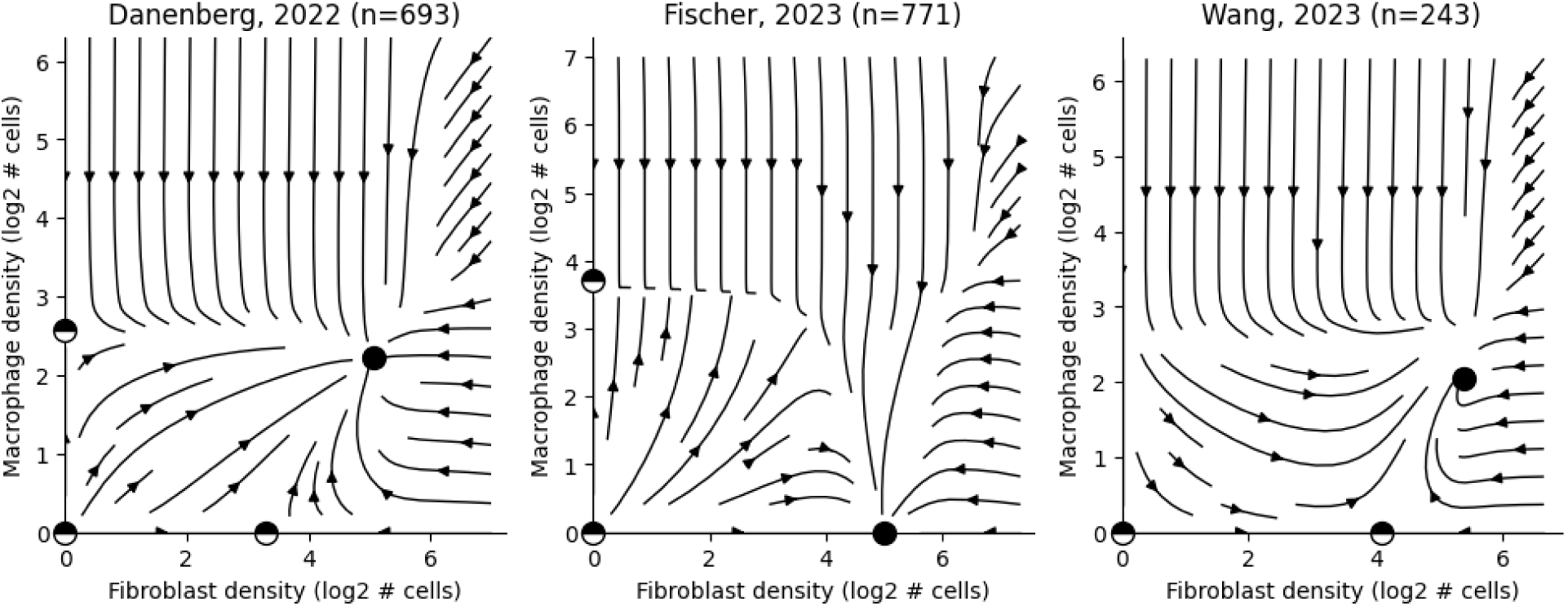

The FM phase-portrait in 3 independent datasets, fixing tumor density to 80 cells (this is approximately the point from which the Fischer dataset displays both fixed points, see the figure below for a range of values of Tumor density in all 3 datasets):

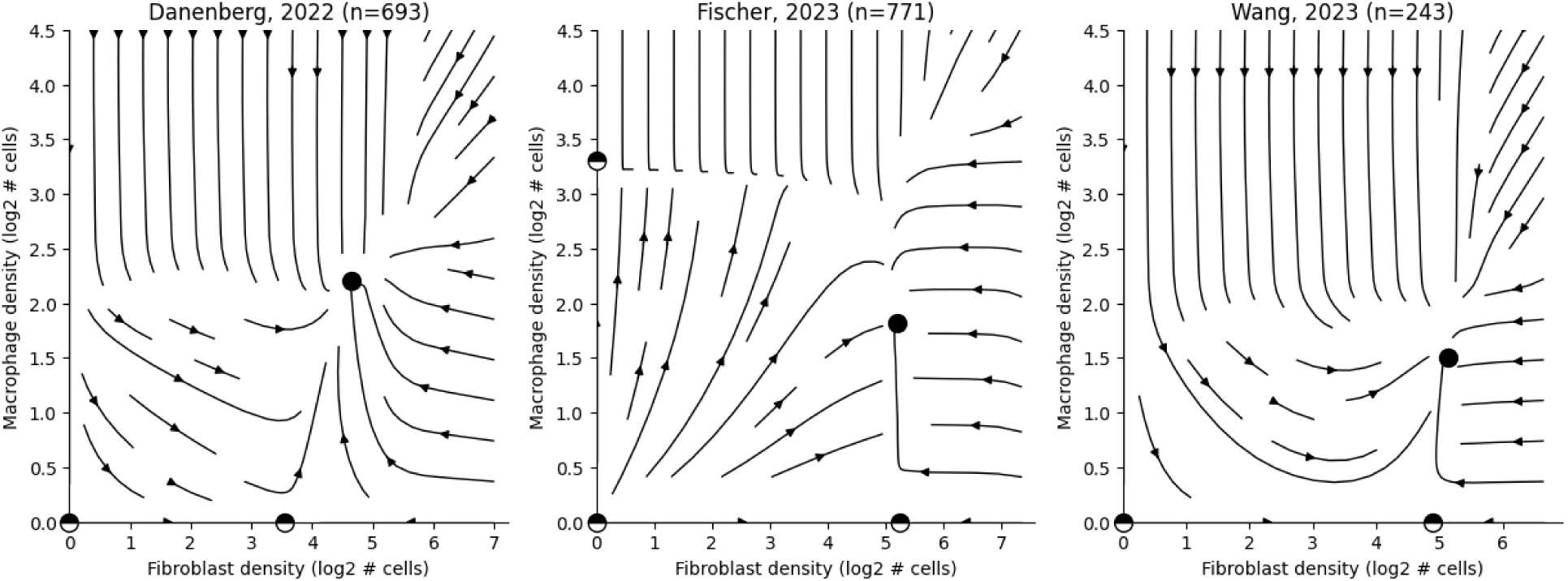

### S3L - Phase-portraits under increasing densities of tumor cells in 3 datasets

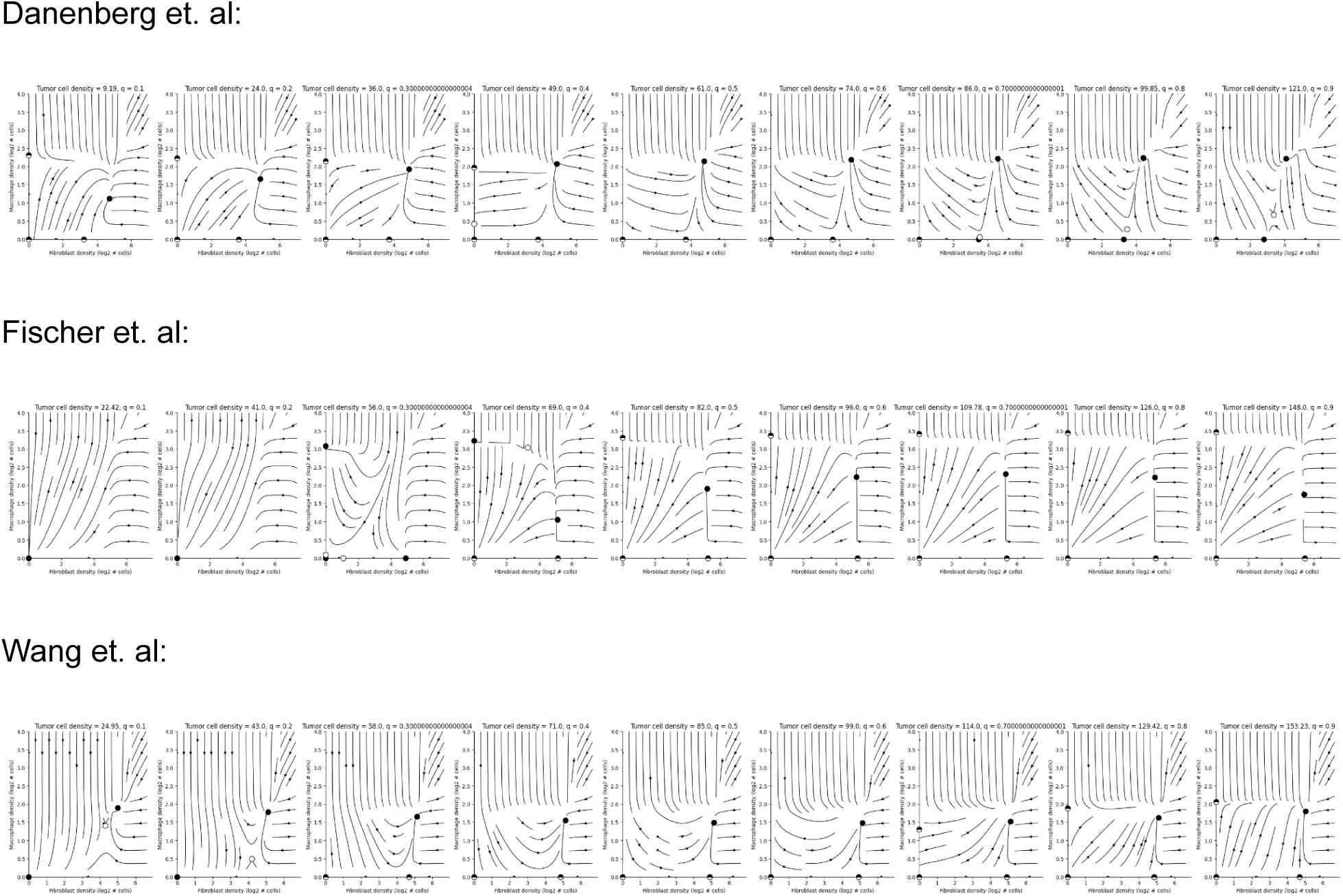

### S3M - Estimating magnitude of error due to unmodeled terms: Empirical position of tissues in state-space

Estimated dynamics drive tissues towards a region with a high density of tissues in state-space. If dynamics were completely deterministic all tissues would converge to the fixed points. We expect the error to have dispersing effect.

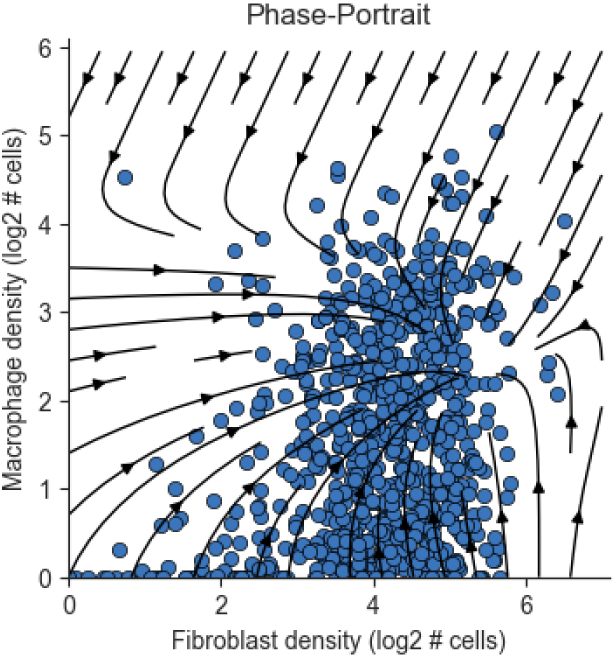

### S3N - Estimating magnitude of error due to unmodeled terms: Estimated density of tissues across the state-space

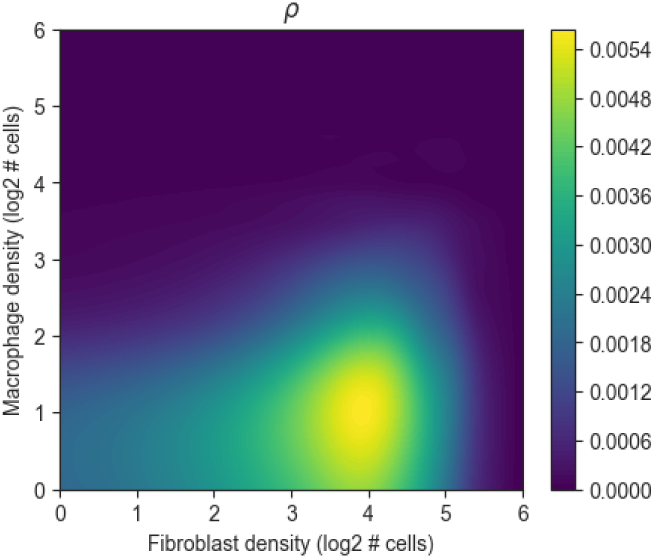

### S3O - Estimating magnitude of error due to unmodeled terms: Estimated divergence of the error field (without diffusion)

As expected, the error field produces a net effect that disperses cells from the peak density.

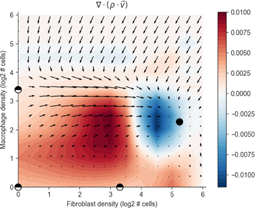

### S3P - Estimating magnitude of error due to unmodeled terms: Variance of error explained by diffusion

Accordingly, a simple (one parameter) diffusion explains 0.45 of the variance in the error and is statistically significant p=1e-23.

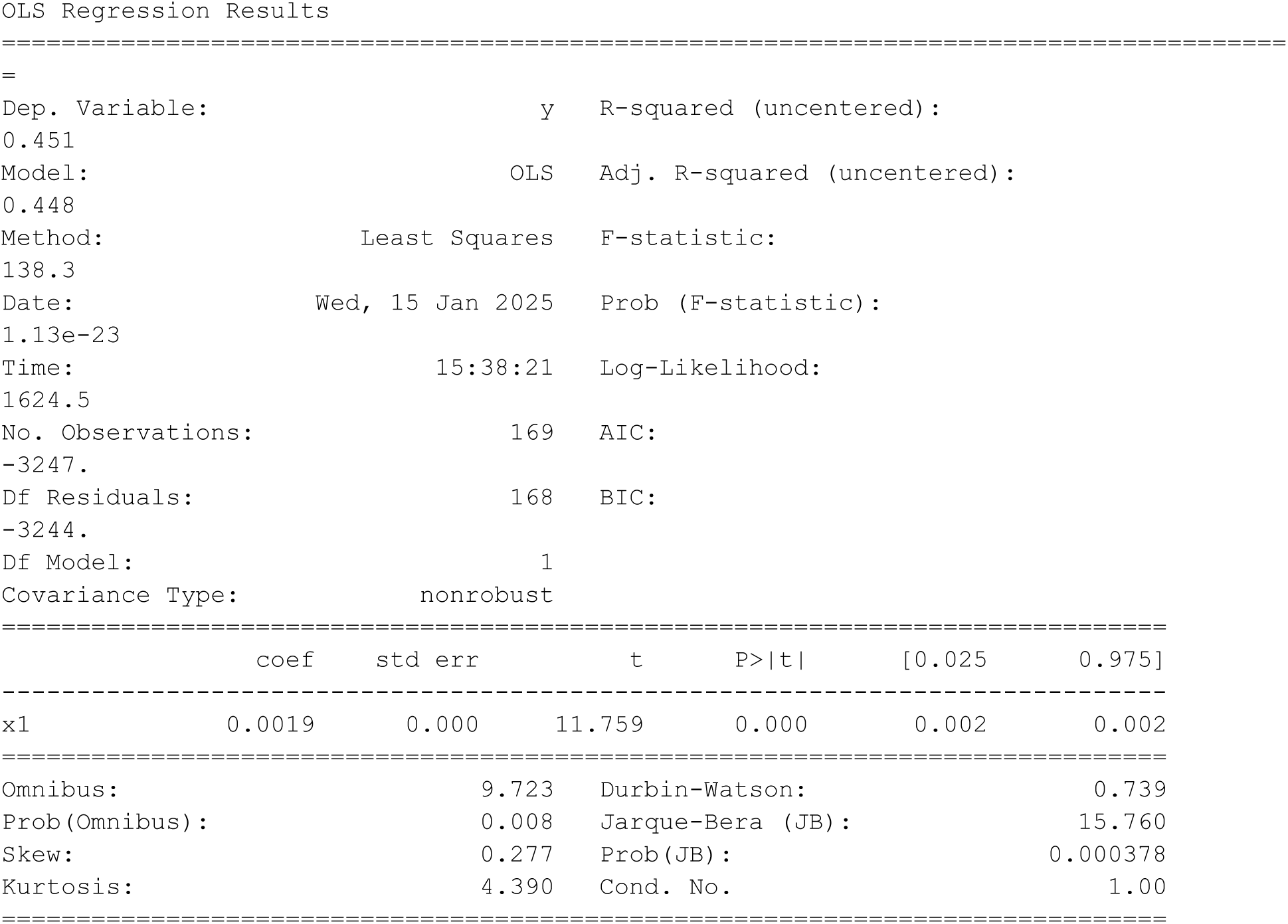

## Supplementary 4 - T-cell B-cell dynamics

### S4A - Model Selection

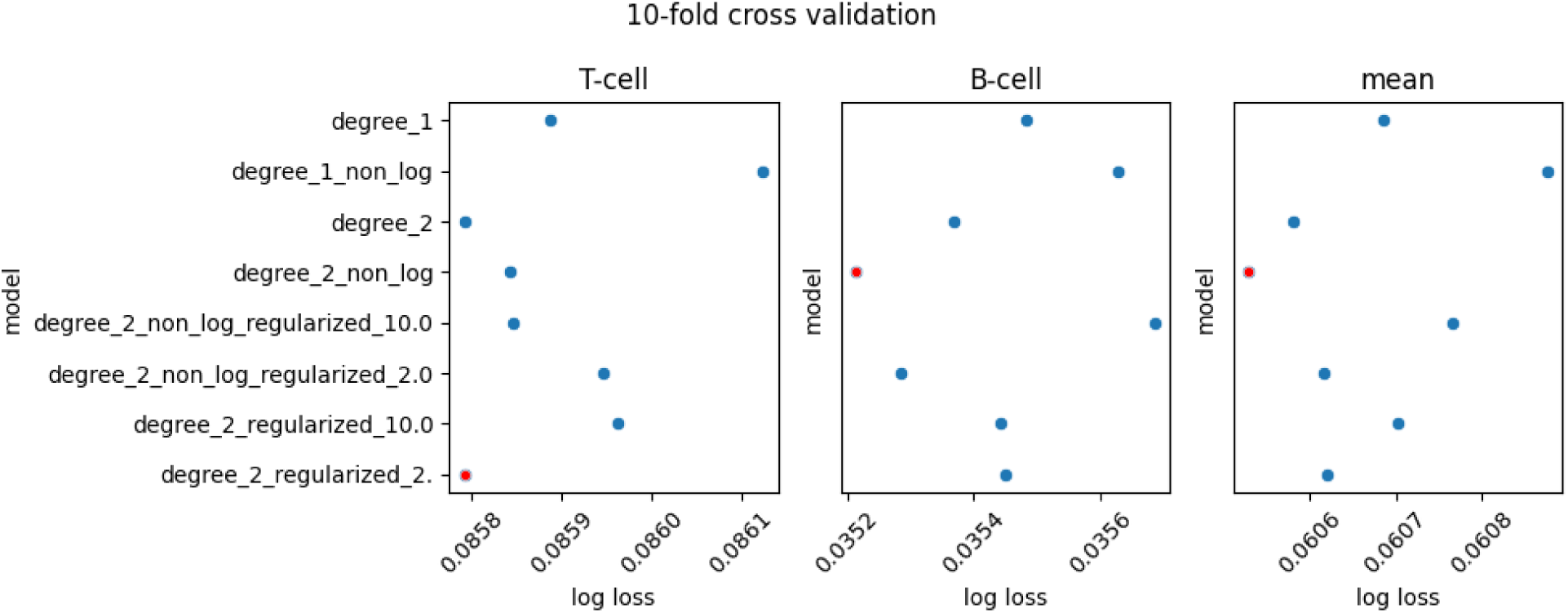

### S4B - The T-cell, B-cell phase-portrait is robust to cell-level bootstrap resampling

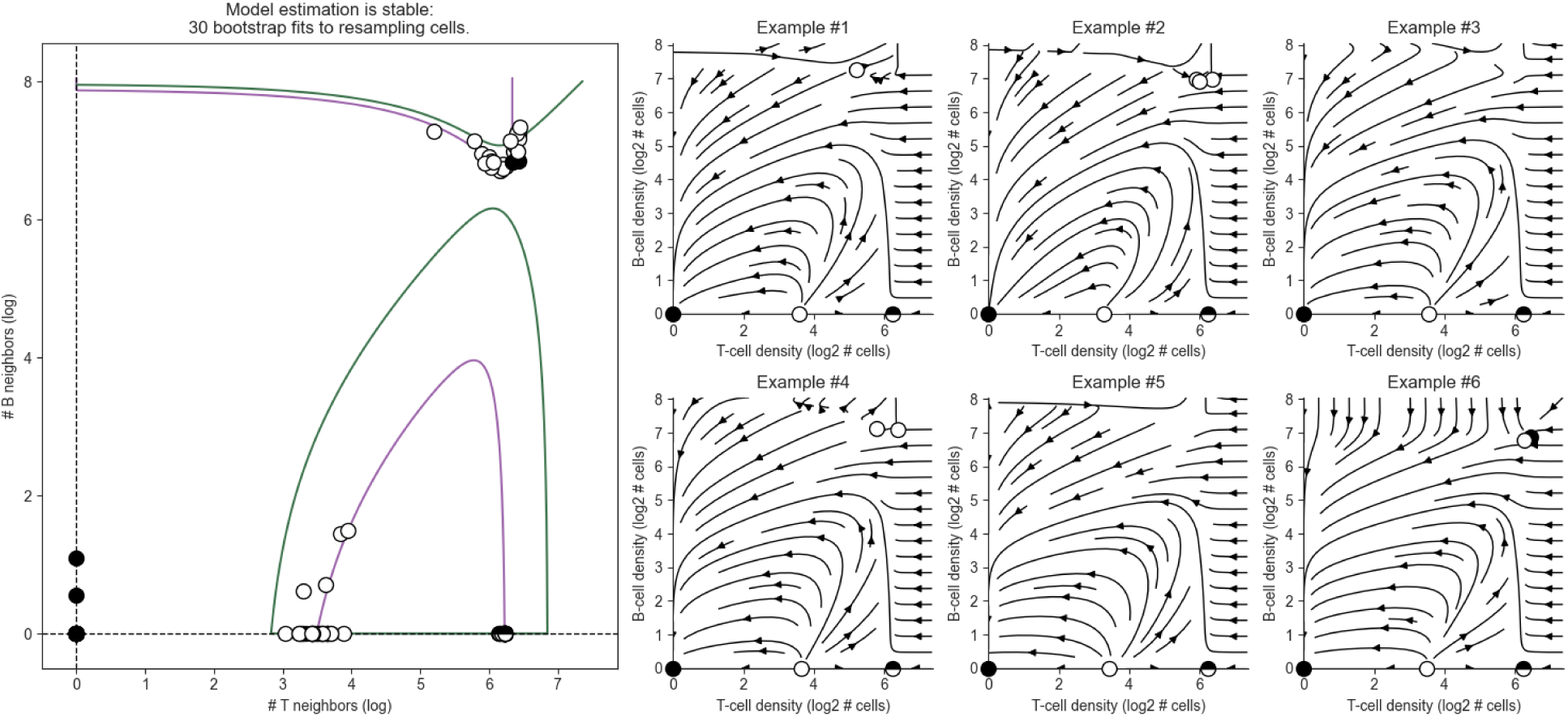

### S4C - The T-cell, B-cell phase-portrait is robust to patient-level bootstrap resampling

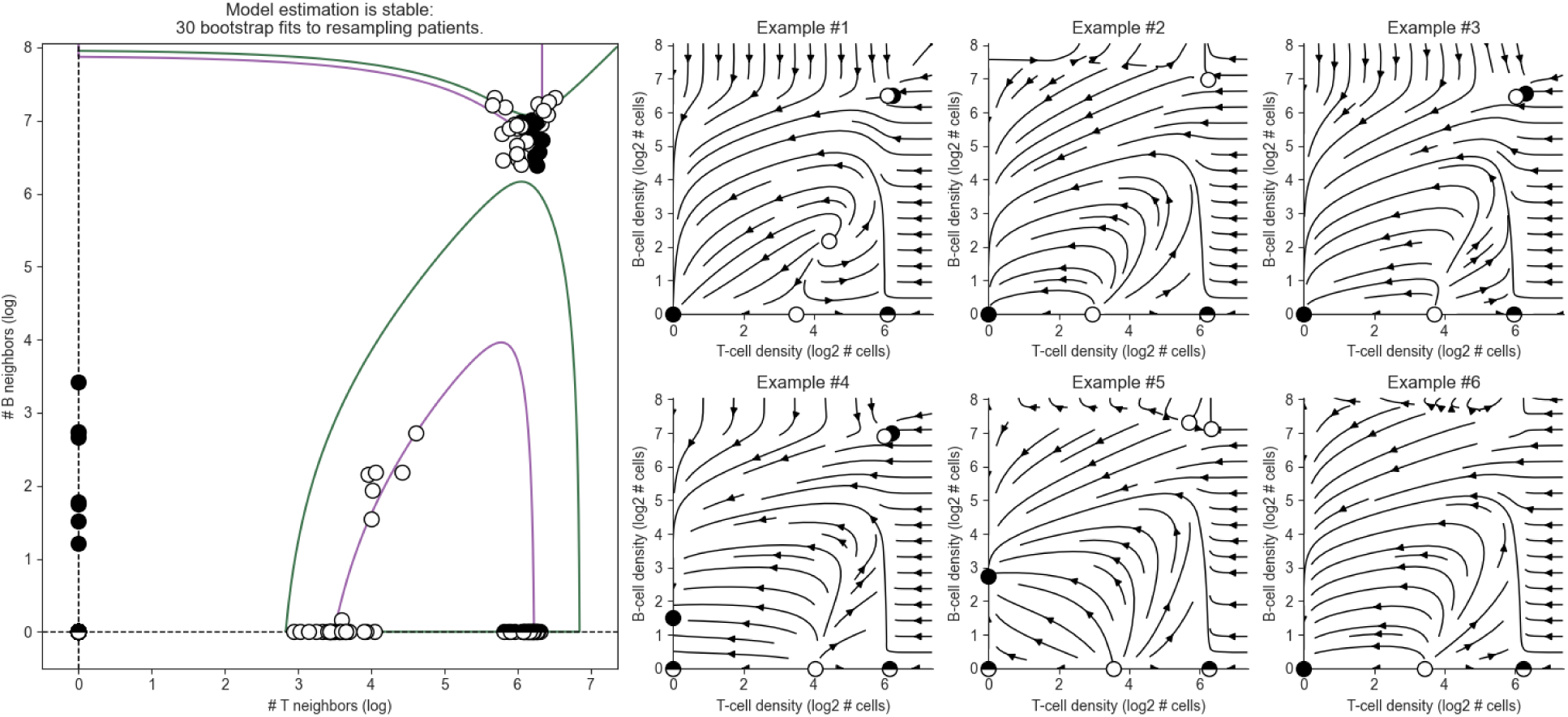

### S4D - T-cell B-cell vector field without smoothing

We plot the T-cell B-cell vector field over a finite grid. Arrow sizes correspond with magnitude of change in absolute numbers of cells. Arrows in gray are velocities from 10 bootstrap samples. See table XXX for velocities at all neighborhood compositions

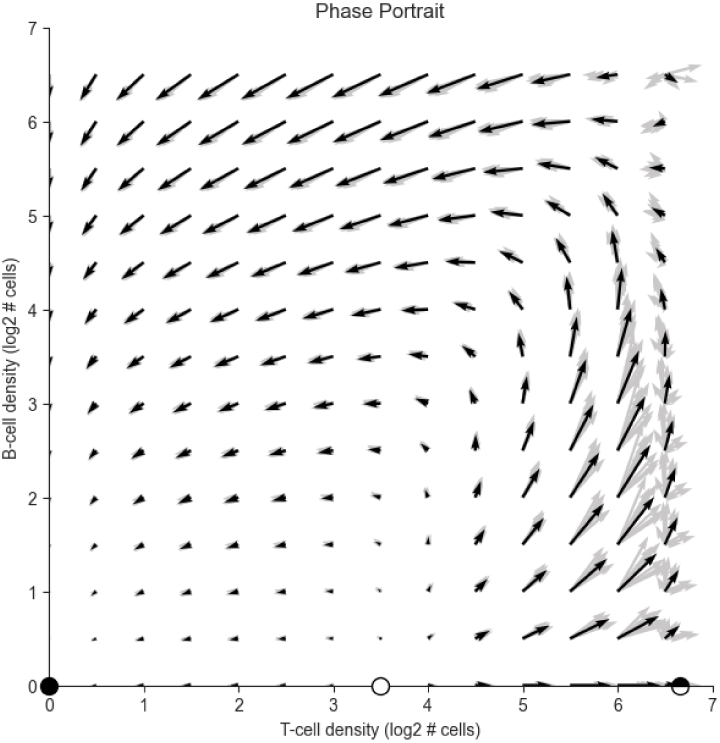

### S4E - The T-cell, B-cell phase-portrait is robust to perturbation of the death estimate

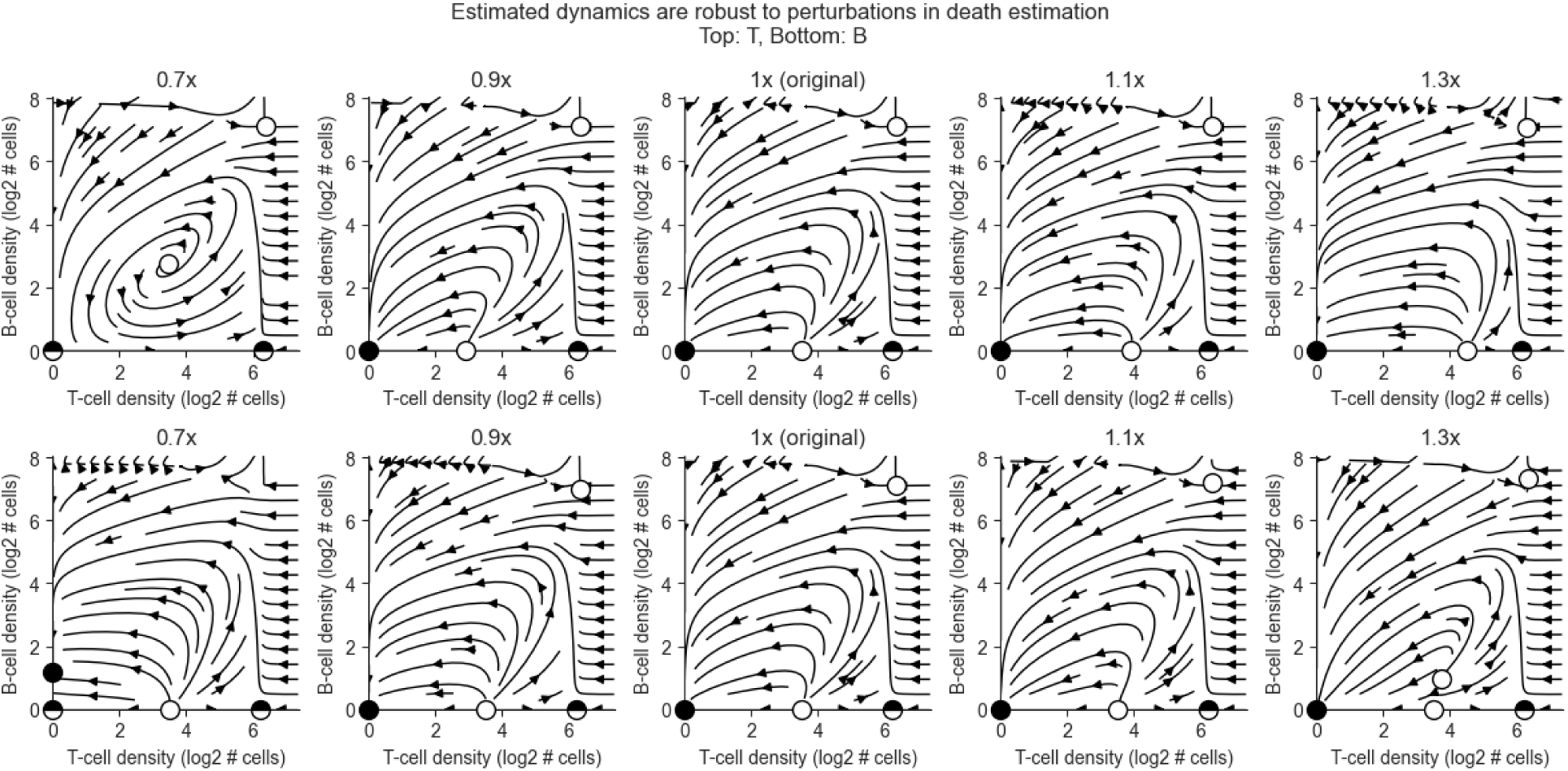

### S4F - Ki67 threshold determines fraction of dividing cells

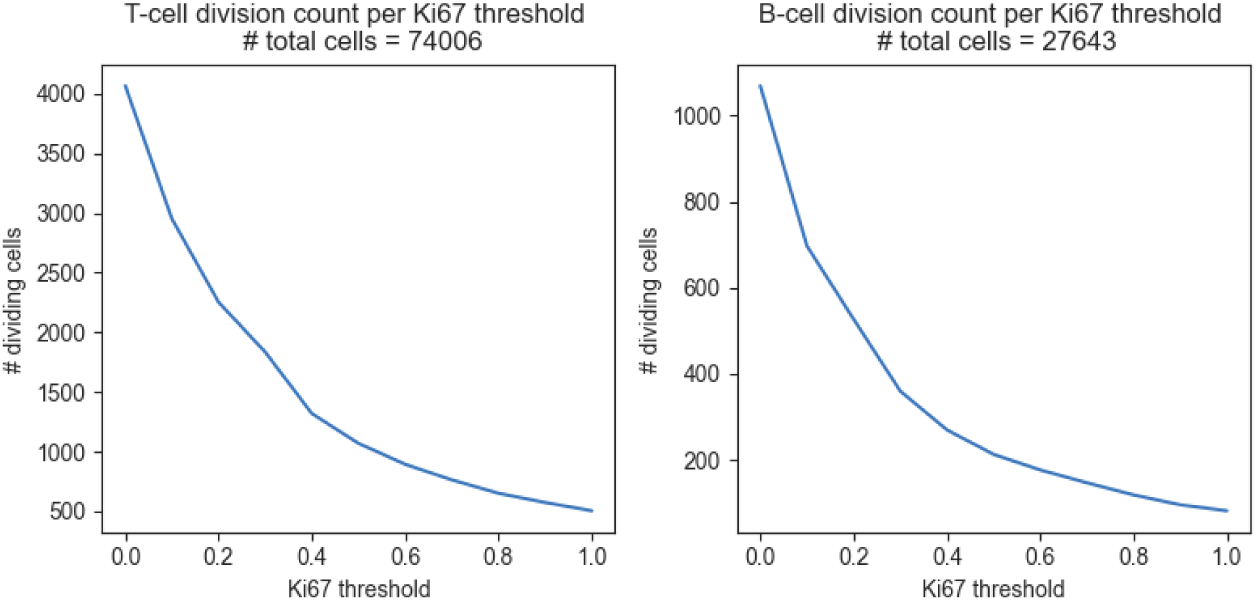

### S4G - The T-cell, B-cell phase-portrait estimate is robust to choice of Ki67 threshold and neighborhood size

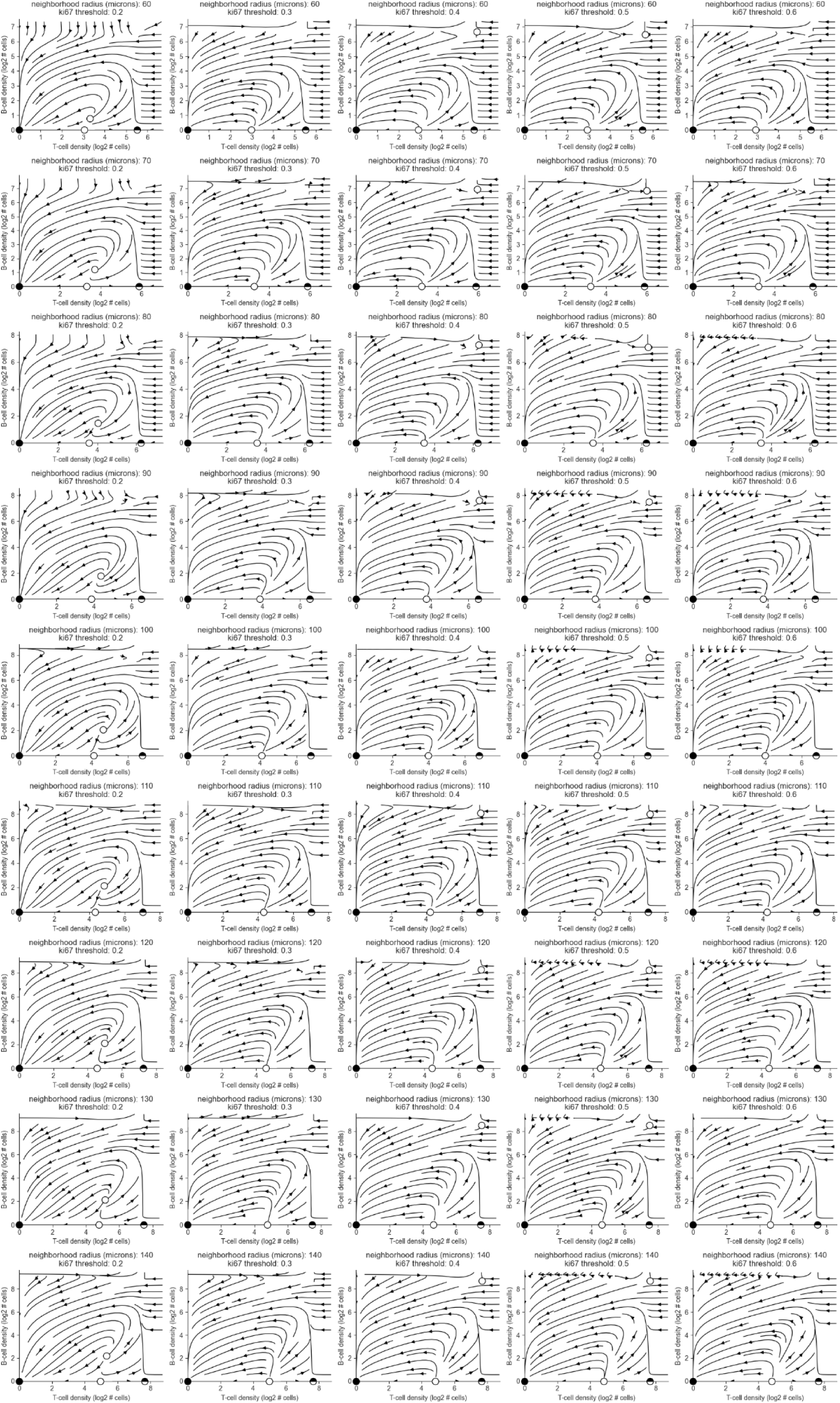

### S4H - T & B cell dynamics are consistent when modeled together with various cell types and under various levels of the 3rd cell

Fibroblast, Macrophage, Tumor & Endothelial cells were present in the neighborhoods in the main analysis but weren’t included in the model. Here, we fit a model including 3 cell types each time, then dynamics are plotted for values ranging from the 20th to the 80th quantile density of the 3rd cell type.

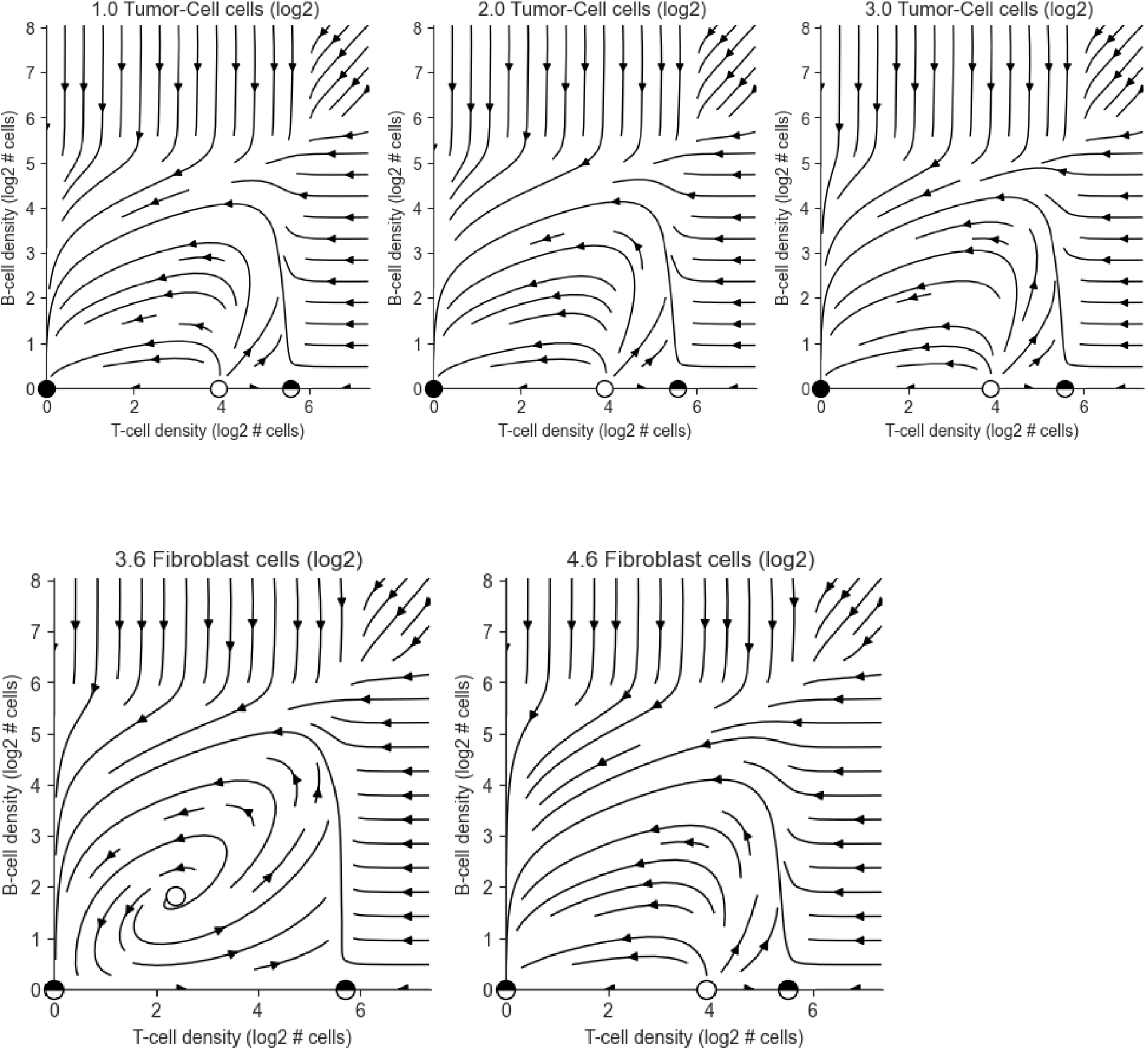
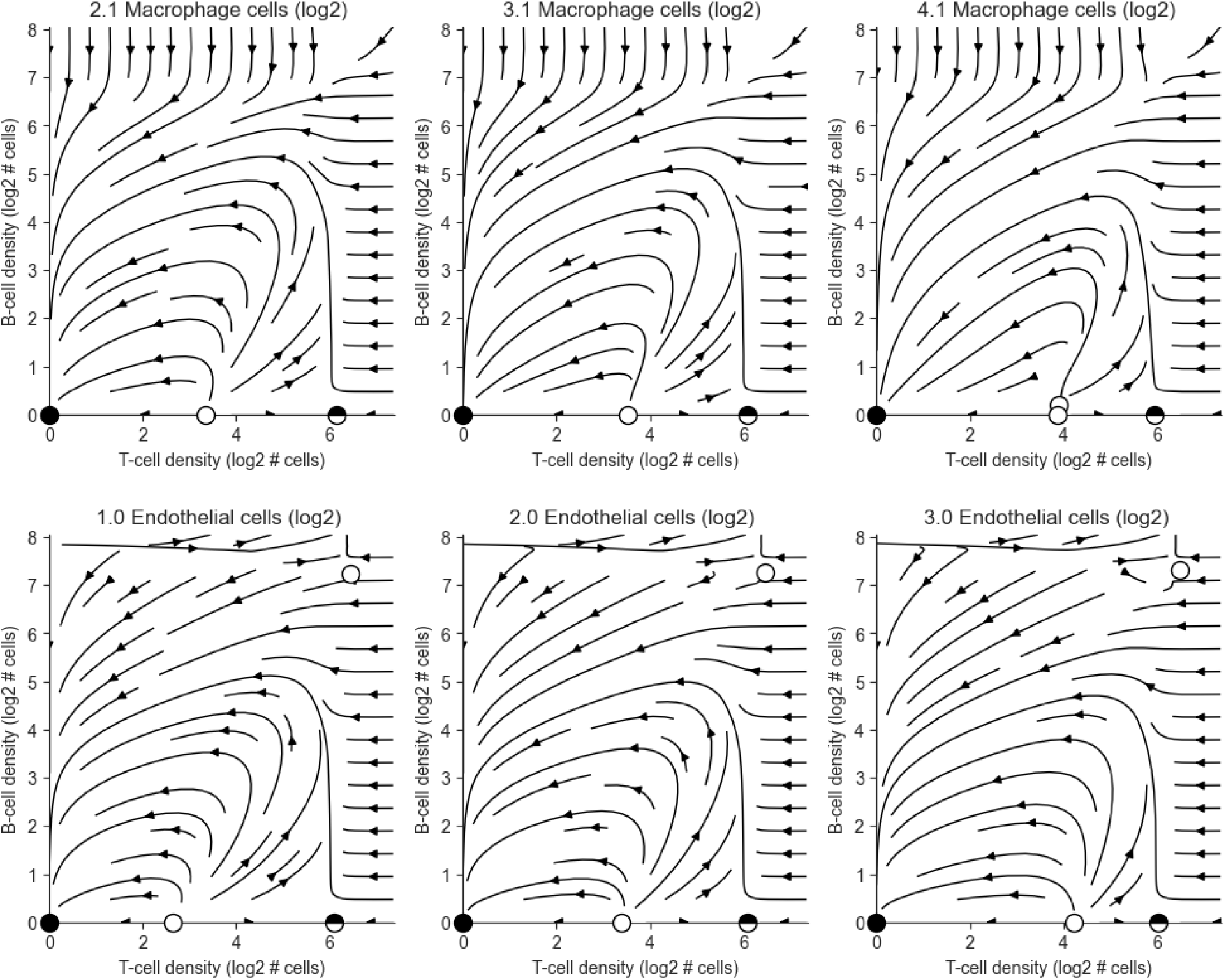

### S4I - T & B cell Dynamics are conserved across various patient subgroups

We test the model separately subgroups of triple-negative patients and receptor positive (HER2+, ER+, PR+) patients, as they display different dynamics. We note the number of patients in parenthesis to make explicit when the quality of a phase-portrait is likely due to low sample size.

Receptor positive patient phase-portraits:

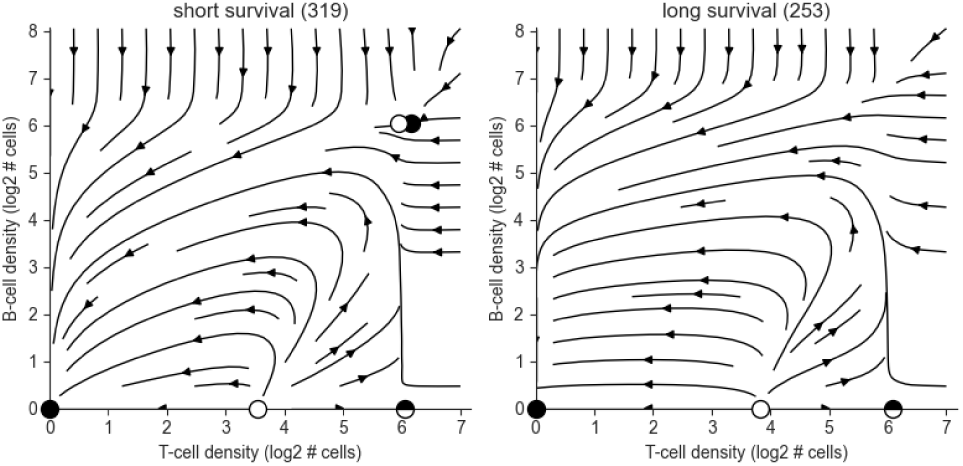
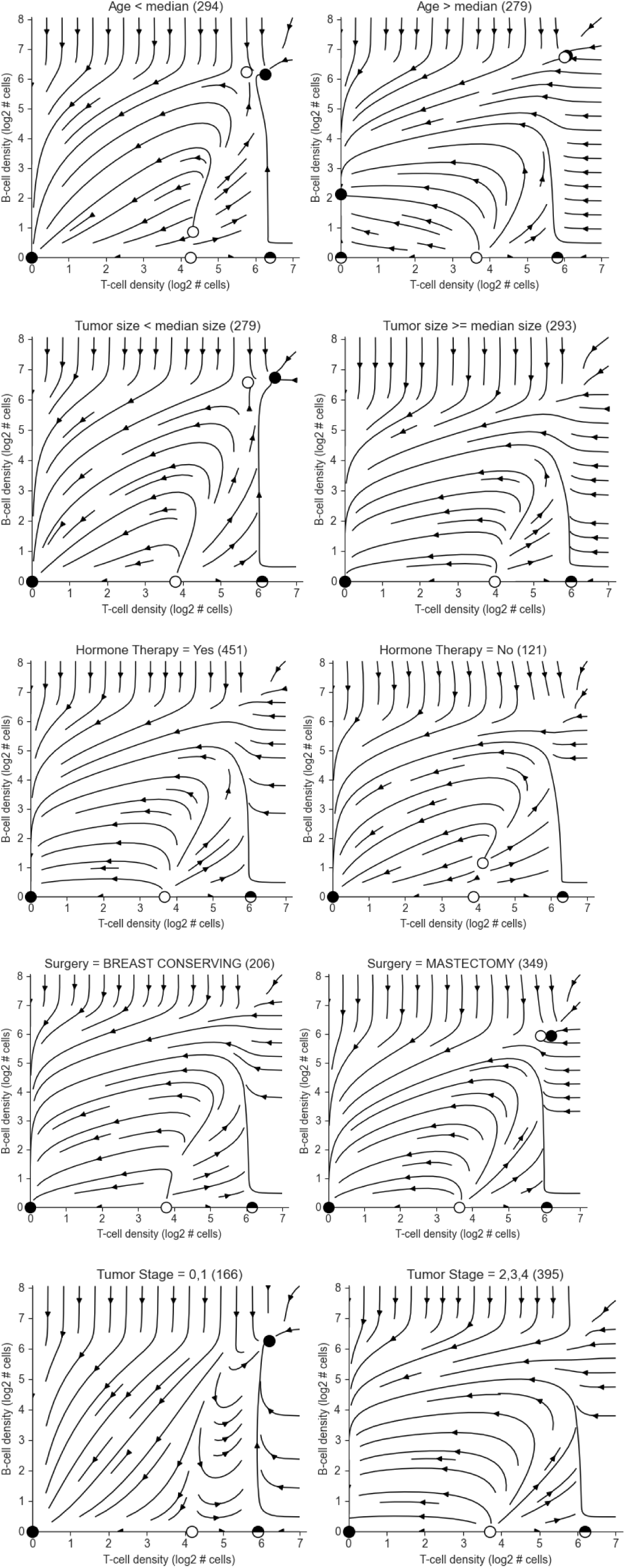
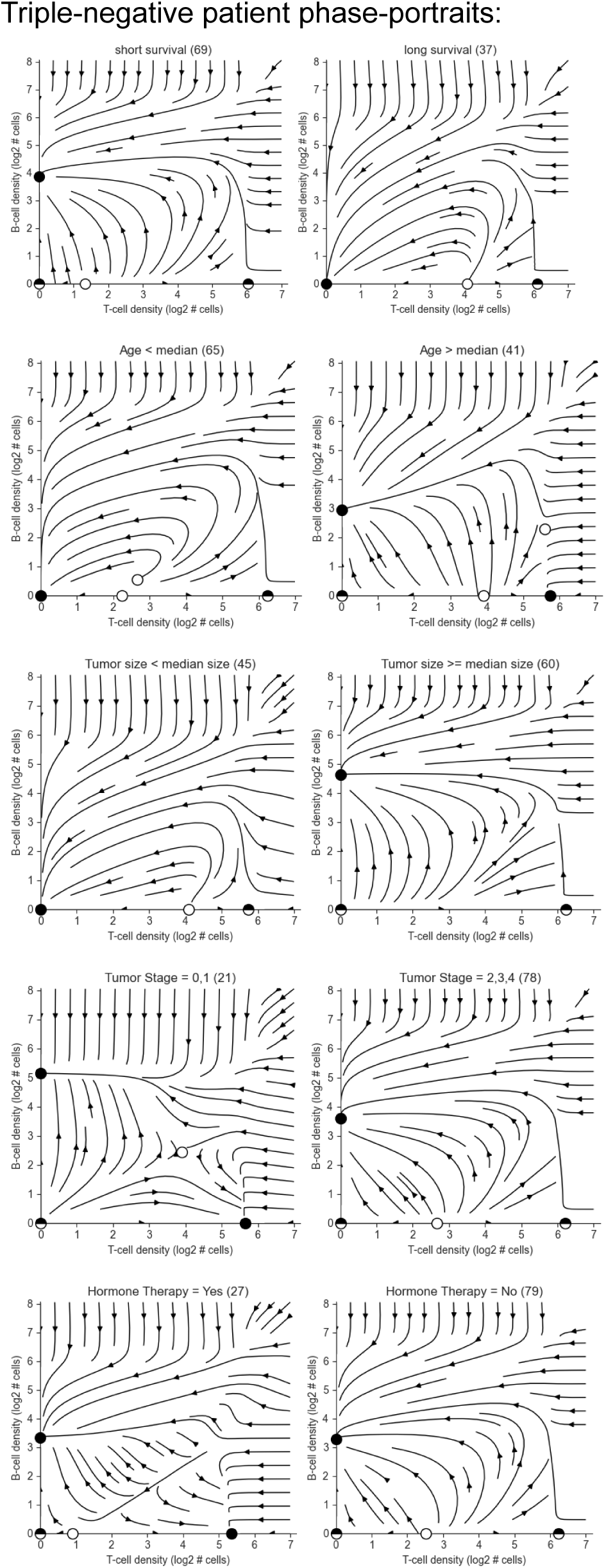
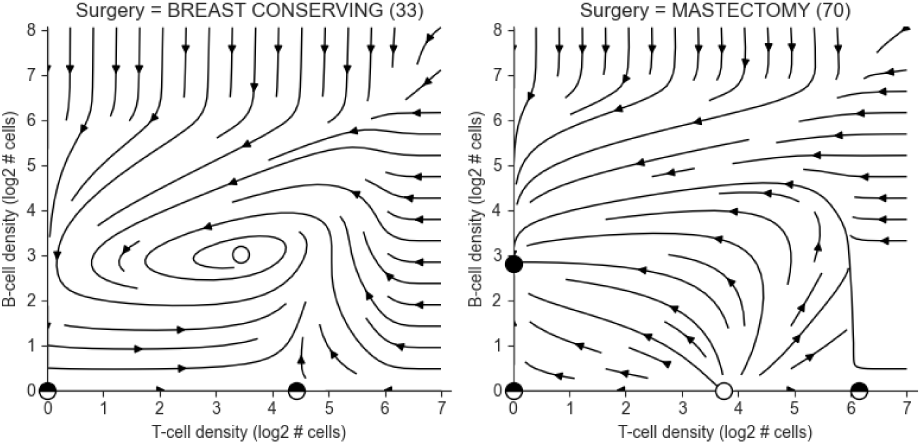

### S4J - T- and B- cell dynamics across breast cancer subtypes

Fitting separate models to breast cancer subtypes in the Danenberg et. al dataset reveals similar fibroblast-macrophage dynamics.

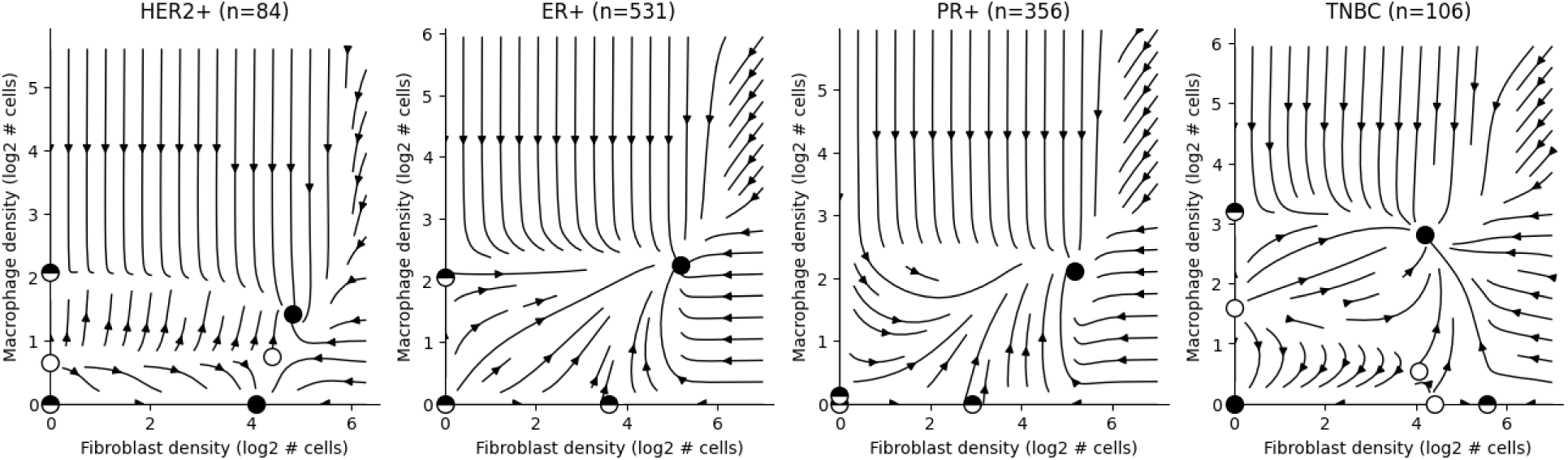

The dynamics between T- and B- cells in hormone receptor positive types displays the the immune flare discussed in the main text. In TNBC the flare doesn’t terminate with the collapse of both cells, but rather a stable B-cell fixed point.

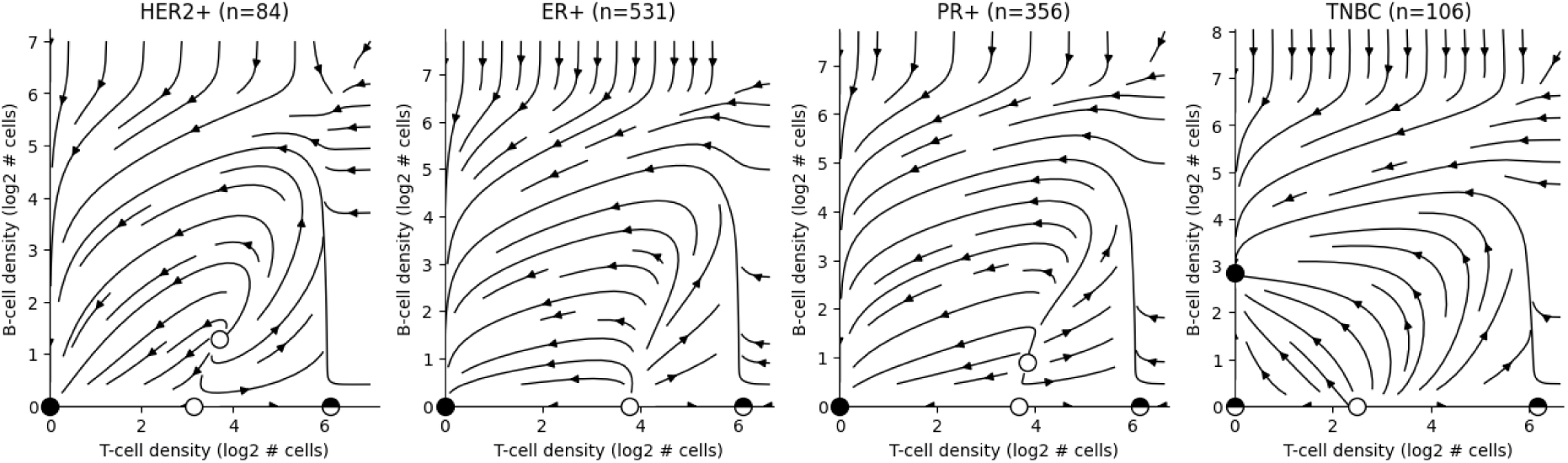

The immune flare (ending with collapse of both cells) is apparent in the mixed cohorts by Danenberg and Fischer, but the Wang cohort which consists only of triple negative breast cancer patients displays the stable B-cell fixed-point seen above.

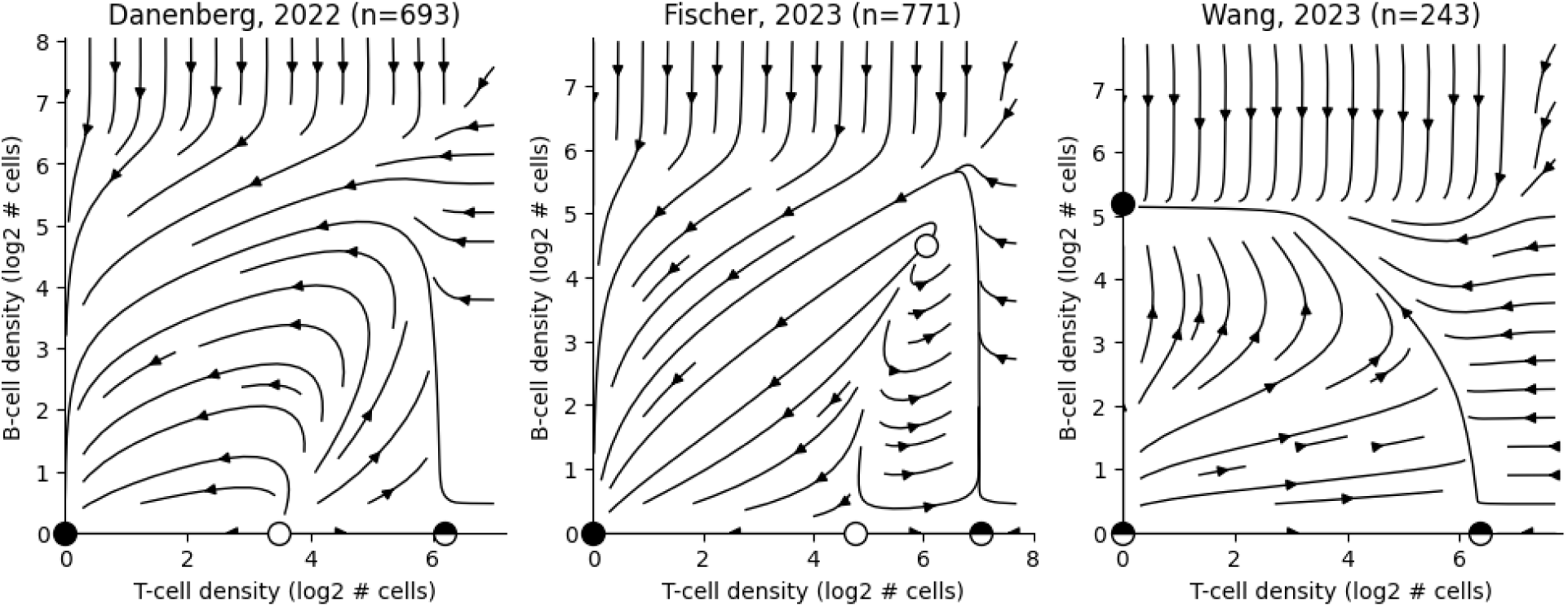

### S4K - FM dynamics are reproduced as components of a 6D model fit to all cells in the data

Model was fit without restricting the analysis to fibroblasts and macrophages with no neighboring immune cells. The model has 6-dimensions, including terms for fibroblasts, macrophages, tumor, endothelial, T- and B- cells. To plot a 2D phase-portrait, we fix the levels of other cells (here fixed to their mean value).

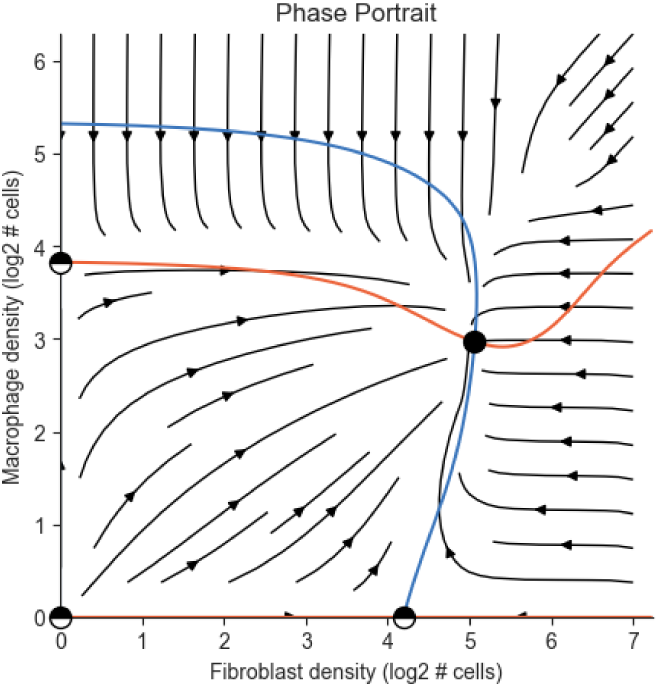

The FM dynamics within the 6D model are robust bootstrap resampling.

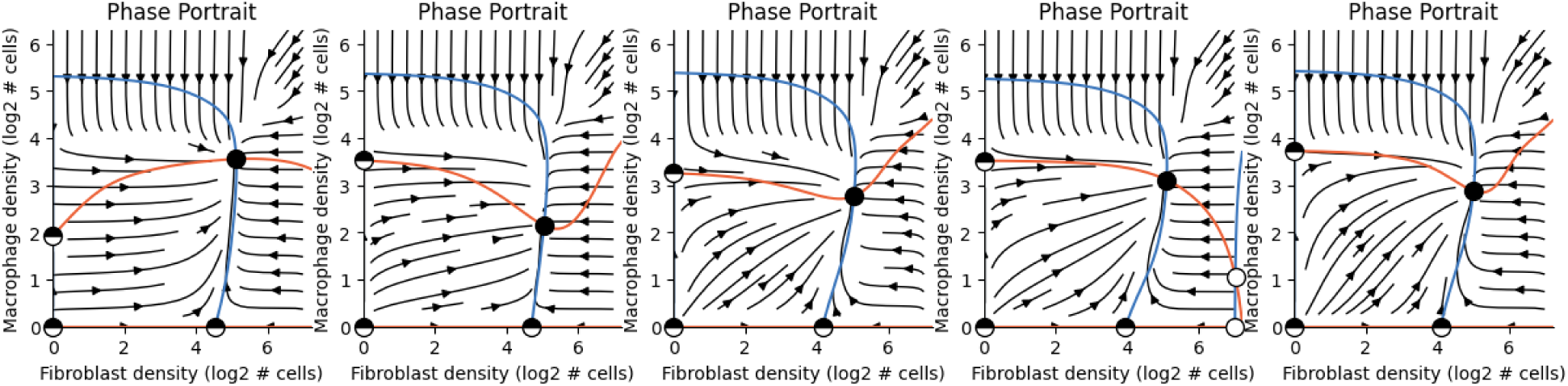

### S4L - The T- and B-cell phase-portrait is reproduced as a component of a full 6D model with fibroblasts, macrophages, tumor cells, endothelial cells, T- and B-cells

Phase-portrait is obtained by fixing other cells to their mean density.

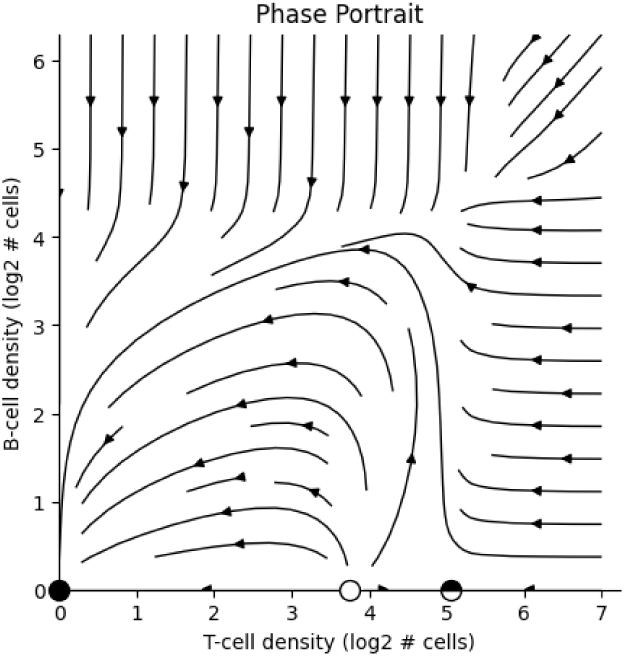

The TB dynamics under the 6D model are stable under bootstrap resampling

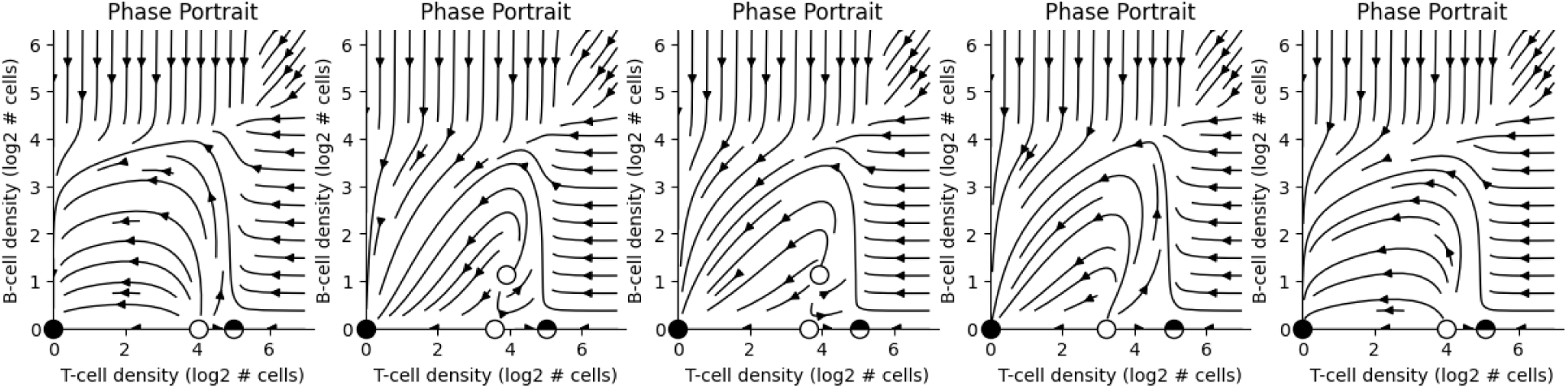

### S4M - Estimating magnitude of error due to unmodeled terms: Empirical position of tissues in state-space

Excitable dynamics drive tissues that cross the threshold density of T-cells into high T- and B-cell states, followed by a collapse of both cell types. We expect the missing component to capture migration - an external source of lymphocytes into the tissue from circulation.

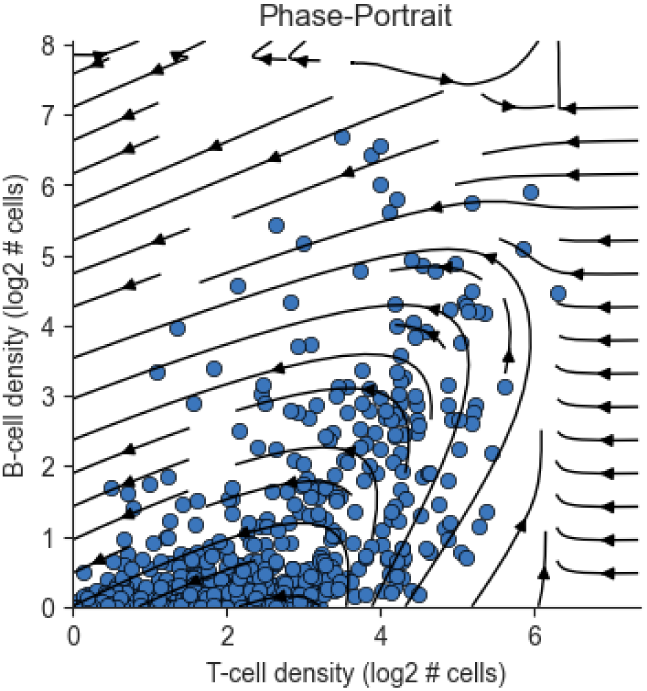

### S4N - Estimating magnitude of error due to unmodeled terms: Estimated density of tissues across the state-space

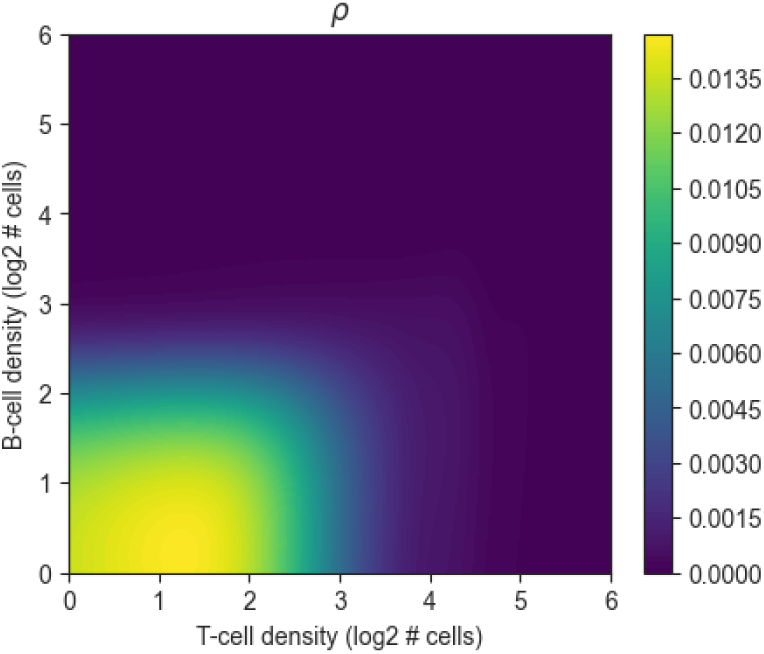

### S4O - Estimating magnitude of error due to unmodeled terms: Estimated divergence of the error field (without diffusion)

The error term has a source at (0,0) and a sink near the threshold required for an immune flare. There is near-zero influx over the rest of the state-space. The error agrees with a migration of T- and B-cells into the tissue, driving tissues with no T- or B- cells towards the region of the initial threshold of an immune flare. Thus, the high density of immune cells observed at the peak of a flare is primarily driven by local proliferation of adaptive immune cells.

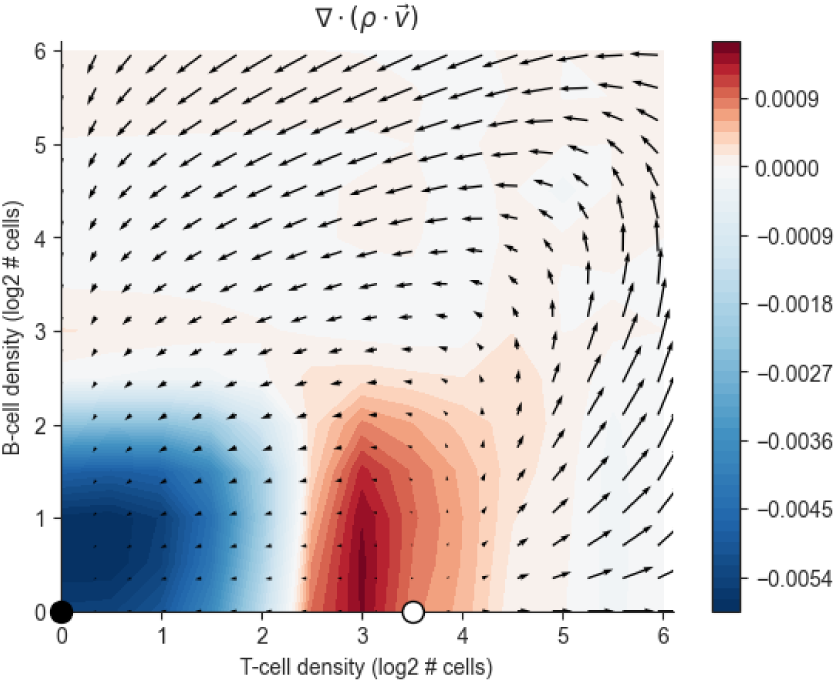

### S4P - Estimating magnitude of error due to unmodeled terms: Variance of error explained by diffusion

Diffusion doesn’t explain the error (negative coefficient):

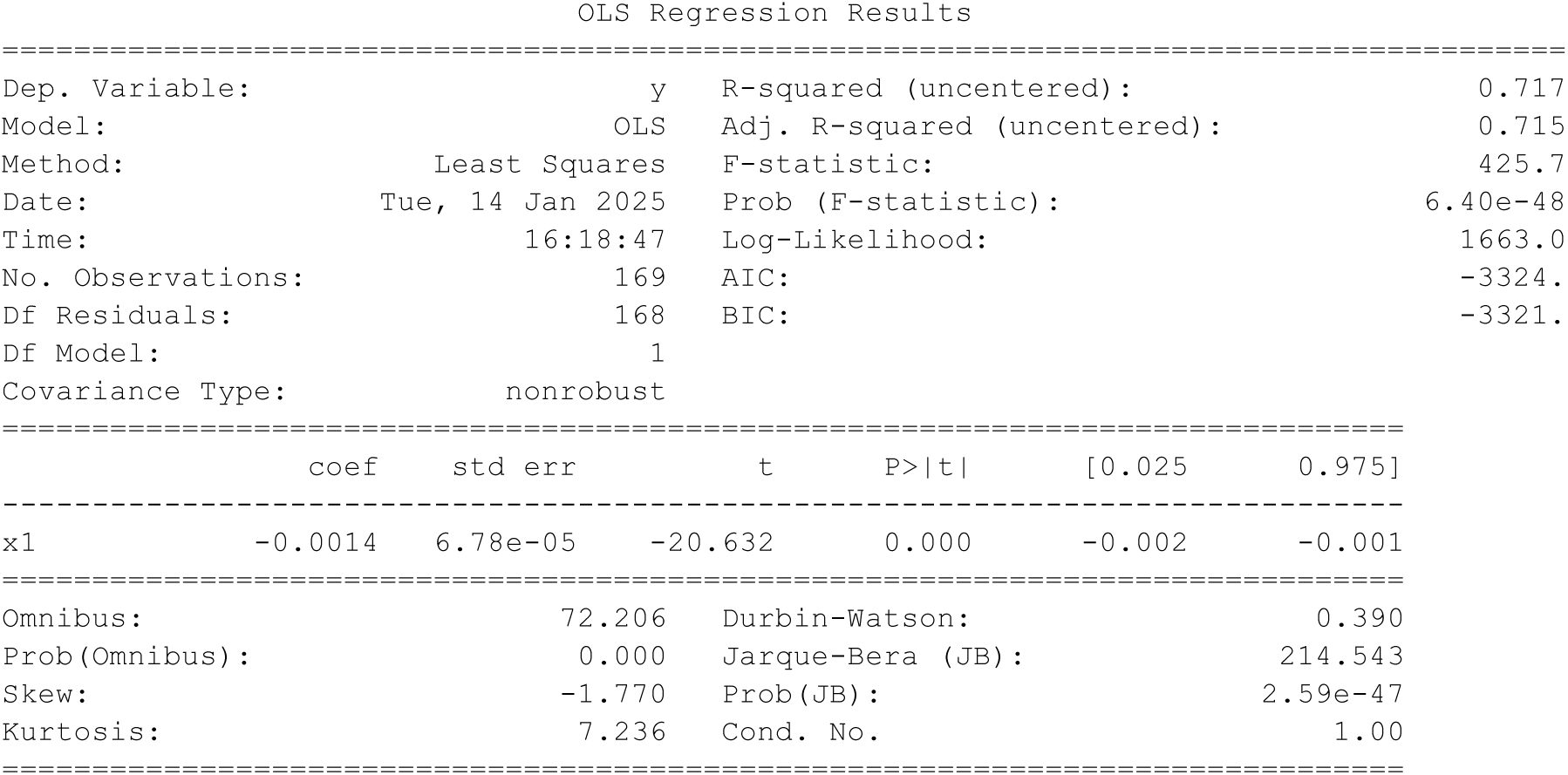

## Supplementary 5 - OSDR predicts response to chemotherapy and immunotherapy

### S5A - Prediction is robust to patient bootstrap resampling

Here we establish the robustness of the trajectories using bootstrap resampling. We randomly sample a subset of responders and non-responders, and fit a model to each group. We then apply each of the models to all early-treatment biopsies. This controls for initial tumor burden and TME composition, isolating the effect of the interactions between cells over time (“dynamics”). Plots display the distribution of final tumor density over 50 bootstrap samples.

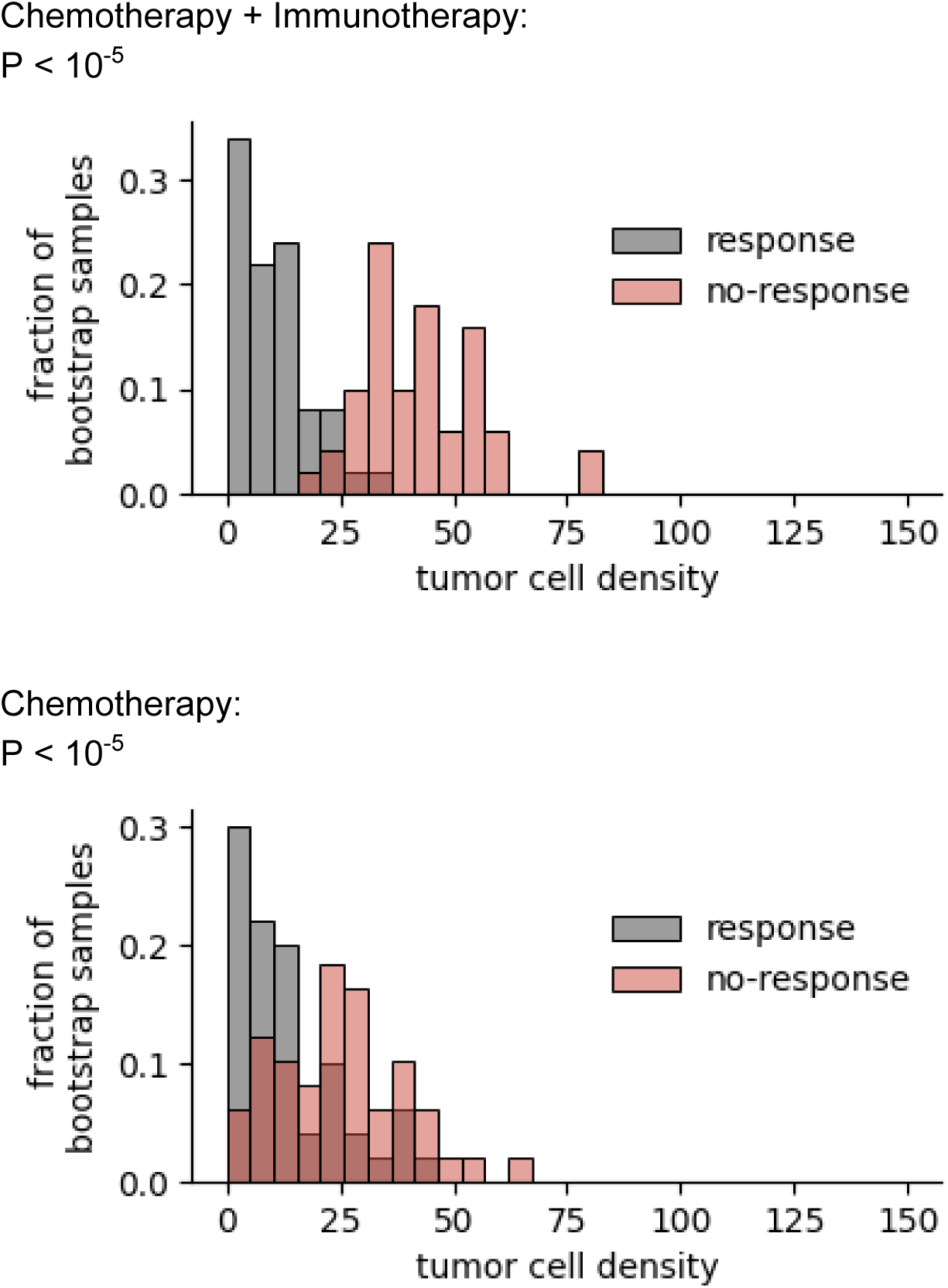

One-sided (response tumor density < no-response tumor density) Mann-Whitney U-tests were applied to compare responder and non-responder distributions.

### S5B - Early-treatment tumor cell average division rates are similar between responder and non-responder treatment groups

Fig 5b shows the predicted collapse of tumor cells in responders but not in non-responders. Here we show that this separation isn’t simply a result of a difference in the mean division rate of tumor cells between groups. That is, it is not the case that the responder model simply predicts a lower proliferation for all tumor cells. We fit a separate model to responders and non-responders and plot the resulting rates for each neighborhood in the early treatment biopsies (from all patients). Distributions largely overlap, and division rate is even higher for responders in the immunotherapy arm. This result suggests that when applying OSDR to all initial tissue compositions, the collapse of the tumor population is not due to a “crude” difference in tumor proliferation under both models.

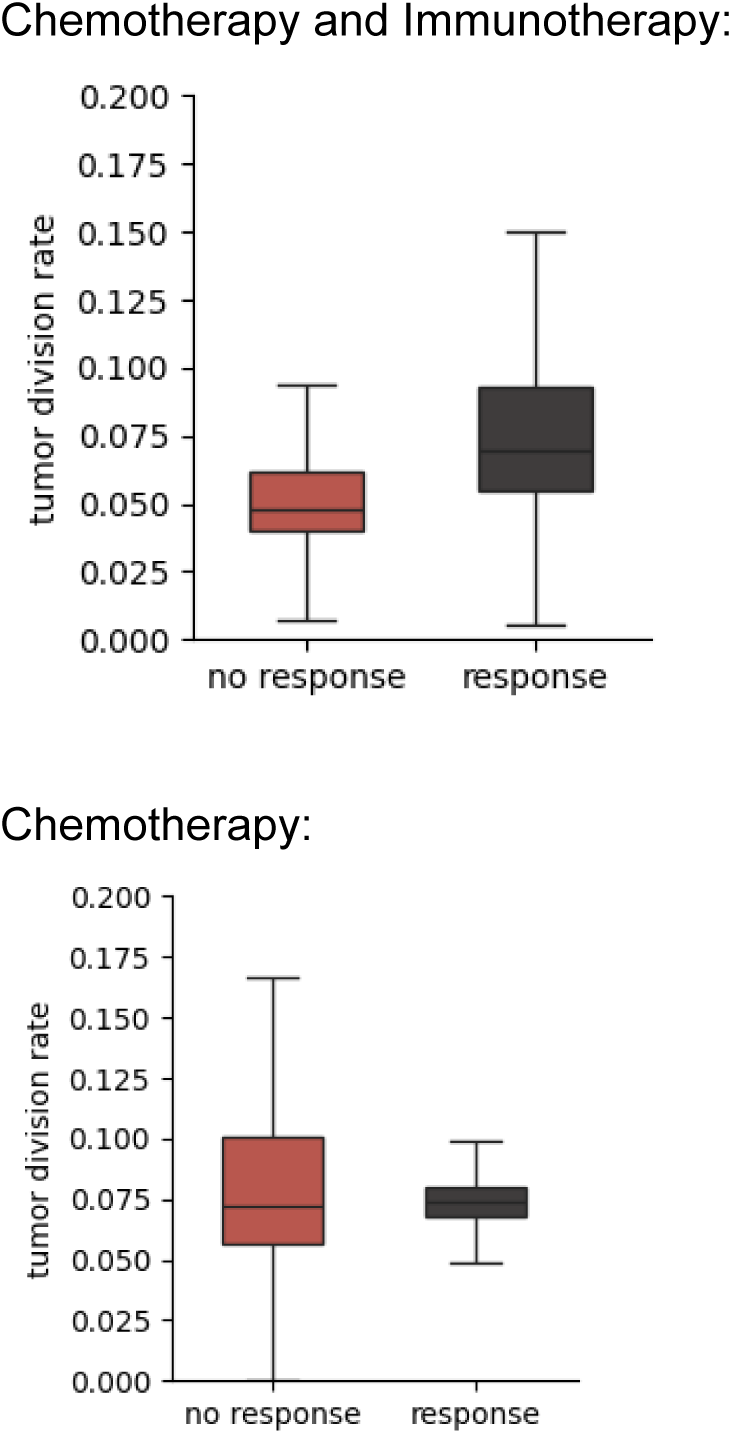

### S5C - division rates change markedly over the course of treatment

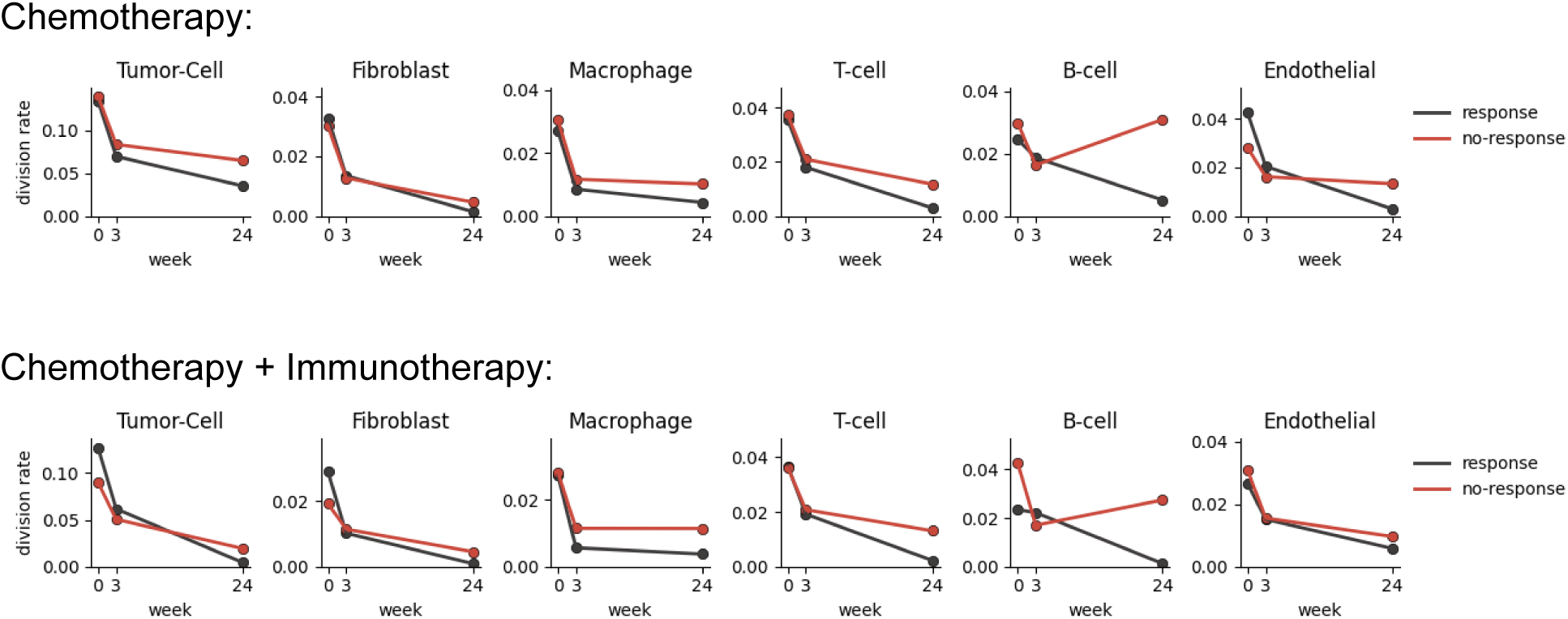

In principle, the division rate can drop over time as a result of changes in the TME, namely a different distribution of neighborhood compositions. For example, fibroblast division rate could drop because their density increased towards a stable fixed point, and not due an intrinsic change in their proliferation at the same neighborhood compositions. To control for changes in the TME we plot the distributions of cell proliferation rates over the same distribution of neighborhood states (all early-treatment states). Each row corresponds to one time point. Top row: baseline, Middle row: early treatment (3 weeks), Bottom row: post-treatment (24 weeks). Gray: response, red: no-response.

These plots suggest that each cell’s proliferation rate, over the same states, changes considerably over the half-year course of treatment. Notably, division rates drop for all cells. Possibly a cumulative effect of the chemotherapy.

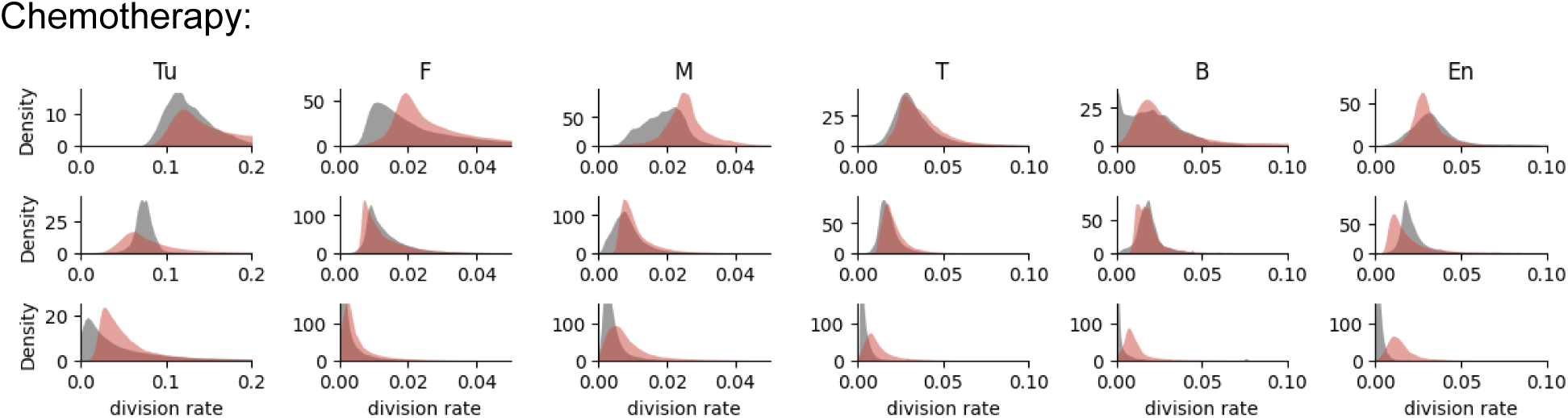

Chemotherapy + Immunotherapy:

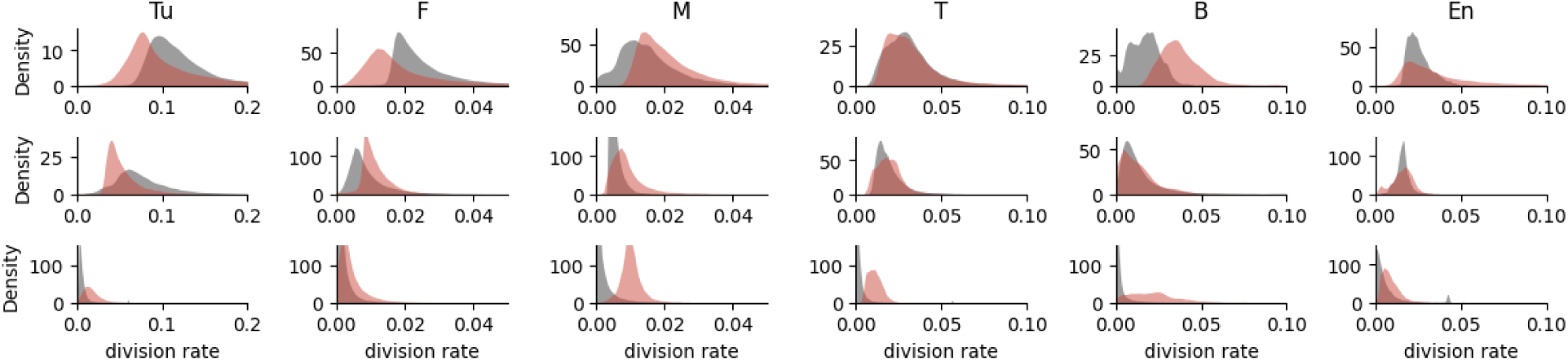

### S5D - Within and between patient variability are on similar scales

Some patients had multiple biopsies. For each patient we plotted the average density of each cell type in each of the biopsies, i.e number of cells per tissue, scaled to a fixed area of a neighborhood (top row). Error bars correspond with minimal and maximal densities over the patient’s samples.

The bottom row shows the distribution of cell densities across patients. Comparing the scale of within and between patient variance shows that most patients have biopsies from widely different locations in the between-patient distribution. Some patients even have biopsies from the two extremes of the distribution.

This plot highlights that sampling a larger tissue section should provide a larger variance of neighborhood compositions. Possibly, enough to model dynamics across the entire state-space from a single biopsy. It also suggests that predictions of response based on a single biopsy could be sensitive to the sampled tumor region.

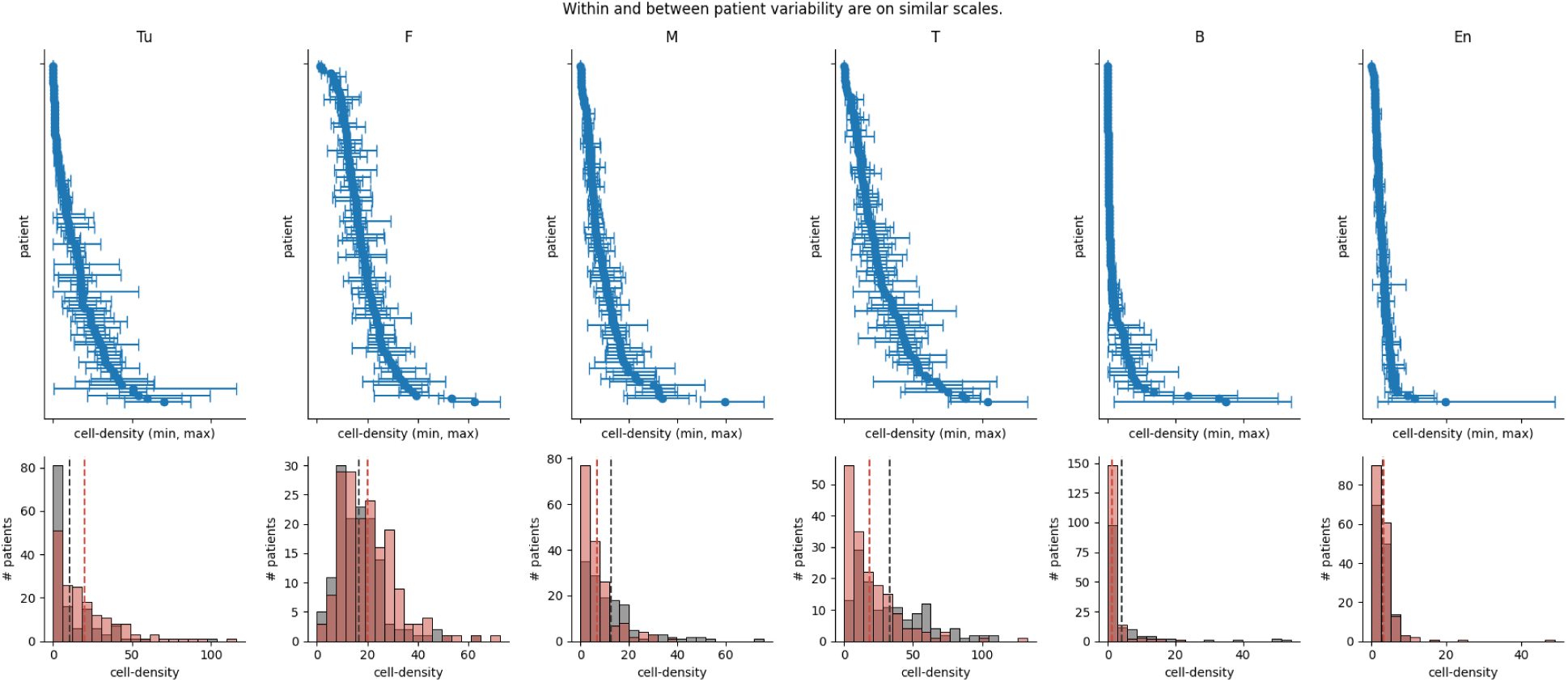

## Notes

### Competing Interest Statement

The authors have declared no competing interest.

### Summary of Updates

This revision includes analyses of two new breast-cancer cohorts as well as temporal validation of the method using longitudinal biopsies from a triple-negative breast cancer cohort.

